# Genomic impact of the second plague pandemic on three human populations

**DOI:** 10.64898/2026.06.29.730585

**Authors:** Xiaodong Liu, Kristján Moore, S. Sunna Ebenesersdóttir, Caroline Ahlström Arcini, Gordon Turner-Walker, Sean Dexter Denham, Philip Slavin, Maja Birk Søtofte, Sascha Dreyer Nielsen, Martin R. Ellegaard, Åshild J. Vågene, Ashot Margaryan, Miren Iraeta-Orbegozo, Dorothea Mylopotamitaki, Jason Laffoon, Bente Philippsen, Maria Cinthio, Rimantas Jankauskas, Justina Kozakaitė, Ethel Vol, Ariel Lomes, Shamam Waldman, Shai Carmi, Lars Alfredsson, Gianpiero L. Cavalleri, Hao Shan Chen, Olivia Cheronet, Lea Demetz, Doris Nguemo Emeruem, Daniel Fernandes, Pere Gelabert, Edmund Gilbert, Thomas F. Hansen, Eivind Hovig, Ingrid Kockum, Alejandro Llanos-Lizcano, Victoria Oberreiter, Tomas Olsson, Ron Pinhasi, Elisa Praxmarer, Birgitte Skar, Thomas Werge, James F Wilson, Hans K. Stenøien, Hannes Schroeder, Michael D. Martin, Axel Christophersen, Kári Stefánsson, M Thomas P Gilbert, Shyam Gopalakrishnan, Agnar Helgason, Ida Moltke

**Affiliations:** Section for Computational and RNA Biology, Department of Biology, University of Copenhagen, Denmark; Amgen deCODE Genetics, Iceland; Department of Anthropology, University of Iceland, Iceland; National Museum of Iceland, Suðurgata 41, 101 Reykjavík, Iceland; The Archaeologists, National Historical Museums, Odlarevägen 5, SE-226 60 Lund, Sweden; Department of Archaeology and Anthropology, National Museum of Natural Science, Taiwan; Department of Cultural Heritage, Museum of Archaeology, University of Stavanger, 4036 Stavanger, Norway; Department of History, Heritage and Politics, University of Stirling, UK; The Globe Institute, University of Copenhagen, Denmark; Department of Natural History, NTNU University Museum, Norwegian University of Science and Technology (NTNU), Trondheim, Norway; Department of Archaeological Science, Leiden University, Netherlands; National Laboratory for Age Determination, NTNU University Museum, Norwegian University of Science and Technology (NTNU), Sem Sælands vei 5, 7034 Trondheim, Norway; Kulturen’s Museums, Tegnersplatsen 6, 22350 Lund, Sweden; Center for Human Bioarchaeology and Paleogenetics, Translational Health Research Institute, Faculty of Medicine, Vilnius University, Lithuania; Braun School of Public Health and Community Medicine, The Hebrew University of Jerusalem, Jerusalem, Israel; Department of Human Evolutionary Biology, Harvard University, Cambridge, MA, USA; Institute of Environmental Medicine, Karolinska Institutet, Sweden; School of Pharmacy and Biomolecular Sciences, RCSI, Dublin, Ireland; Research Ireland FutureNeuro Centre, RCSI, Dublin, Ireland; Department of Evolutionary Anthropology, University of Vienna, Vienna, Austria; CIAS, Department of Life Sciences, University of Coimbra, Coimbra, Portugal; Human Evolution and Archaeological Sciences (HEAS), University of Vienna, Vienna, Austria; Health Technology, Danish Technical University, DTU; Neurogenomic, Translational Research Center, Danish Headache Center, Copenhagen University Hospital, 2600 Glostrup, Denmark; Department of Tumor Biology, Institute for Cancer Research, Oslo University Hospital, Oslo, Norway; Department of Informatics, University of Oslo, Oslo, Norway; Center for Molecular Medicine, Department of Clinical Neuroscience, Neuroimmunology Unit, Karolinska Institutet, Stockholm, Sweden; Department of Archaeology and Cultural History, NTNU University Museum, Norwegian University of Science and Technology (NTNU), Erling Skakkes Gate 47b, 7012 Trondheim, Norway; The Lundbeck Foundation Initiative for Integrative Psychiatric Research, iPSYCH, Aarhus, Denmark; Institute of Biological Psychiatry, MHC Sct. Hans, Mental Health Services Copenhagen, Roskilde, Denmark; Department of Clinical Medicine, University of Copenhagen, Copenhagen, Denmark; Usher Institute of Population Health Sciences and Informatics, University of Edinburgh, Edinburgh, UK; Faculty of Medicine, University of Iceland, Reykjavík, Iceland; Department of Anthropology, University of Iceland, Reykjavík, Iceland

**Author notes:** These authors contributed equally to this work (Xiaodong Liu, Kristján Moore, S. Sunna Ebenesersdóttir). Correspondence should be addressed to S.G., A.H. and I.M.

## Abstract

The second plague pandemic (early 14th – early 19th centuries), caused by *Yersinia pestis,* had a profound demographic, socio-economic and cultural impact across Eurasia and North Africa. Many regions in Europe and the Middle East are estimated to have lost 40–60% of their human populations, with some areas suffering even higher mortality. Whether exposure to *Y. pestis* drove strong positive selection on protective genetic variants in the human genome, and how the pandemic shaped migration patterns, remains debated despite several recent studies based on ancient DNA. Here, we analyse a markedly larger, higher-coverage, and geographically diverse whole-genome sequencing dataset from 529 ancient individuals (mean depth 8.8x) dating to either before or after the arrival of the pandemic at three sites in northern Europe: Trondheim (Norway), Lund (Sweden) and Vilnius (Lithuania). Genome-wide scans for selection provide no evidence for strong positive selection acting on specific genetic variants driven by *Y. pestis* exposure: we neither replicate selection signatures reported by previous studies nor identify credible novel genome-wide significant candidates. However, at all three sites, we observe a reduction in long-range immigration, indicated by a drop in the diversity of ancestry that followed the arrival of *Y. pestis* and broadly coincided with the end of the Viking Age, Christianisation and the onset of the Little Ice Age. Our results shed important light on the demographic impact of major sociohistorical changes that occurred during the late Medieval period in Scandinavia and the Baltic region and link Christianisation to increased diversity in ancestry before the pandemic.

## Introduction

The Black Death (1347-1353) and the subsequent waves of the second plague pandemic rank among the deadliest events in human history, yet their impact on the gene pools of affected populations is a subject of ongoing debate. After entering Europe in 1347 and claiming some 40–60 percent of its population^1–3^, the causative agent, *Y. pestis*, circulated in the continent for over 450 years, both in domestic reservoirs and via reintroductions, causing repeated waves of mass mortality. Historical evidence indicates that this pandemic had a substantial socio-economic impact across Eurasia and North Africa^4^. For instance, it is thought to have contributed to the decline of feudalism and serfdom in England during the 14th and 15th centuries^5^, and to the emergence of serfdom in eastern Europe on a longer time-scale^6,7^. By dramatically reducing population size, the pandemic also altered the land-to-labour ratios across affected regions, increasing per capita access to land^8^.

The exceptionally high mortality attributed to *Y. pestis*, particularly during the initial waves of the pandemic^3,9,10^, created the conditions for intense selective pressure on human populations, favouring genetic variants that conferred even partial protection against infection or mitigated disease severity. Some early studies of present-day populations proposed that a 32-base pair deletion in the *CCR5* gene was subject to positive selection due to plague mortality, with an estimated allele age of only 700 years and an allele frequency of around 10-15% in present-day northern Europeans^11^. However, subsequent studies cast doubt on this result, based both on simulations and questions about the potential biological mechanism^12–14^. More recently, analyses of present-day populations led to hypotheses of plague-driven selection on the Toll-like receptor (*TLR*) 1, 6 and 10 genes^15^. Importantly, it has now become possible to directly assess changes in the frequency of variants driven by selection or other evolutionary forces, by sequencing the genomes of individuals who died before, during and after the arrival of the Black Death. Ancient DNA studies based on targeted capture of sequences from immune genes^16,17^ reported evidence for plague-driven selection acting on several loci: *ERAP2*, *TICAM2*, *NFATC1,* and *CTLA4*. However, whole-genome studies^18–20^ did not find compelling evidence for selection at those or other loci, aside from a suggestive signal in *RIPK2* (Hui et al. 2024). Moreover, the findings of Klunk et al. (2022) have been challenged on methodological grounds^21^ and sample sizes in that and other previous studies have been limited. Thus, it remains unclear whether the second plague pandemic altered the frequency of genetic variants due to protective effects against infection or mortality.

To better understand the impact of the pandemic, we analysed a large dataset of 529 ancient genomes from cemeteries or mass burials in three northern European countries: Lund (N=337) in Sweden and Trondheim (N=135) in Norway, which experienced mortality close to 50% during the Black Death^4^, as well as Vilnius (N=57) in Lithuania (Fig.1, Supplementary Tables 1 and 2, Supplementary Note 1) where the impact of the pandemic’s early waves was more moderate^22,23^. Each of these sites provided well-dated, stratified burial contexts, before and after the arrival of the Black Death, enabling direct temporal comparison of allele frequencies with the aim of identifying possible targets of selection.

**Figure 1:**
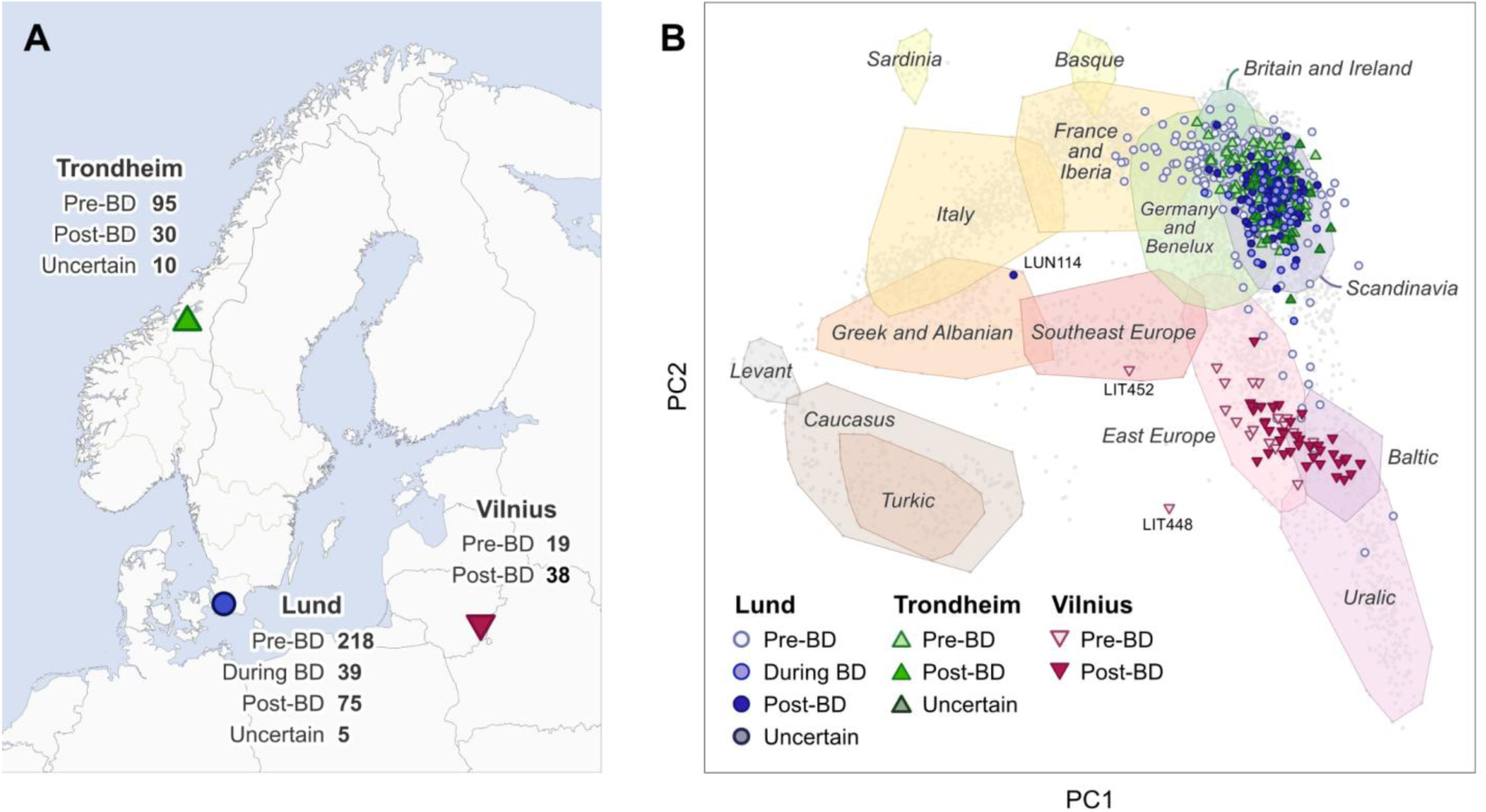
Overview of dataset. **A)** Map showing the three sites: Lund, Trondheim, and Vilnius, and the number of ancient individuals from each site by period that passed quality control. **B)** Projections of all ancient individuals onto a PCA of present-day West Eurasian populations with PC1 broadly representing north-south differentiation and PC2 east-west differentiation. Grey points represent present-day individuals used to calculate the PCA, whose coordinates are summarised by convex hulls (transparent polygons) drawn to include 95% of the labelled group (italic text). Notable outliers among the ancient individuals are labelled with non-italic text.

Another important question relates to the demographic impact of the second plague pandemic. Historical evidence indicates substantial changes to migration patterns during and after plague outbreaks in several parts of Eurasia, including England, the Low Countries and Scandinavia, with increased immigration to replenish depopulated areas in some regions and reduced long-distance mobility in others^24–27^. Ancient DNA analyses of European populations have revealed differing degrees of changes in ancestry coinciding with the arrival of the second plague pandemic. One study of 24 ancient and 30 present-day genomes indicated a shift in ancestry following the arrival of the Black Death in Trondheim in 1349^18^. In contrast, several other studies reported no change in ancestry attributable to the Black Death or subsequent waves of the pandemic^17,19,28^. However, these previous studies of the genetic consequences of either the Black Death specifically, or the second plague pandemic more generally, are limited either by small sample sizes^17–19^, single sampling locations^17–19^, or targeting only a small subset of the genome (mtDNA sequences from 264 individuals from London and across Denmark^28^).

Using the 529 ancient genomes recovered from the cemeteries in Lund, Trondheim and Vilnius, we evaluated evidence for temporal changes in the composition of ancestry, assigned at the level of countries and regions within Scandinavia, before and after the arrival of the Black Death. The arrival of the Black Death broadly coincided with the onset of the Little Ice Age^29,30^ and two centuries earlier was the end of the Viking Age, both potential drivers of demographic and social and economic changes. No less important was the introduction of Christianity, which occurred during the 10th–12th centuries in Scandinavia and was seeded primarily by missionaries and emissaries from Germany and the British Isles^31–33^. This occurred even later in the Baltic region, during the 12th–14th centuries, where Orthodox Christianity was introduced from the east^34^. These factors are known to have had a far-reaching impact on the socioeconomic and demographic histories of Lund, Trondheim and Vilnius, including population size and long-range migration^35^. We used the temporal stratification of the ancient genomes from these sites to assess whether this tumultuous period was associated with detectable changes in ancestry.

Remarkably, our results provide no evidence of genetic variants that affected the reproductive success of carriers during the second plague pandemic, leading to statistically significant changes in allele frequencies. However, they reveal marked changes in the profiles of ancestry observed in the cemeteries of Lund, Trondheim and Vilnius, indicating a shift towards more insular and less cosmopolitan societies. While plague may have contributed to these observed changes, other concurrent historical processes such as Christianisation were likely important factors.

## Results

### Description of dataset

The initial dataset contained 575 ancient genomes from the three sites: 353 from Lund, 164 from Trondheim (24 published previously in Gopalakrishnan et al. 2022^18^), and 58 from Vilnius. Following quality control, 38 genomes were excluded due to low sequencing depth (depth of coverage <0.1x, Supplementary Figs. 1 and 2) and eight more were removed based on evidence of contamination, leaving 529 high-quality genomes for downstream analyses (Supplementary Table 1). For each of the three sites, individuals were assigned as dating to before or after the arrival of the Black Death in 1349/50, hereafter referred to as pre-BD (N=332) and post-BD (N=143) individuals (Fig. 1A). Temporal groups could not be confidently assigned to 15 individuals based on archaeological context and 39 individuals from Lund were assigned as during-BD. These latter individuals were from an assemblage previously linked to the Black Death and two subsequent plague outbreaks, tentatively dated to the late 14th century and characterised by increased burial activity and the occurrence of double and triple graves^36^. The average depth of coverage for the 529 ancient genomes was 8.8x (median: 10.3x; range: 0.1–39.0x; Supplementary Fig. 2). Genotypes were imputed for each of the ancient individuals using GLIMPSE2^37^ based on a whole genome sequencing reference panel of 50,129 contemporary individuals of predominantly European ancestry^38^. The resulting dataset comprises genotypes for 99.6 million loci. Identity-by-descent (IBD) kinship analysis identified relatively few close relatives within these urban sites: three first-degree, one second-degree and three third-degree relative pairs in Lund (all pairs within the same archeological layer or across adjacent layers); one first-degree pair in Trondheim (both pre-BD in nearby burial locations); and two first-degree and one third-degree pair in Vilnius (all pairs within the same burial locations) (Supplementary Fig. 3, Supplementary Table 3). These were all included in the final dataset. We observed no close relatives between sites and no instances of elevated inbreeding (*F_ROH_*>0.03) (Supplementary Table 4).

To examine the broad-scale ancestry of the ancient individuals, we projected them onto a principal components analysis (PCA) of 9,052 present-day individuals from 67 West Eurasian populations^35^. Figure 1B shows that the first two principal components (PCs) broadly reproduce the geographic structure of Europe, with PC1 reflecting north-south differentiation and PC2 east-west. As expected, most ancient individuals from Lund, Trondheim and Vilnius project close to present-day individuals from the same regions, consistent with continuity in ancestry. However, a substantial number of individuals fall outside these local clusters, indicating that the burial sites sampled from these populations were not homogeneous in terms of ancestry.

### No evidence for large allele frequency changes due to plague-driven selection

To investigate whether *Y. pestis* drove selection on specific genetic variants, we performed a genome-wide association analysis (GWAS) within each of the three cohorts, testing for differences in allele frequencies between the pre- and post-BD groups. Analyses were restricted to approximately 5.6 million common variants with minor allele frequency (MAF) >0.05, calculated separately within each GWAS dataset, that passed stringent quality-control filters (see Methods). Given the heterogeneity of ancestry observed among ancient individuals from each site (Fig. 1B) and the presence of a few close relatives, we used a linear mixed model to account for population structure and relatedness. In the model, we included the first seven PCs from the West Eurasian PCA (Supplementary Fig. 4) and sex as covariates. Q-Q plots showed limited signs of inflation of the test statistics for Lund (λ=1.040, Supplementary Fig. 5) and Trondheim (λ=1.075, Supplementary Fig. 6), indicating that the effects of population structure and relatedness were adequately controlled. In contrast, the test statistics were substantially more inflated for Vilnius (λ=1.211, Supplementary Fig. 7A), presumably due to small sample size (Supplementary Fig. 7B), and the results for Vilnius were therefore excluded from further selection analyses. To increase statistical power, we combined the association results for Lund and Trondheim in a meta-analysis, using the union of loci from the two sites (Fig. 2A left, Supplementary Fig. 8) under the assumption that the genetic architecture of susceptibility to and/or protection from *Y. pestis* would be similar in the two populations. Neither the separate analysis of each site nor the meta-analysis revealed any genetic variants below the functionally informed genome-wide significance thresholds proposed by Sveinbjornsson et al.^39^.

**Figure 2:**
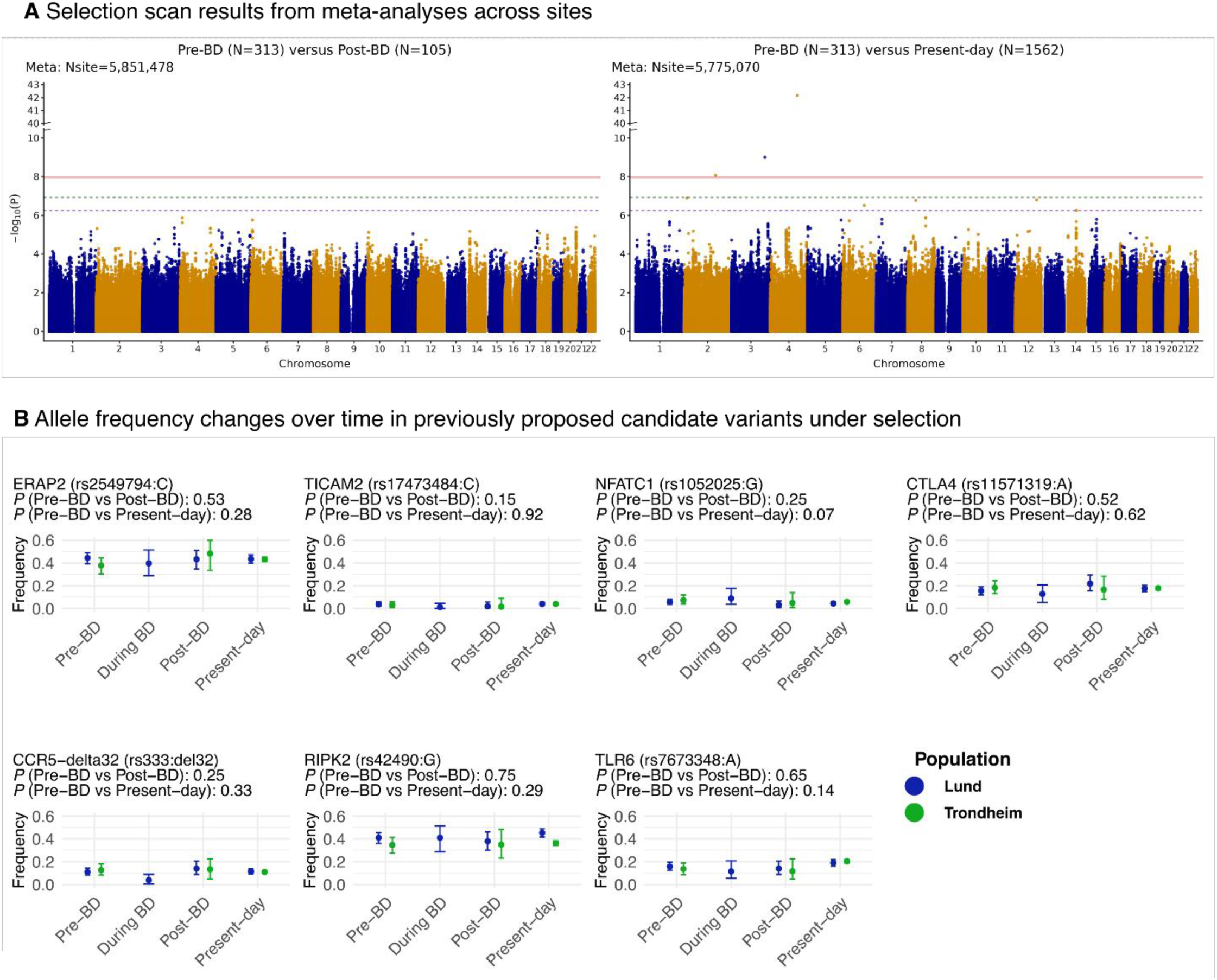
Results of selection analysis. **A)** Manhattan plots of temporal GWAS meta-analyses comparing pre-BD and post-BD ancient individuals from Lund and Trondheim (left) and pre-BD ancient individuals versus present-day proxies across sites (right). Present-day proxy populations comprise individuals from Denmark and Skåne (Sweden) for Lund, and individuals from Norway for Trondheim. Red, green and purple horizontal lines indicate genome-wide significance thresholds for low-impact (1.1×10^-8^), moderate-impact (1.2×10^-7^) and loss-of-function variants (5.8×10^-7^), respectively, as proposed in ^39^. The marker set for each meta-analysis comprised the union of markers included in the corresponding single-site GWAS: 5,637,038 for Lund and 5,633,253 for Trondheim in the pre-BD versus post-BD comparison, and 5,652,057 for Lund and 5,655,420 for Trondheim in the pre-BD versus present-day comparison. **B)** Temporal allele frequency trajectories for variants previously reported to be under selection associated with the Black Death, including loci reported by Klunk et al.^16^ (*ERAP2*, *TICAM2*, *NFATC1, CTLA4*), Stephens et al.^11^ (*CCR5*-delta32), Laayouni et al.^15^ (*TLR* gene cluster), and Hui et al.^19^ (*RIPK2*). For the *TLR1*, *TLR6* and *TLR10* gene cluster, only the variant with the lowest *P*-value in the meta-analysis of pre-BD vs present-day is shown. Frequencies are shown for available time periods: pre-BD, during-BD, post-BD and present day. The during-BD time point is available only for Lund. In each panel, the plotted frequency corresponds to the allele indicated in the panel label, shown as gene (rsID: allele). Error bars indicate 95% confidence intervals.

To further identify genetic variants with significant temporal changes in frequency, we performed GWAS comparing pre-BD individuals from Lund to present-day individuals from Denmark and Skåne, which is the region of Sweden that Lund is located in (increasing from N_post_=75 to N_present-day_=380) and pre-BD individuals from Trondheim to present-day individuals from Norway (increasing from N_post_=30 to N_present-day_=1182) (Supplementary Figs. 9 and 10). Since sequencing data are only available for very few present-day Baltic individuals, we could not perform a similar analysis for Vilnius. Hence, as before, we performed a meta-analysis of Lund and Trondheim only (Fig. 2A bottom panel, Supplementary Fig. 11), and observed very little inflation of test statistics (Lund λ=1.011, Trondheim λ=1.018, meta-analysis λ=1.022, Supplementary Figs. 9-11, Fig. 2A right). In the meta-analysis, we found three genome-wide significant signals on chromosomes 2, 3 and 4 (Fig. 2A right, Supplementary Fig. 11, Supplementary Table 6a). However, further inspection suggested that it is very unlikely that any of these reflects allele frequency changes caused by selection associated with the Black Death, as all three signals showed evidence of poor aDNA mappability and consisted of only a single variant each (Supplementary Fig. 12, Supplementary Note 2, Supplementary Table 6b). We also identified two genome-wide significant signals in population-specific analyses, both of which were observed in Lund. Of these, only one warrants a brief description (Supplementary Note 2). This signal consists of several genome-wide significant variants within the T-cell receptor alpha (*TCRA)* gene segments in the q11.2 region on chromosome 14 (lead SNP rs71418263, chr14:22193177, *P*=3.02×10^-12^ in Lund with frequency estimates of 0.09 in pre-BD individuals (95% CI: 0.06‒0.12), 0.15 in post-BD individuals (95% CI: 0.09‒0.21), and 0.29 in present-day individuals (95% CI: 0.26‒0.32); Supplementary Table 6A, Supplementary Fig. 12). However, we caution that the TCRA region is highly polymorphic and structurally complex, with many similar duplicated gene segments making it difficult to map and genotype confidently^40^. We therefore do not consider it a strong candidate locus, despite its role in immune response to pathogens. For a list of top 500 variants in each GWAS, see Supplementary Table 7.

Finally, we examined the evidence in Lund and Trondheim for changes in the frequencies of the genetic variants reported by Klunk et al. as selection targets against *Y. pestis* in the London and Danish cohorts^16^ and other previous papers^15,19^. None of these variants exhibited significant allele frequency changes based on the meta-analyses, even when using a lenient replication *P* value threshold of 0.05 (Fig. 2B, Supplementary Table 8). The only variant that comes close to replication is rs42490 from *RIPK2*^19^ with *P*=0.04 in Lund pre-BD vs. present-day. However, the direction of allele frequency change is opposite to that reported by Hui et al. and neither the meta-analysis nor any of the other single-site analyses is significant at *P*<0.05 for rs42490.

Our finding of no genome-wide significant differences of allele frequencies attributable to *Y. pestis* does not rule out selection driven by the second plague pandemic, but it does rule out strong selection resulting in large allele frequency changes. To shed light on the range of frequency changes that can be ruled out for the well-powered sampling sites of Lund and Trondheim, we conducted simple power analyses both per-site and with the two sites combined. For the latter, we assumed that the genetic architecture of susceptibility to *Y. pestis* infection was the same in both populations, so that the sample sizes for each time period could be summed across the two sites. This implies that these power estimates are upper limits for the power of the meta-analysis. Despite substantially larger and broader sampling than previous studies, our study design provides only modest power to detect genome-wide signals of allele frequency change when using a *P*-value threshold (α) of 1.1×10^-8^. For variants with combined MAF ≥ 0.05 across pre- and post-BD individuals, well-powered detection (power > 80%) required minimum frequency differences in the range of 0.12 and 0.26 in the meta-analysis of pre-BD versus post-BD comparison (Lund: 0.15-0.31, Trondheim: 0.22-0.49). The exact minimum difference that is detectable depends on the pre-BD allele frequency (Supplementary Fig. 13). For the pre-BD versus present-day comparison, the equivalent range was 0.07 to 0.15 (Lund: 0.08-0.20, Trondheim: 0.13-0.24). In contrast, our design was well-powered to detect frequency changes for previously reported candidate variants using a nominal replication threshold of α=0.05 (power > 80% in meta-analysis for all except *TICAM2,* where the power is above 65%, Supplementary Fig. 14, Supplementary Table 9). For instance, we expect to have over 97.6% power to detect the allele frequency changes reported for the *ERAP2* variant (rs2549794)^16^, in our meta-analysis assuming the reported allele frequencies were accurate and similar in Scandinavia (Lund: 91.5%, Trondheim: 57.8%).

### Greater diversity of ancestry before the Black Death

While most pre- and post-BD individuals from the Lund, Trondheim and Vilnius sites cluster close to present-day individuals from the same regions in the PCA of West Eurasian populations, we also observe outliers and temporal changes (Fig. 1B, Supplementary Fig. 15). A previous smaller study of Trondheim found that pre-BD individuals were more diverse in ancestry than those sampled from later periods^18^, and similar findings have been reported for other regions of Scandinavia^35^. Using the considerably larger sample sizes assembled in this study, we performed detailed ancestry analyses across the three sites to assess whether the demographic disruption caused by the arrival of the second plague pandemic and other factors such as Christianisation and the Little Ice Age, were associated with changes in ancestry.

At all three sites, we observe greater diversity of ancestry prior to the arrival of the Black Death than afterwards. Pre-BD individuals are more dispersed in the West Eurasian PCA space than post-BD individuals, as indicated by larger generalized variance across the first seven PCs and by significant differences between pre-BD and post-BD covariance matrices at each site (Box’s M test: Lund *P*<2.2×10^-16^, Trondheim *P*=1.04×10^-6^, Vilnius *P*=5.77×10-^4^; Supplementary Table 10). Notably, when focusing only on PC5, which captures differentiation between Scandinavian and British-Irish ancestry, ANOVA tests reveal a directional shift between pre-BD and post-BD individuals in both Lund (*P*=1.83×10^-7^) and Trondheim (*P*=4.99×10^-5^), concordant with a decline in British-Irish ancestry. Consistent with this pattern, supervised ADMIXTURE analysis using Norse and Gaelic reference populations reveals a reduction in ancestry from British Isles and Ireland from pre-BD to post-BD individuals in both Trondheim (*P*=1.56×10^-12^, beta-regression, Supplementary Fig. 16) and Lund (*P*=1.6×10^-3^). These findings are in line with a previous study of Medieval Trondheim (Gopalakrishnan et al. 2022).

To further test for temporal differences in the diversity of ancestry based on the PCA, we calculated the Mahalanobis distance (*D_M_*) between each ancient individual and present-day populations included in the West Eurasian PCA (Fig. 1) using the first seven PCs. For this purpose, we used individuals from Denmark (N=797), Norway (N=510) and Lithuania (N=121) as present-day proxies for Lund, Trondheim and Vilnius, respectively. The distribution of *D_M_* shows that post-BD individuals from Lund and Vilnius are closer to their corresponding present-day populations than pre-BD individuals (*P*<0.01, Wilcoxon test) (Fig. 3A top panel, Supplementary Fig. 17). While this difference is not significant for Trondheim (*P*=0.14, Wilcoxon test), it is in the same direction. Overall, PCA-based analyses indicate a reduction in the diversity of ancestry at all three sites following the arrival of the Black Death, which is particularly marked in Lund and Vilnius.

**Figure 3:**
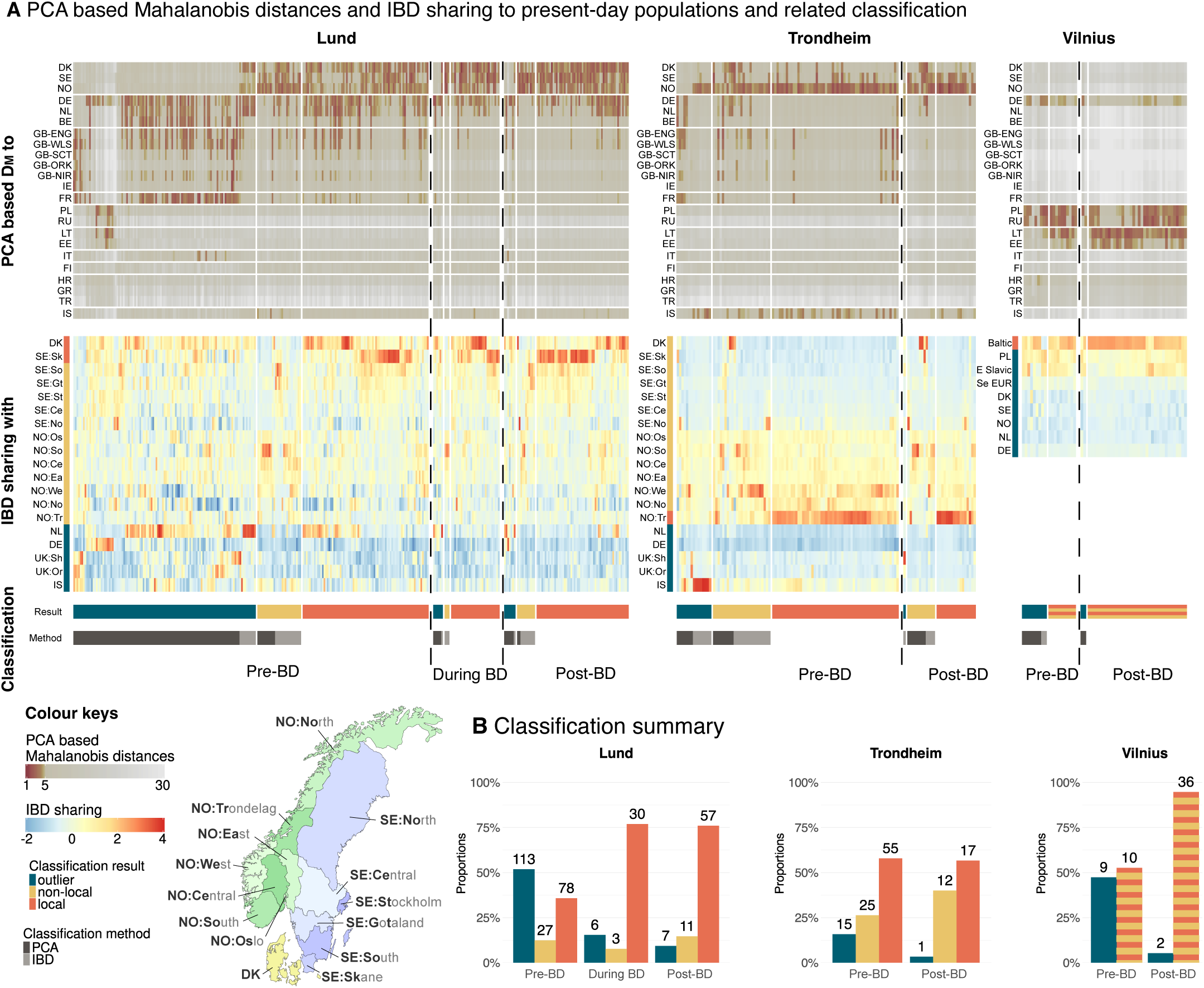
Genetic ancestry of ancient individuals. **A Top:** Heatmap of PCA-based *D_M_* from ancient individuals to present-day Eurasian populations. **Middle**: Heatmap of IBD sharing between ancient individuals and present-day populations, with values standardized within each ancient individual. On the y-axis, present-day proxies for ancient individuals are indicated with vertical red bars, populations from other Scandinavian regions are shown in yellow, and other European regions (outside Scandinavia for Lund and Trondheim, and outside the Baltic for Vilnius) are shown in blue. Row names in the heatmaps in the top and middle panels show two-letter ISO-3166 country codes. In the middle heatmap for Lund and Trondheim, labels beginning with “SE:”, “NO:”, and “UK:” denote regions within Sweden, Norway and the United Kingdom, respectively. For Sweden and Norway, the regions referred to and their abbreviations are illustrated on the map. For the UK, “Sh” and “Or” refer to Shetland and Orkney, respectively. In the middle heatmap for Vilnius, “Baltic”, “E Slavic” and “Se EUR” denote the Baltic, East Slavic, and Southeast European regions, respectively (Supplementary Table 5). Bottom: Classification of ancient individuals. For Lund and Trondheim, individuals are classified as outliers (non-Scandinavian; dark blue), non-locals (yellow) or locals (red). For Vilnius, individuals are classified as outliers (non-Baltic; dark blue) or Baltic (horizontal yellow-red stripes). The method used to identify outliers and non-local Scandinavians (the latter only for Lund and Trondheim) is indicated by bars beneath the classification results: PCA-based *D_M_* (dark grey) or IBD sharing (light grey). **B)** Summary of the classification from bottom panel of A.

Next, we used IBD sharing between ancient and present-day individuals, restricting to shared fragments longer than 6 cM (Supplementary Figs. 18 and 19), to obtain more fine-grained assessments of ancestry over time. For Lund and Trondheim, we used a reference set (“OmniExpress set”) comprising 29,525 present-day individuals from Scandinavia (Denmark and 13 sub-regions across Norway and Sweden), and adjacent non-Scandinavian regions (the Netherlands, Germany, Iceland, Shetland, and Orkney) (Supplementary Table 5). These non-Scandinavian populations were included because they are genetically close to Scandinavians and cannot be easily distinguished from them using only PCA-based *D_M_* (Supplementary Fig. 20, Supplementary Note 3). For Vilnius, we estimated IBD sharing using a subset of individuals from the UK Biobank who trace their ancestry to the Baltic, Eastern Europe, and neighbouring regions (Supplementary Table 5, Supplementary Fig. 21), restricting to fragments longer than 7 cM due to lower marker density. As with the PCA-based *D_M_* analysis, we used these results to assess the similarity of our ancient individuals to present-day groups from the same region (Fig. 3A middle panel). We used present-day inhabitants from Trøndelag (N=1345) as proxies for local Trondheim ancestry, and those from Skåne (N=841) and Denmark (N=1606) as proxies for Lund (Skåne was largely under Danish rule from the mid-10th century until 1658). As we did not have present-day individuals from Lithuania in the OmniExpress set, we used present-day individuals from the UK Biobank, originating from the Baltic region (N=139), including Lithuania, Latvia and Estonia, as present-day proxies for ancient individuals from Vilnius. In line with the PCA-based results, a greater proportion of post-BD than pre-BD individuals in Lund share most IBD with their corresponding present-day populations. Trondheim and Vilnius show the same pattern, although the differences are not significant (Fisher’s exact test; Lund: pre-BD 0.33, post-BD 0.67, *P*=3.4×10^-7^; Trondheim: pre-BD 0.49; post-BD 0.53, *P*=0.68; Vilnius: pre-BD 0.74, post-BD 0.87, *P*=0.28).

#### Detecting and classifying ancestry outliers

Given the diversity of ancestry observed at all three sites, we used tests based on a combination of *D_M_* from the West Eurasian PCA and IBD sharing to identify ancestry outliers: non-Scandinavian for Lund and Trondheim, and non-Baltic for Vilnius. Specifically, ancient individuals from Lund and Trondheim were classified as non-Scandinavian either if their squared *D_M_* values against present-day Norwegians (N=510), Swedes (N=516), and Danes (N=797) all exceeded the chi-square threshold (*D_M_* >4.93, corresponding to *P*<0.001) or if the pattern of IBD sharing indicated they were more closely related to non-Scandinavian populations in the OmniExpress set (Welch’s *t*-test *P*<0.05, see Methods). The same approach was applied to classify ancient individuals from Vilnius as non-Baltic: *D_M_* was calculated using present-day Lithuanians (N=121) and Estonians (N=55) from the West Eurasian PCA, and IBD sharing was assessed using individuals from the Baltic region in the UK Biobank dataset as Baltic references. We used a combination of *D_M_* and IBD sharing because leave-one-out cross-validation shows that IBD sharing better distinguishes closely related ancestries, such as Dutch and Danish (Supplementary Note 3). Across all three sites combined, the proportion of ancestry outliers declined markedly from the pre-BD to the post-BD period (Fig. 3B; Mantel-Haenszel test stratified by site, χ^2^ =54.4, *P*=1.64×10^-13^). This decline was significant in Lund and Vilnius, but not in Trondheim when the sites were tested separately (Fisher’s exact test: Lund: pre-BD 0.52 versus post-BD 0.09, *P*=1.37×10^-13^; Trondheim: pre-BD 0.16 versus post-BD 0.03, *P*=0.11; Vilnius: pre-BD 0.47 versus post-BD 0.05, *P*=3.71×10^-4^). In Lund, where some of the ancient individuals were specifically dated to the Black Death period, we also detected a marked reduction in the proportion of outliers when comparing pre-BD to during-BD individuals (Lund: 0.52 to 0.15, *P*=1.84×10^-5^). The decline in the proportion of outliers suggests less long-range immigration in the post-BD period.

For Lund and Trondheim, ancient individuals that were classified as Scandinavian were further subdivided into two categories: local and non-local. Locals are individuals whose ancestry indicates origins from the region of the sampling site, while non-locals are those whose ancestry indicates origins elsewhere in Scandinavia. This classification was obtained using *D_M_* and IBD analyses based on the subset of present-day Scandinavian individuals from the OmniExpress set (N=16,639, Supplementary Table 5, Supplementary Figs. 22-24). The present-day proxies for local ancestry were those described above: Denmark and Skåne for Lund, and Trøndelag for Trondheim. Ancient individuals were classified as non-local if their squared *D_M_* values versus the present-day proxies exceeded the chi-square threshold (*D_M_* >6.14, corresponding to *P*<0.001), computed on the first 15 PCs (Supplementary Fig. 4), or if they shared significantly more IBD with another Scandinavian region than with the present-day proxies (*P*<0.05, Welch’s *t*-test). The remaining individuals were classified as locals. We observed no significant change in the proportions of local and non-local Scandinavians between the pre-BD and post-BD populations (Fisher’s exact test: Lund *P*=0.20, Trondheim *P*=0.17), suggesting similar migration rates within Scandinavia pre– and post–Black Death.

#### Assessing the origin of outliers and non-locals

To further characterise the ancestry of the outliers identified above, we first used *D_M_* based on the West Eurasian PCA (Supplementary Notes 4, 5). In Lund, we see a high diversity of likely sources for outliers in the pre-BD period. Most outliers are consistent with present-day populations from Germany and the Benelux region (N=67), France (N=54), the British Isles and Ireland (N=42) (Fig. 4A,B), with a few showing greatest affinity to eastern Europe (N=14), the Baltics (N=7), and southeastern Europe (N=1). For post-BD Lund, the most likely source for the majority of outliers is Germany and the Benelux region. For Trondheim and Vilnius, outliers are less diverse in ancestry. In Trondheim, we observe a much higher rate of outliers pre-BD than post-BD, and most pre-BD outliers are consistent with present-day populations from either Iceland or the British Isles and Ireland (Fig. 4A,B). In Vilnius, most outliers show ancestry consistent with present-day Slavic speakers in East Europe (Fig. 4B), with two notable exceptions in the pre-BD group (Fig. 4A): LIT452 has ancestry similar to present-day and Medieval Southeast Europeans, while LIT448 is assigned 22% Asian ancestry in a supervised ADMIXTURE analysis based on 1000 Genomes populations^41^ (Supplementary Note 6).

**Figure 4:**
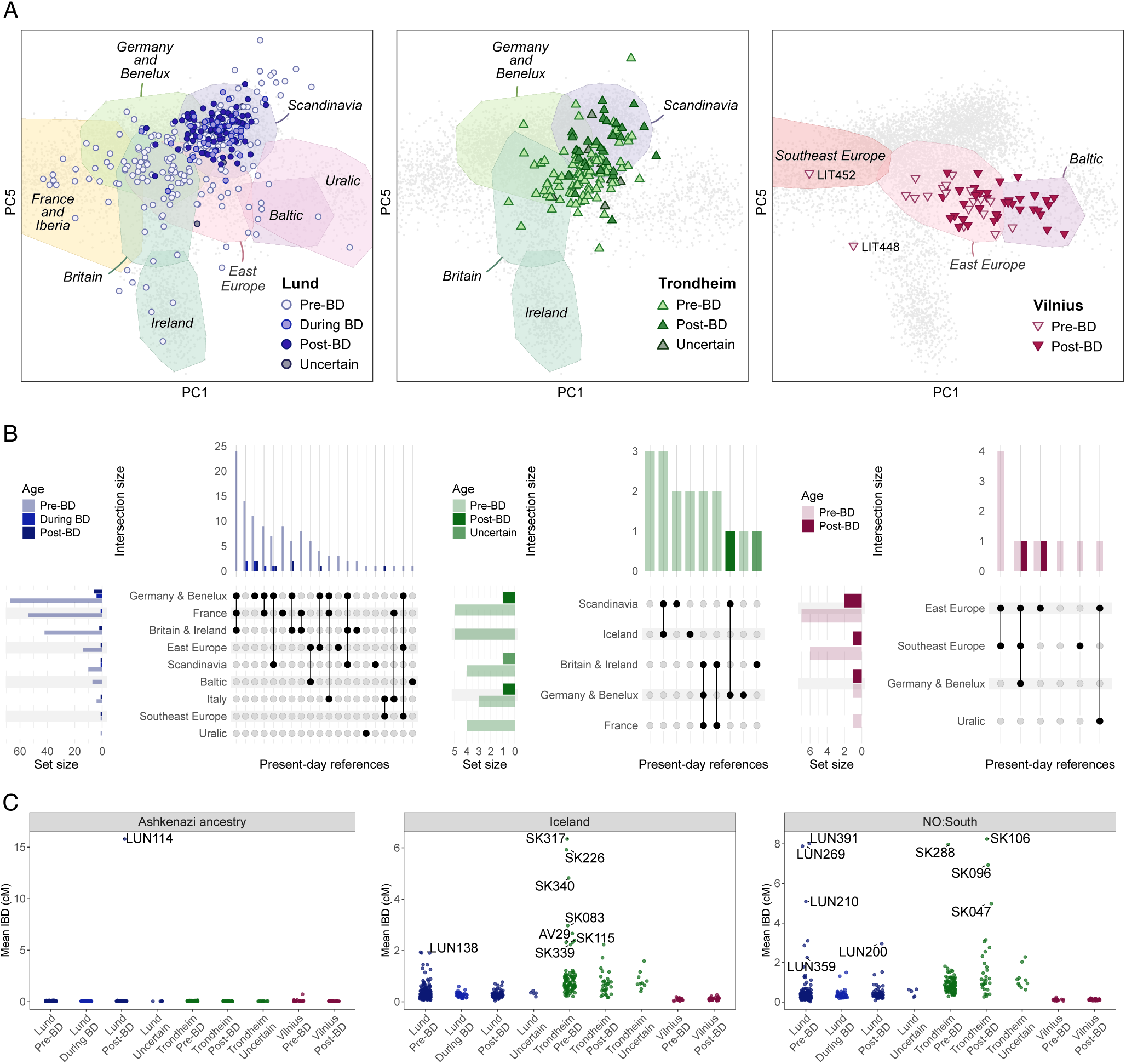
Ancestry characterisation of outliers using PCA and IBD. **A)** Projections of all ancient individuals onto the PCA of present-day West Eurasians, shown separately for the three sites. PC1 reflects north-south differentiation, while PC5 captures British-Irish ancestry. **B)** UpSet plot summarizing the regional assignments of ancient individuals classified as outliers according to PCA-based *D_M_*. Filled-in circles in a column mark a combination of non-excluded populations, with vertical bars above indicating the count of individuals showing that combination (“intersection size”). Horizontal bars to the left indicate the count of individuals that include the corresponding population among their non-excluded populations (“set size”). See Supplementary Figs. 34-36 for figures including all ancient individuals. **C)** IBD sharing between ancient individuals and selected present-day reference populations. Points show the average IBD sharing for each ancient individual, stratified by site and period along the x-axis, with the selected present-day reference population indicated in the top strip. Ancient individuals are labelled only when their IBD-sharing profiles support a likely origin related to the corresponding reference population after multiple-testing correction (see Methods for detail and Supplementary Fig. 33 for figures covering all reference populations).

To obtain finer-scale resolution of the likely origin of the outliers, we analysed their IBD sharing with present-day populations (Fig. 4C, Supplementary Note 5) using the UK Biobank set to provide geographic breadth and the OmniExpress set to provide well-powered tests for Northern European ancestries. We note that due to differing marker sets and minimum cM thresholds, absolute values of IBD sharing are not directly comparable between the two present-day reference sets. Among the particularly interesting outliers in Lund characterised in this way are two pre-BD individuals (LUN145 and LUN298) who show the highest IBD sharing with present-day individuals from Wales, but not elevated IBD sharing with individuals from Scotland or Ireland, strongly suggesting Welsh origin. One individual post-BD, LUN114, carries Ashkenazi Jewish-related ancestry (Fig. 4C, Supplementary Note 7). Finally, LUN101 (pre-BD) shares most IBD with present-day Finns (Fig. 4C) and exhibits elevated East Asian and Siberian ancestry in a 1000 Genomes supervised ADMIXTURE analysis (Supplementary Fig. 25). Among the outliers in Trondheim, seven pre-BD individuals share most IBD with present-day Icelanders (Figs. 3B and 4C). An Icelandic origin for these individuals is further supported by IBD sharing analyses against a larger set of present-day Icelanders (N=136,064) and strontium isotope results for three of the seven (Supplementary Fig. 26). In addition, one Trondheim individual (SK090, uncertain date) shares most IBD with present-day inhabitants of the Shetland Islands, which were under Norwegian rule until 1472 CE.

Our leave-one-out tests suggest that IBD-sharing tests based on the OmniExpress set can be used to distinguish between Scandinavian countries with fairly low error rates, and even between geographical regions within countries (particularly Sweden) but with greater error rates (Supplementary Fig. 24, Supplementary Note 3). Thus, we are also able to identify likely examples of migration within Scandinavia. For instance, three individuals from pre-BD Lund share most IBD with present-day southern Norwegians (LUN391, LUN269, LUN210). Several individuals from Trondheim share most IBD with present-day Danes, including one pre-BD individual (SK088) and two post-BD individuals (SK014, SK022). One post-BD individual from Trondheim (WF629) shares most IBD with present-day northern Scandinavians, while another from Lund (LUN98) shares most IBD with populations specifically from northern Sweden. In Trondheim, four post-BD individuals share IBD with present-day southern Norwegians (SK288, SK106, SK096, SK047), while one pre-BD individual shares most IBD with western Norway.

## Discussion

Due to the massive mortality attributable to *Y. pestis* during the second plague pandemic, it is often considered one of the strongest selective pressures to have affected human populations during the past millennium. In line with this expectation, one earlier study of immune-related loci in ancient individuals reported strong signals at several candidate genes, most notably ERAP2^16^. However, our analyses of more than five hundred ancient genomes from three northern European urban centres, spanning the arrival of the Black Death, did not reveal evidence for strong positive selection that affected the frequency of these or any other specific genetic variants. Given the similarity of these populations in terms of ancestry, sequence variation, and the strains of *Y. pestis* that affected them^42,43^, it is reasonable to assume that variants subject to strong selection at one of the locations would also be selected at the others. This assumption underpins our use of meta-analysis combining association analyses from Trondheim and Lund, and suggests that we should be able to replicate the findings of previous studies.

Importantly, using simple power calculations we demonstrate that our study is well-powered to detect frequency changes of the magnitude reported for variants suggested to be under selection in previous studies. However, the calculations also show that our study is only well powered to detect novel variants with selection-driven allele frequency changes larger than 0.12-0.26 when comparing pre-BD versus post-BD, with the exact threshold depending on the pre-BD allele frequency. When comparing pre-BD to modern individuals, the threshold range is 0.07-0.15. To detect variants with more subtle allele frequency changes, substantially larger sample sizes would be needed. We note that greater statistical power could be gained by comparing individuals who died during the pandemic to those who survived, since allele frequency differences between these groups are expected to be larger than those between pre-BD and post-BD individuals^44^. However, this sampling scheme might be hard to achieve in practice, given the uncertainty about causes of death for skeletal remains.

The lack of selection signals we observe might seem surprising given the massive mortality wrought by the second plague pandemic in northern Europe. However, we note two fundamental weaknesses in the underlying hypothesis that the pandemic caused large allele frequency changes at individual variants. First, most mortality was concentrated in an evolutionarily brief interval, where even strong selection pressure is expected to induce only modest allele frequency changes^44^; such subtle shifts are difficult to detect, even with our substantially larger sample sizes. Second, the hypothesis assumes that medieval Europeans carried reasonably common variants with large effects on survival against a novel strain of *Y. pestis*, which may simply be unfounded, especially given the lack of recent plague exposure in these populations. In addition to these considerations, we note that our stringent site-level filtering in the selection scan, including the Hardy-Weinberg equilibrium test, may reduce sensitivity for detecting selection signals in immune-related genes, many of which are highly polymorphic and particularly susceptible to genotyping uncertainty even in present-day DNA.

While we did not find evidence for a pronounced selective impact of the second plague pandemic, our findings revealed remarkably parallel declines in the diversity of ancestry consistent with reduced long-distance immigration to Trondheim, Lund and Vilnius after the arrival of the Black Death. These analyses reveal the considerable power provided by analyses of multidimensional PCA spaces and IBD sharing, applied to large reference data sets from present-day populations, to resolve fine-scale differences in ancestry for individuals from medieval Europe – even at the level of regional differences within countries. Although the geographical structure of genetic variation in present-day populations may differ subtly from those of the past, previous studies indicate that the key features of structure in present-day Scandinavia closely match those of medieval times^35,45,46^. The reduction in the diversity of ancestry was particularly pronounced in Lund, where around 52% of the pre-BD individuals traced their ancestry to regions outside Scandinavia, including the Low Countries, Britain and Ireland, France, and Central Europe. In contrast, after the arrival of the pandemic, such outliers were markedly rarer (9%), with a substantially greater proportion of individuals resembling the present-day inhabitants of this region (36% vs 76%, *P*=1.57×10^-9^). Similar striking results are seen for the proportion of outliers in Vilnius, with a consistent trend in Trondheim, albeit not statistically significant.

These results indicate that the ancestry profile at all three sites, and Lund in particular, was relatively cosmopolitan during the pre-BD period. Given that present-day populations from these regions show substantial continuity of ancestry over the last millennium^35,47^, it seems unlikely that the long-range immigration causing such high proportions of outliers pre-BD could have been longstanding. It is more likely that this extensive mobility was due to a combination of transient socioeconomic factors. In Scandinavia, these include the gravitational pull of wealth and power during the Viking Age and the slave trade associated with it^48^. Another crucial factor is the introduction of Christianity beginning in the late 10th century, with both Lund and Trondheim acting as early pivotal locations, attracting missionaries and emissaries from continental Europe and the British Isles^31,32^. Interestingly, our results are broadly consistent with historical and archaeological views of the spread of Christianity in Scandinavia, which indicate more diverse sources for Lund, including the British Isles until 1070 CE and thereafter Hamburg-Bremen, but primarily the British Isles for Norway^31–33^. The more cosmopolitan nature of pre-BD individuals from Lund is consistent with its more prominent role in the spread of Christianity, as evidenced by its designation as the archiepiscopal seat for all Scandinavia in 1104 CE^31,33,49^. A distinctive feature of the pre-BD individuals from Trondheim is that seven of the 15 outliers show clear indications of Icelandic ancestry. This is consistent with the close ancestral and cultural connections between Iceland and Norway^50,51^; further, Iceland was Christianised from Norway around 1000 CE and the Icelandic church was under the ecclesiastical authority of the Archbishop of Nidaros (Trondheim) from 1152 CE. Trondheim was also the primary residence of Norwegian kings until ca. 1217 CE and Icelandic medieval texts contain many accounts of Icelanders travelling there during the pre-BD period with some dying and being buried there^52^.

The influence of Christianity on pre-BD diversity in Lund and Trondheim raises an important potential caveat relating to ascertainment of skeletal remains for this period, which were recovered from Christian cemeteries. Many of the individuals we label pre-BD lived during periods when the local population was substantially pagan, so the urban churchyard burials we analyse might be biased towards migrant Christians. While such bias might inflate differences in the diversity of ancestry we observe between the pre-BD and post-BD periods, there is good reason to expect a genuine reduction in diversity. Indeed, this temporal transition also broadly coincides with the end of the Viking Age and the onset of the Little Ice Age^29,30^, both of which contributed to the marginalisation of Scandinavia and a decline in the factors that would previously have attracted long-range immigrants. No less important was the unprecedented mass mortality that accompanied the arrival of the second plague pandemic, inevitably exacerbating economic decline, in part by disrupting commercial, ecclesiastical and political networks that had previously facilitated long-distance mobility^53^. Such disruptions would have reduced the movement of merchants, clerics and craftsmen who had contributed to the cosmopolitan character of medieval urban centres in Scandinavia. Notably, the decline we observe is specific to long-range mobility. While short-range mobility may also have been affected by these events, the temporal differences we observe in the proportions of non-local Scandinavian individuals in Lund and Trondheim are not statistically significant.

In Vilnius, part of the observed decline in the diversity of ancestry may also be explained by ascertainment biases linked to Christianity and migration. Christianisation in Lithuania only began to intensify in the decades following the Black Death, nearly four centuries after Scandinavia had been largely converted, and was initially driven more by Eastern Orthodox Slavs than by Catholics from the west^34^. The pre-BD individuals in Vilnius were sampled from an Orthodox churchyard in the old Ruthenian (East Slavic–speaking) quarter and are mostly assigned Slavic- rather than Baltic-associated ancestry, consistent with pagan Lithuanians being under-represented in burial sites for migrant Christian Slavs. Additionally, the presence of two strong ancestry outliers in the pre-BD churchyard coincides with a documented campaign by the Lithuanian monarch Gediminas to attract far-flung, generally Christian colonists to the newly founded capital^54^ – a clear mechanism for genuine high diversity during this period. In contrast, post-BD individuals from a 15-16th century plague-associated non-churchyard site and a 17th century churchyard show a shift towards Baltic-associated ancestry, with the latter possibly reflecting the completed Christianisation of Lithuanian Balts.

In summary, a comparison of pre-BD and post-BD individuals at Lund, Trondheim and Vilnius revealed the genetic impact of far-reaching changes that affected fundamental aspects of culture, religion, and economic organisation, which ultimately led to a decrease in long-range migration. Although the second plague pandemic was only one of several factors contributing to these changes, it is nonetheless known to have inflicted enormous social and demographic repercussions in the studied regions and elsewhere in Europe. Our analyses of more than five hundred ancient genomes revealed no credible signals of strong positive selection. However, our results show that much larger numbers of ancient genomes will be needed to determine the biological impact of the pandemic, in terms of altering the frequency of individual genetic variants due to their impact on differential reproductive success.

## Online methods

### Skeletal material analysed

The skeletal material analysed in this study derived from three principal locations: Lund (formerly part of Denmark, today part of Sweden), Trondheim (Norway) and Vilnius (Lithuania). Medieval Lund, founded in the late 10th century, was a major political and ecclesiastical centre of the Danish kingdom and the archiepiscopal seat of the Nordic countries^55^. The ancient genomes analysed here derive primarily from teeth excavated from the Trinitatis (Trinity) churchyard, the largest medieval burial ground in Lund, in use from ca. 990 to 1536 CE. The cemetery has been extensively excavated and is well dated through stratigraphy, dendrochronology, and radiocarbon analysis, allowing burials to be assigned to established chronological phases spanning the foundation of the town through the Late Middle Ages, including periods affected by the Black Death. Trondheim (medieval Nidaros) emerged in the late 10th century as a royal and commercial centre and later became the ecclesiastical capital of medieval Norway^56,57^. Individuals included in this study were excavated from four churchyards representing distinct social contexts and time periods, ranging from medieval urban parish and monastic settings to post-medieval burials associated with Nidaros Cathedral. These sites capture variation in status, function, and chronology from the 12th to the 19th centuries, providing a diachronic and socially heterogeneous sample from the same urban centre. Vilnius developed as a major multicultural centre following the expansion and Christianization of the Grand Duchy of Lithuania from the 13th century onward^34,58^. Skeletal material, mainly temporal bones (with some teeth), was sampled from three burial grounds reflecting different social and historical contexts. These include an early Orthodox Christian cemetery dating from the 13th–15th centuries, a 15th–16th century burial ground discovered outside the medieval city walls with evidence of mass and atypical burials, and a 17th-century Orthodox cemetery associated with the Church of the Holy Spirit. Together, these sites capture early urbanization, religious diversity, and later early modern population dynamics in Vilnius. Additional details on the skeletal remains and sites can be found in the supplemental information (Supplementary Table 1, Supplementary Note 1).

### DNA library preparation, enrichment and sequencing

Several different methods were used to extract the DNA, convert the DNA into sequencing libraries, and sequence the libraries. Full details for each sample are indicated in Supplementary Table 2. In brief, the genomes from Trondheim individuals are a mixture of (i) previously published target capture and shotgun sequence data as described in Gopalakrishnan et al. (2022), supplemented with (ii) shotgun sequence data generated from additional new tooth and bone samples generated for this study. DNA from all new Trondheim samples was extracted following Allentoft et al. (2015)^59^, after which DNA was converted into ‘BEST’ double stranded sequencing libraries for either Illumina (following Carøe et al., 2018^60^) or BGIseq (following Mak et al., 2017^61^) sequencing. DNA from a subset of the Trondheim libraries was subjected to USER treatment to remove deaminated cytosines, following Rohland et al. (2015)^62^. DNA from Lund and Vilnius tooth and bone samples was principally extracted by incorporating an initial predigestion at 37° for 30 minutes using 1 mL of a digestion buffer consisting of 0.463M EDTA pH8, 10mM Tris, 0.5% N-laurylsarcosine, 30%, 1x Phenol red and ca 0.2 mg/ml Proteinase K^63^. Subsequently, after discarding the supernatant an additional 4ml of digestion buffer was added to each sample, after which they were incubated overnight at 37°C on a rotor. DNA was purified from 1.8ml of the digest using a modified version of Qiagen PB buffer (cat: 19066) and centrifugation through Roche large volume silica columns (cat: 05114403001) following McColl et al. (2024)^64^, with final elution of DNA using 65 µL Qiagen EBT buffer into 1.5ml Eppendorf LoBind tubes. The extracted DNA was then converted into single-strand DNA Illumina libraries following the ‘SCR’ protocol of Kapp et al. (2021)^65^. While the majority of the sequence data from Vilnius and Lund was subjected to USER treatment, a subset was left untreated to enable assessment of DNA damage profiles.

### Processing and alignment of sequencing reads

All sequencing reads were processed using an in-house pipeline at deCODE genetics. Firstly, AdapterRemoval v2 ^66^ was used to remove adapter sequences and stretches of consecutive low-quality bases and ambiguous nucleotides (N’s) and discard reads shorter than 25 bp, which are commonly produced by postmortem DNA degradation. Processed reads were then aligned to the human reference genome GRCh38 using BWA backtrack algorithm implemented in ‘bwa aln’ v0.7.17^67^, with parameters optimised for ancient DNA (aDNA) (-n 0.01 -o 2 -l 1024 -q15 -k 1). Mapped reads were merged by library, followed by removal of PCR duplicates using Picard (PicardTools v.2.21, http://broadinstitute.github.io/picard/) and filtering of reads with mapping quality <20 using SAMtools^68^. Libraries were subsequently merged at the sample level, and sequencing depth and genomic coverage were calculated using BEDtools^69^ and an in-house Python script. Finally, mapDamage2.0^70^ was applied to rescale base quality scores, excluding likely damaged bases. Summary statistics for all sequenced libraries are provided in Supplementary Table 1.

Multiple approaches were applied to validate the authenticity and quality of the dataset. First, we confirmed that all libraries produced fragment length distribution typical of aDNA (Supplementary Table 1) and verified nucleotide misincorporation patterns consistent with post-mortem damage using mapDamage2.0. Secondly, X-chromosome contamination was assessed for genetically male individuals using ANGSD^71,72^, with analyses restricted to reads with a base quality of ≥ 20 and a mapping quality of ≥ 20. ANGSD implements two estimators: (i) a method using all available bases, which provides greater statistical power under the assumption of independent errors across reads and loci, and (ii) a method that randomly samples a single base per locus to reduce bias, albeit at the cost of reduced precision. Additionally, autosomal contamination was evaluated using an in-house implementation of ContamLD^73^, which leverages breakdowns in linkage disequilibrium to infer contamination levels. A threshold of 10% was applied to the ContamLD estimate to identify contaminated genomes. Finally, biological sex was determined for all individuals based on the relative X- and Y-chromosome coverage following Skoglund et al. (2013)^74^.

### Genotype imputation

Genotype likelihood based imputation was performed separately for each sample using GLIMPSE2^37^. Imputation used a phased reference panel from deCODE genetics (“deCODE 50k”), comprising 50,129 individuals from diverse global populations with enriched representation of European, and particularly Scandinavian, ancestries. The reference panel contains 99,610,455 loci, including bi-allelic and multi-allelic SNPs as well as INDELs. GLIMPSE2 performed genotype imputation and phasing directly from the BAM files in overlapping 2-Mb windows across each chromosome. Only genomes with a minimum autosomal depth of coverage of 0.1x and no clear signs of contamination were included in the imputation step. Based on the set of included genomes, imputation INFO scores were recalibrated using the impute-info plugin in BCFtools.

### Haplogroup calling

mtDNA and Y-chromosomal haplogroups (Supplementary Table 1) were called using HaploGrouper^75^. mtDNA haplogroups were based on the PhyloTree build 17 phylogeny, and Y-chromosome haplogroups were based on a previously defined fork of the 14.196 ISOGG 2019 phylogeny, as described in the HaploGrouper Gitlab repository.

### Kinship analysis and inbreeding

We estimated the degree of relatedness among ancient individuals using identity-by-descent (IBD) inferred from imputed genotypes. IBD fragments shared between ancient individuals were called using ancIBD^76^. Loci and allele frequency files bundled with ancIBD, together with the recombination map included in the .snp file for AADR v51.1^77^, were lifted over to hg38 to match the genotype data generated in this study. Imputed genotypes were restricted to the required loci and VCF files for each chromosome were converted to HDF5 format. These files were used for input for ancIBD-run using the --IBD2 flag, which enables joint detection of genomic segments in which a pair of individuals shared one allele identical by descent (IBD1), and segments in which they shared both allele identical by descent (IBD2). All remaining parameters were set to their default settings. Degrees of relatedness were inferred based on the estimated proportions of the genome shared as IBD1 and IBD2. Runs of homozygosity in imputed genotypes were estimated as in Ellegaard et al. (2024)^46^, with the MAF>0.03 threshold defined with respect to the deCODE 50k imputation panel.

### Principal components analysis

We performed PCA projection using present-day reference datasets: (i) a European dataset comprising 168,599 SNPs from 9,052 individuals representing 67 fine-scale West Eurasian population labels, previously described in Rodríguez-Varela et al. (2023)^35^; and (ii) a Scandinavian dataset comprising 16,639 individuals from 41 geographical divisions plus 887 Icelanders, described in Mattingsdal et al. (2021)^45^. To improve power, these fine-scale divisions were merged into larger groups based on geography, linguistics, and hierarchical clustering of mean PC coordinates: 17 for the West Eurasian set, and 14 for Scandinavia. Details of the present-day reference datasets used for PCA and other ancestry-related analyses are summarised in Supplementary Table 5. Principal components were calculated on the present-day reference individuals using the *smartpca* module in EIGENSOFT^78^, with LD regression applied using the parameters ‘‘ldregress: 200’’ and ‘‘ldposlimit: 100000”. Ancient individuals were subsequently projected onto the inferred principal components using an in-house script based on imputed genotypes. A scalar correction factor was applied to reduce the shrinkage of projected PCA coordinates using the method described in Lee et al. (2010)^79^.

### Selection scan

To investigate genetic variants associated with survival during the Black Death, we performed selection scans. For each site, we conducted a genome-wide association study (GWAS) to directly compare allele frequencies between pre-BD and post-BD individuals. To account for population structure and relatedness, association testing was performed using a linear mixed model implemented in GEMMA^80^. The model included a genetic relationship matrix (GRM) as a random effect, with sex and the first seven PCs from the West Eurasian PCA projection as covariates. The number of PCs was chosen based on repeated hold-out analyses of present-day West Eurasian reference populations, as described in the ancestry analyses below and shown in Supplementary Fig. 4. For each GWAS run, the GRM was estimated using a set of high-quality SNPs filtered using the following criteria: a minor allele frequency (MAF) ≥ 0.05, calculated separately among the individuals included in that site-specific GWAS dataset, an imputation INFO score ≥ 0.98 and location within the 1000 Genome Project strict accessibility mask for hg38^41^. We further excluded SNPs flagged for low quality by deCODE genetics pipelines^81^ and those located in regions of high linkage disequilibrium from GRM calculation.

For each GWAS, we applied stringent site-level filtering criteria. We excluded variants with MAF < 0.05, with MAF calculated separately among the individuals included in the corresponding site-specific GWAS dataset. We also excluded variants with high imputation uncertainty, defined by an INFO score < 0.95 or > 5% of genotypes having a maximum genotype probability (GP) below 0.9 (Supplementary Fig. 27). In addition, we excluded variants flagged for low quality by deCODE genetics pipelines in the present-day sequences forming the imputation panel, variants significantly deviating from Hardy-Weinberg equilibrium (HWE) in all pre-BD individuals after accounting for population structure using PCAngsd (*P*<10^-6^), and multi-allelic indels. Finally, we removed markers located within 5 bp of another variant, as such high-density sites are often missing or extremely rare in gnomAD non-Finnish European population, suggesting technical artifacts.

To increase statistical power, we additionally performed GWAS comparing pre-BD individuals with present-day individuals, separately for Lund and Trondheim. This effectively increased the control sample size from N_post_=75 to N_present_=380 in Lund and from N_post_=30 to N_present_=1182 in Trondheim. The same site-level filtering criteria were applied to each of the pre-BD versus present-day GWAS outputs, with the additional exclusion of loci significantly deviating from HWE (*P*<10^-6^) in each present-day cohort. After filtering, the number of retained markers ranged from 5,633,253 to 5,655,420 across the GWAS datasets. This analysis assumes that allele frequency changes from the post-BD period to the present-day era are largely independent of the Black Death.

To assess the effect of markers across sites, we performed meta-analyses of the GWAS summary statistics using METAL^82^. Meta-analyses were conducted separately for the two comparison sets: pre-BD versus post-BD individuals, and pre-BD versus present-day individuals. For each comparison set, meta-analysis was performed using the union of filtered loci from the two sites. We used an inverse-variance-weighted approach and applied the significance threshold adjusted according to functional categories, as suggested by Sveinbjornsson et al.^39^. Lead variants were evaluated using regional association plots generated with LocusZoom^83^ and annotations from Ensembl VEP (release 100)^84^.

### Change of allele frequency at previously reported loci

We examined allele frequencies at loci reported to be under selection driven by the Black Death in previous studies. These included the four loci reported by Klunk et al. (2022)^16^, a putative locus reported by Hui et al. (2024)^19^, variants in the Toll-like receptor (TLR) gene cluster^15^ and *CCR5*-delta32. We compared the allele frequency between the pre-BD and post-BD individuals at these loci. Due to the small sample size in Vilnius, temporal allele-frequency comparisons were restricted to Lund and Trondheim. We also included allele frequencies from present-day proxy populations for these two sites. Confidence intervals for allele frequencies were estimated assuming binomial sampling of alleles.

### Power analysis

To determine the magnitude of allele frequency differences that our study was well powered to detect, we performed genotype-based power calculations for both pre-BD versus post-BD, and pre-BD versus present-day comparisons. The significance thresholds and the sample sizes for this power analysis were matched to those used in our selection analyses. For each combination of pre-BD and comparison-cohort allele frequencies, we performed 10,000 simulation replicates. In each replicate, we generated diploid genotypes for individuals in each cohort assuming Hardy-Weinberg equilibrium, which corresponds to a binomial model: *G_ij_* ∼ Binomial(2, *p_i_*), where *p_i_* is the allele frequency in cohort *i*, and *G_ij_* is the simulated genotype for individual *j* in cohort *i*. We then tested for association using a linear regression model with cohort as the outcome and genotype dosage as the predictor, matching the fixed-effect structure of GEMMA linear mixed model used in the selection analyses but without the random-effect component. Statistical power was estimated as the proportion of replicates in which the genotype effect was significant at the specified threshold.

Two significance thresholds were considered: a genome-wide significance threshold (α=1.1×10^-8^) corresponding to the multiple-testing burden in GWAS and a nominal significance threshold (α=0.05) for evaluating previously reported variants. For each significance level, simulations were performed across combinations of pre-BD and comparison-cohort allele frequencies ranging from 0% to 100% in increments of 0.5%. A combined MAF threshold of 5% across the pre-BD and comparison cohorts was applied. Power calculations were performed using sample sizes corresponding to the Lund individuals, the Trondheim individuals and the combined meta-analysis. For previously reported variants, we specifically estimated statistical power using the allele frequencies reported in the earlier studies to assess the expected probability of replication in our datasets.

### Calling identity-by-descent segments to present-day populations

We identified IBD fragments shared between the ancient individuals and two sets of present-day reference populations using an in-house Python script. For this analysis, we used two present-day reference panels. The first was UK Biobank^81^, which includes a very large number of individuals with ancestry from the British Isles, as well as individuals with ancestry from many other regions of Europe and elsewhere, although with relatively small sample sizes for each non-British population. The second was the OmniExpress European dataset (expanded from the Scandinavian reference panel described in^35^ using the same filtering procedures), which has a narrower geographic focus but substantially larger sample sizes. Specifically, this panel includes 16,639 present-day Scandinavian individuals divided into 14 groups, together with 887 Icelanders, 6,322 Dutch, 1,294 Germans, 2,174 individuals from Shetland and 1,325 individuals from Orkney. The Shetland and Orkney groups were used as proxies for British ancestry.

The method is based on an algorithm similar to IBIS^85^ implemented in Python, which takes unphased, imputed genotypes as input and defines candidate IBD segments by breaking segments at sites with opposite homozygotes. Due to imputation inaccuracy at rare variants, IBD calling was restricted to variants with minor allele frequency (MAF) > 0.02 in our imputation reference panel. IBD fragments were retained only if they were supported by at least 200 loci and had a rate of opposite homozygotes <0.3%. We further filtered IBD fragments based on the number of loci falling in long (>1000 bp) gaps in the 1kGP strict accessibility mask, corresponding to regions outside the stringently accessible portion of the genome. Fragments were kept only if they had >100 markers outside these gaps per centimorgan of total called length. For the Orkney and Shetland reference populations in the OmniExpress panel, this threshold was relaxed to >95 markers per centimorgan due to their slightly lower marker density. We also imposed a minimum genetic length threshold, which we adapted to the marker density of the reference panels: 6 cM for the OmniExpress European panel, which was used for ancestry classification in the Lund and Trondheim analyses, and 7 cM for the UK Biobank panel, which was used for ancestry classification in the Vilnius analysis.

Reference individuals from the UK Biobank were filtered and grouped as follows. As demonstrated previously^81^, UMAP of principal components reveals clusters of individuals with British-Irish ancestry, non-British European ancestry, and Ashkenazi ancestry, which was corroborated by the same K=5 supervised ADMIXTURE procedure described below. The British-Irish and non-British European clusters were subclassified according to birth country, with further grouping of low-count birth countries based on geography (Supplementary Table 5). When creating groups for France, Germany, the Netherlands, and Scandinavian countries, individuals in the British cluster self-reporting “White Other” ethnicity were included due to imperfect distinction between British ancestry and ancestry typical of these regions.

### Comparing diversity of ancestry between pre-BD and post-BD

To test for temporal differences in diversity of ancestry, we measured extent of dispersion and distance to present-day reference populations based on PCA. For each site, we compared the dispersion of pre-BD and post-BD individuals in the West Eurasian PCA using covariance matrices calculated from the first seven PCs. Multivariate dispersion was summarised using the determinant of the covariance matrix, which reflects the volume occupied by individuals in PCA space. Differences between pre-BD and post-BD covariance matrices were assessed using Box’s M test.

We next calculated the Mahalanobis distance (*D_M_*) between each ancient individual and present-day populations. We determined the number of the PCs to include using repeated hold-out analyses of the present-day reference populations. In each of ten replicates, we held out one individual from each population and calculated its *D_M_* to all populations using different counts of PCs. We chose the number of PCs that maximised the rate at which the correct population showed the lowest *D_M_*, as the number of PCs had little effect on the rate at which the true population was incorrectly excluded (Supplementary Fig. 4). Based on this criterion, we used the first seven PCs for *D_M_* calculations in the West Eurasian PCA. The same procedure was applied to the Scandinavian PCA, except that ten individuals from each population were held out in each replicate. Based on the same criterion, we used the first 15 PCs for *D_M_* calculations in the Scandinavian PCA (Supplementary Fig. 4). Differences in *D_M_* to present-day proxies from the same region between pre-BD and post-BD individuals were assessed using Wilcoxon tests.

To obtain a more fine-grained assessment of diversity of ancestry over time, we compared patterns of IBD sharing between ancient individuals and present-day reference populations. We used IBD sharing with the OmniExpress European dataset for Lund and Trondheim, and used IBD sharing with the UK Biobank dataset for Vilnius. As in the PCA-based *D_M_*, we used these results to assess the similarity of ancient individuals to present-day groups from the same region. Specifically, we used present-day individuals from Trøndelag, Norway for Trondheim, and individuals from Denmark and Skåne, Sweden as proxies for Lund (Skåne was largely under Danish rule from the mid-10th century until 1658^86^. For Vilnius, we used present-day individuals originating from the Baltic region (N=139), including Lithuania, Latvia and Estonia, as proxies because of the limited sampling resolution available in the UK Biobank. For each ancient individual, we identified the present-day population with which it shared the highest amount of IBD. We then compared the proportion of pre-BD and post-BD individuals who shared most IBD with the corresponding present-day proxies using Fisher’s exact tests.

### Ancestry classification

To quantify the patterns of ancestry change, we developed a two-step framework that combines PCA-based *D_M_* and IBD sharing to detect outliers among ancient individuals at each site. Outliers are defined as individuals with non-Scandinavian ancestry in Trondheim and Lund, and non-Baltic ancestry in Vilnius. First, we identified individuals that deviated significantly from the distribution of relevant present-day reference populations using *D_M_* calculated from the first seven PCs of the West Eurasian PCA. Squared *D_M_* were compared to a χ^2^ distribution with seven degrees of freedom, using a *P* value threshold of 0.001, a commonly used criterion for multivariate outlier detection^87^. For Trondheim and Lund, individuals were classified as outliers if they show significantly large distances from the distributions of present-day Danes, Swedes and Norwegians. For Vilnius, individuals were classified as outliers if they showed significantly large distances from the distributions of present-day Lithuanians and Estonians; Latvians were excluded from the reference set due to the small sample size (N=7).

Second, to better detect ancestry outliers closely related to the present-day proxy populations used in the PCA-based test, we performed IBD-based tests using summary statistics derived from the IBD fragments identified above. For Trondheim and Lund, we used the OmniExpress reference panel and divided the reference populations into Scandinavian and non-Scandinavian groups. The Scandinavian group comprised 14 regional populations, whereas the non-Scandinavian group comprised Dutch, German, Icelandic, Shetland and Orkney populations. For each ancient individual, we identified the Scandinavian population and non-Scandinavian population with the highest average IBD sharing with that individual, and then compared IBD sharing between these two populations using a one-sided Welch’s *t*-test (*P*-value threshold of 0.05). Individuals were classified as outliers if they showed significantly greater IBD sharing with the highest-sharing non-Scandinavian population than with the highest-sharing Scandinavian population. For Vilnius, we used the UK Biobank panel and divided the populations into Baltic and non-Baltic groups. The Baltic group included Lithuanian, Estonian and Latvian (collapsed due to small sample sizes), whereas the non-Baltic groups included populations from Poland, East Slavic regions, Southeast Europe, Germany, the Netherlands, Denmark, Sweden and Norway. For each Vilnius individual, we identified the Baltic population and non-Baltic population with the highest average IBD sharing with that individual, and compared IBD sharing between these two populations using a one-sided Welch’s t-test (*P*-value threshold of 0.05). Individuals were classified as outliers if they showed significantly greater IBD sharing with the highest-sharing non-Baltic population than with the highest-sharing Baltic population. Across all three sites, the final set of outliers consisted of individuals identified by either the PCA-based or IBD-based analysis.

Among the individuals classified as having Scandinavian ancestry from Trondheim and Lund, we further detect non-local individuals using the same two-step framework. Present-day inhabitants from Trøndelag were used as the local proxy population for Trondheim, and those from Skåne and Denmark were used as the local proxy populations for Lund. First, we identified individuals that deviated significantly from the local proxy populations using squared *D_M_* calculated from the first 15 PCs of the Scandinavian PCA (χ^2^ test, df=15, *P*<0.001). Because PCA has very limited power to resolve fine-scale structure within Scandinavia, whereas IBD sharing is more effective at capturing recent shared ancestry and fine-scale population structure^88^, we complemented the PCA-based analysis with IBD-based tests using the 14 Scandinavian populations in the OmniExpress dataset. For each ancient individual, we identified the local proxy population and non-local Scandinavian population with the highest average IBD sharing, and then compared IBD sharing between these two populations using a one-sided Welch’s *t*-test (*P*-value threshold of 0.05). Individuals showing significantly greater IBD sharing with the highest-sharing non-local population than with the local proxy population were classified as non-local. The final set of non-local individuals included those identified by either the PCA-based or IBD-based analysis.

### Validation of ancestry classification

To evaluate the performance and robustness of our ancestry classification method, we performed leave-one-out cross validation using present-day populations (Supplementary Note 3). For the PCA based analysis, we used *D_M_* calculated in PCA space. One individual at a time was masked from populations with N>40 and tested for exclusion from the reference distributions by comparing squared *D_M_* to a chi-squared distribution. For broad ancestry outlier detection, *D_M_* were calculated from the first seven PCs of the West Eurasian PCA (Supplementary Fig. 21). Specifically, Danes, Swedes and Norwegians were used as Scandinavian references, and Lithuanians and Estonians were used as Baltic references. A separate leave-one-out analysis based on the Scandinavian PCA was performed in the same way, using *D_M_* calculated from the first 15 PCs (Supplementary Fig. 22). For the IBD-based analysis, we performed leave-one-out tests using 100 randomly selected individuals from each reference population in the OmniExpress European dataset. For each masked individual, we tested consistency with each reference population using IBD sharing summary statistics. Individuals were rejected as inconsistent with a given population if they shared significantly more IBD with any other population than with that population, based on a Welch’s *t*-test (Supplementary Fig. 23).

In addition to leave-one-out cross validation, we compared our PCA- and IBD- based ancestry assignments to results from *qpAdm*^89^, a tool commonly used to model admixture using both modern and ancient source populations. Our findings indicate that all three methods converge in their assignments of source populations for the ancient individuals included in this study (Supplementary Note 4).

### Supervised ADMIXTURE

ADMIXTURE was run in supervised mode using two different present-day reference sets as training populations: a 1000G panel focused on K=5 global ancestry groups^35^ and a K=2 Norse-Gaelic panel^18^. Variants and individuals were filtered as described in the relevant citations.

### Characterisation of ancestry in outliers

As we identified a substantial number of individuals with non-Scandinavian ancestry in Lund and Trondheim, and several individuals with non-Baltic ancestry, we explored the likely sources of these outliers using *D_M_* based on the West Eurasian PCA. To provide a broader geographic overview, we grouped present-day populations into regions, as described in Supplementary Notes 4 and 5. For each outlier, we identified the regions with which the individual was statistically consistent by comparing squared *D_M_* values to a chi-square distribution ( χ^2^ test, df=7, *P*>0.01). Regions that could not be excluded as potential ancestry sources were summarised by site and period.

To characterise the ancestry of outliers at higher resolution, we analysed their IBD sharing with present-day populations. We used two present-day reference datasets: the UK Biobank and the OmniExpress European dataset. For each dataset, an individual was assigned a likely ancestry source only when its IBD sharing with a given present-day population was significantly higher than with all other reference populations in the same dataset using Welch’s *t*-test after Bonferroni correction for multiple testing, adjusted for the number of outliers and reference populations tested. These individuals are labelled in Figure 4c and Supplementary Figure 31.

## Code availability

Custom scripts used for the analyses in this study have been deposited in a public GiHub repository and are available at https://github.com/ivanliu3/second_plague_pandemic_genomics.git

## Acknowledgements

The authors thank the NTNU University Museum, the Norwegian National Committee for Research Ethics on Human Remains, Kulturen Museum Lund and Vilnius University for permitting and supporting the sampling of the skeletal remains.

The authors also thank Bent Petersen for computational support and Anders Albrechtsen for insightful discussions.

Finally, the authors thank the funding agencies that made the study possible: MTPG has received funding from Danish National Research Foundation (DNRF143) and The Carlsberg Foundation (CF18-1109), PG has received funding from the Austrian Science Fund FWF (P36433), XL and IM were supported by a Villum Young Investigator grant to IM (VIL19114), and this publication has emanated from research conducted with the financial support of Taighde Éireann – Research Ireland, under Grant number 21/RC/10294_P2 at FutureNeuro Research Ireland Centre for Translational Brain Science.

