## Supplementary Figures and Notes for "Genomic impact of the second plague pandemic on three human populations"

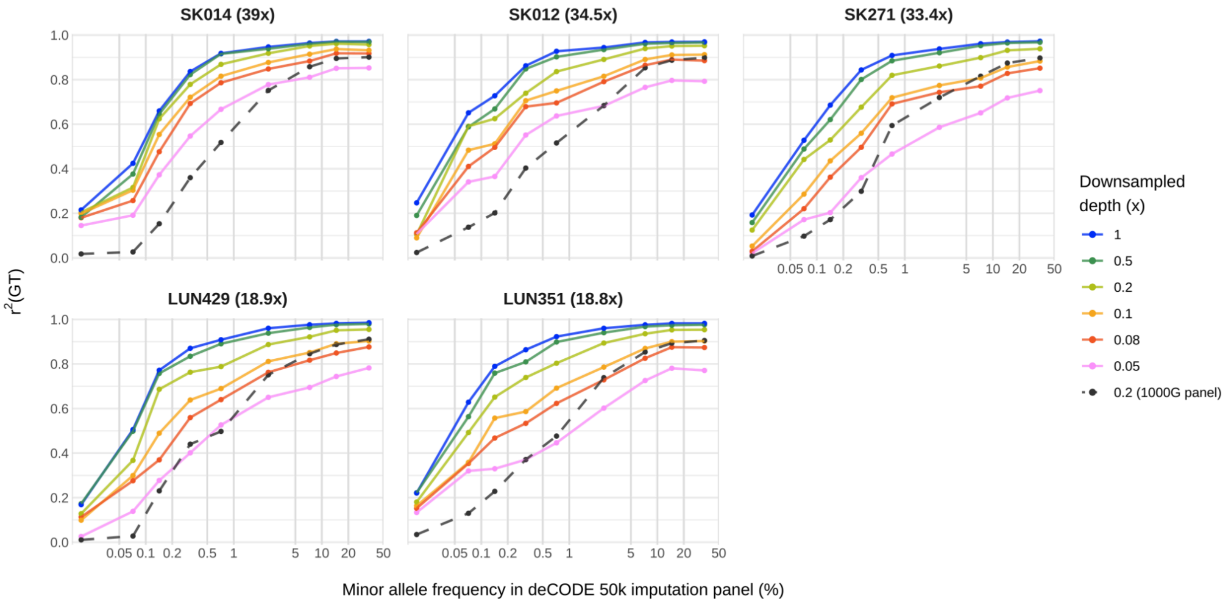


**Supplementary Figure 1.** Imputation accuracy for high-depth individuals after downsampling, stratified by minor allele frequency (*x*-axis). Sample name and original depth are indicated by panel titles. Accuracy (*y*-axis) is defined as the $r^2$ between observed genotypes and imputed genotypes hard-called using a minimum genotype likelihood of 0.9. Results are shown for downsampling to sequencing depths of 0.05x, 0.08x, 0.1x, 0.2x, 0.5x, or 1x (lines) followed by imputation using GLIMPSE2 and our study's "deCODE 50k" panel. Imputation at 0.2x depth using a 1000 Genomes reference panel (black) is shown for comparison to illustrate that the use of our much larger reference dataset for imputation leads to markedly improved accuracy.

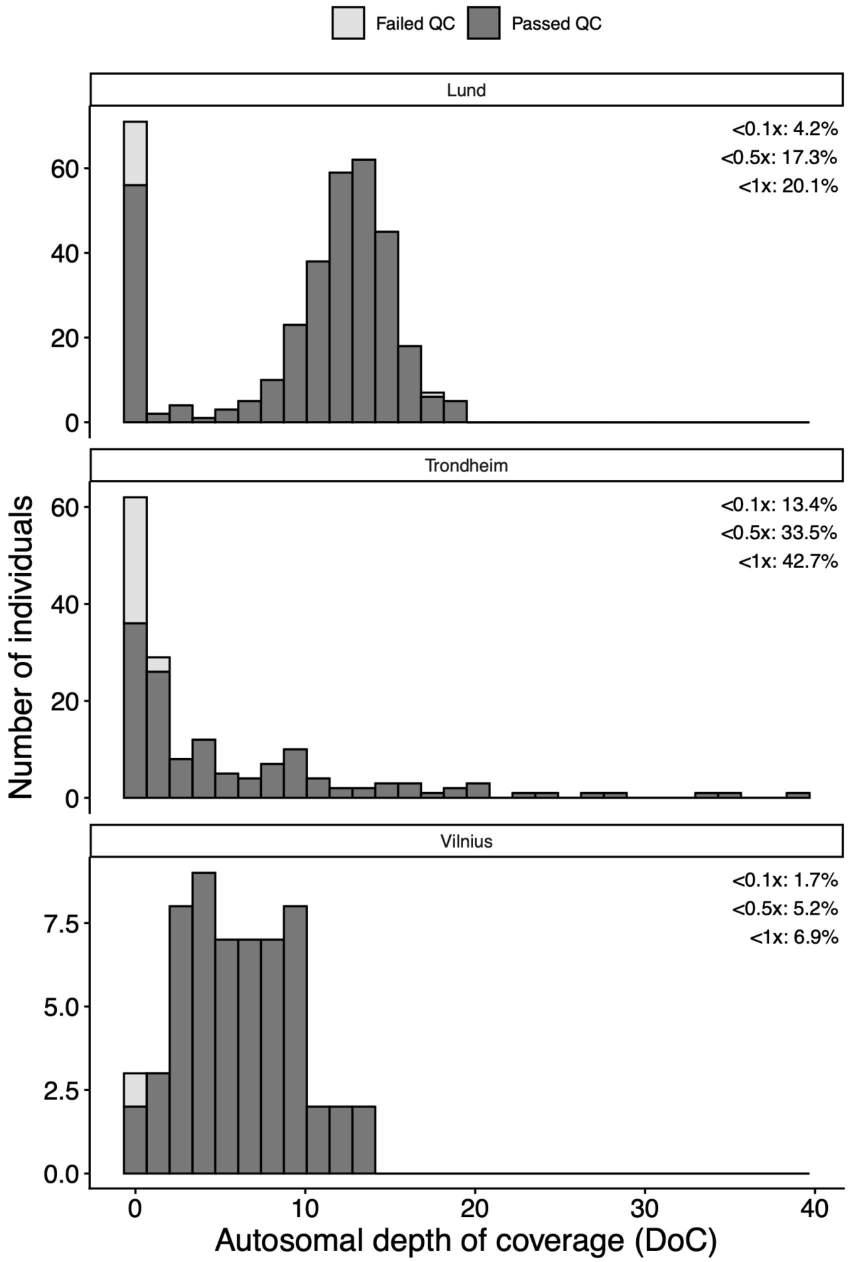


**Supplementary Figure 2.** Distribution of autosomal depth of coverage (DoC) across sampling sites with samples shaded according to whether they passed or failed QC. For each site, the percentage of samples before QC with autosomal depth of coverage below 0.1x, 0.5x, and 1x is shown within the corresponding panel. Descriptive statistics of DoC for samples that passed versus failed QC were as follows: Lund, passed QC N=337, mean=10.30x, median=12.00x, range=0.10-18.86x; failed QC, N=16, mean=1.10x, median=0.04x, range=0.00-16.96x. Trondheim, passed QC N=135, mean=6.13x, median=3.05x, range=0.11-39.01x; failed QC N=29, mean=0.23x, median=0.04x, range=0.00x-1.91x. Vilnius, passed QC N=57, mean=6.14x, median=5.79x, range=0.11-13.42x; failed QC N=1, DoC=0.03x.

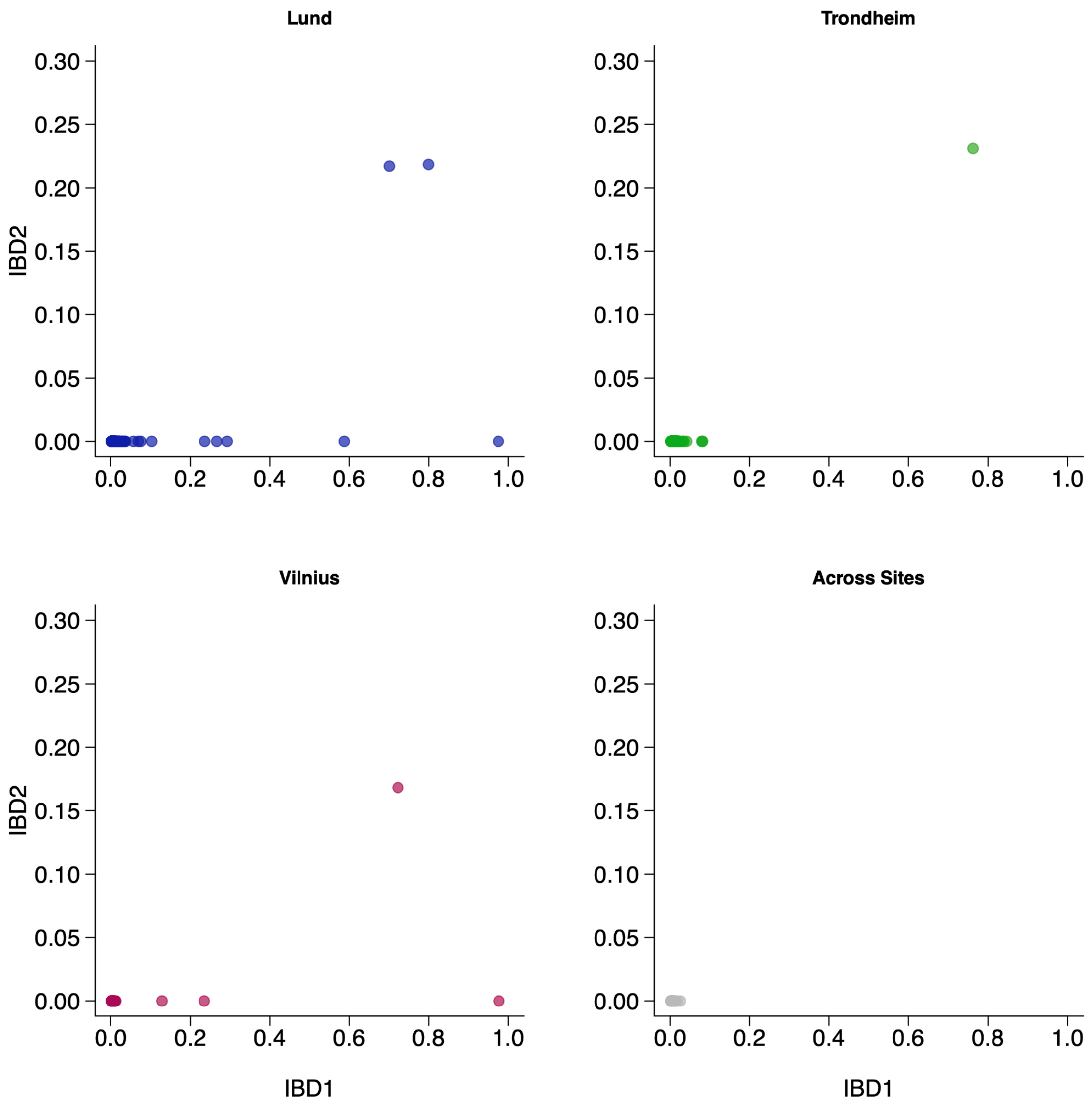


**Supplementary Figure 3.** Relatedness among ancient individuals within Lund (**A**), Trondheim (**B**), and Vilnius (**C**), and across sites (**D**), based on estimates of IBD1 (proportion of the genome where a given pair shares one allele IBD) and IBD2 (proportion of the genome where a given pair shares two alleles IBD).

**A**

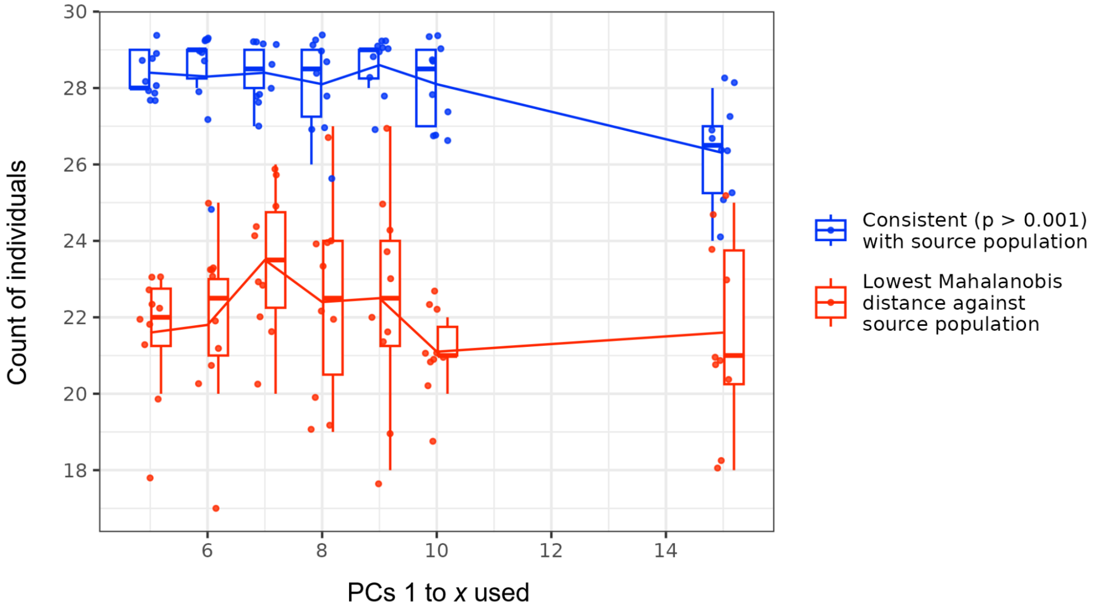


**B**

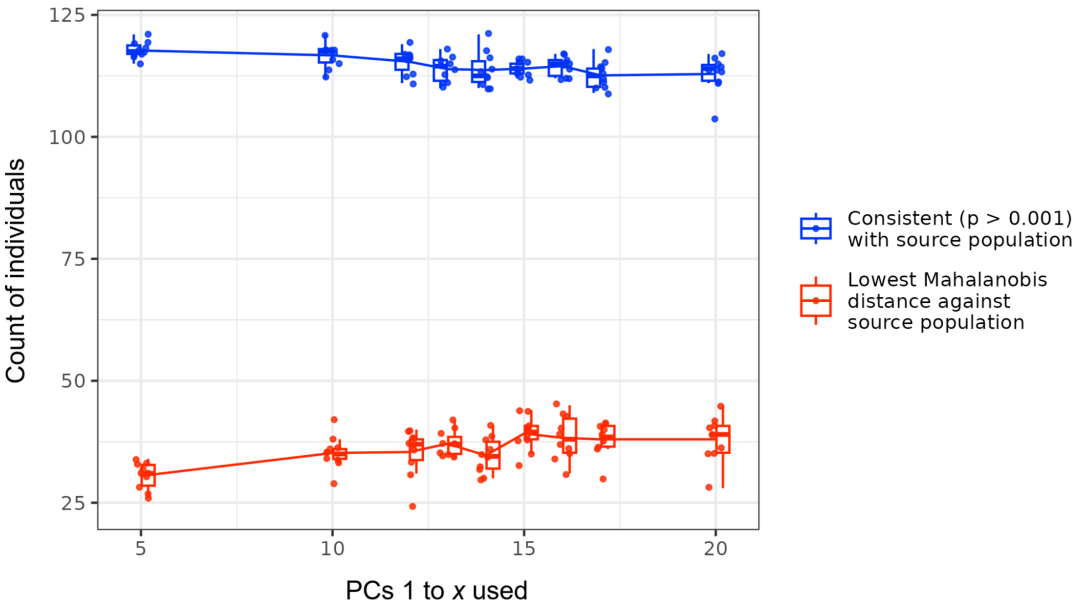


**Supplementary Figure 4.** Hold-out analysis of present-day reference samples informing the selection of PCs in the outlier detection procedure. *Y*-axis represents the count of individuals who are statistically consistent with their source population at *P*<0.001, often non-exclusively (blue), or which show the lowest Mahalanobis distance ($D_M$) with their source population specifically (red). Points represent individual replicates with jitter, boxplots show quartiles and range for replicates, with lines indicating mean value trend. **A**) Results for the West Eurasian PCA (29 individuals per replicate). **B**) Results for the Scandinavian PCA (150 individuals per replicate).

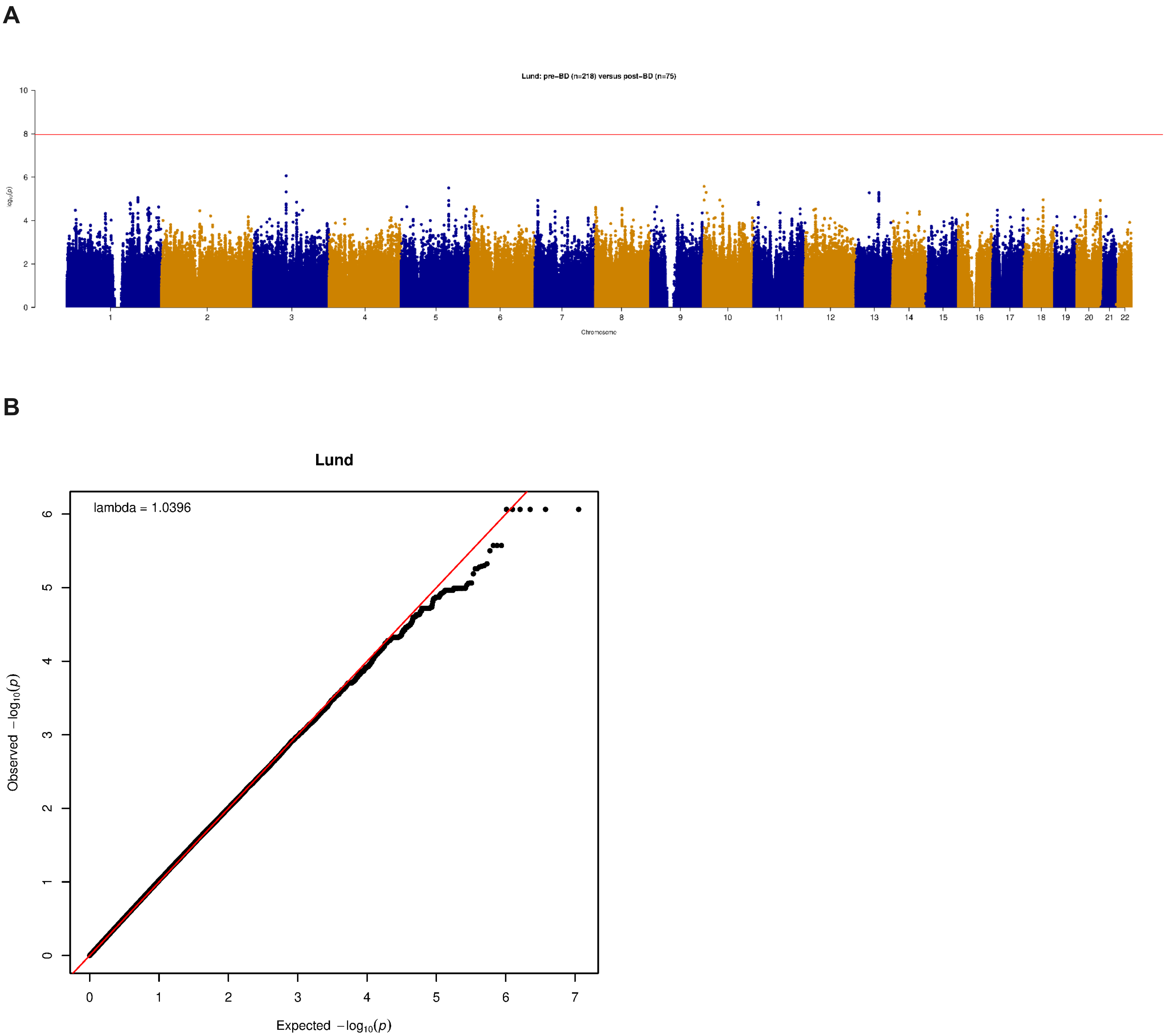


**Supplementary Figure 5. A**) Manhattan plot and **B**) Quantile-quantile (QQ) plot from a genome-wide scan comparing allele frequencies in pre-BD and post-BD individuals in Lund.

**A**

**B**

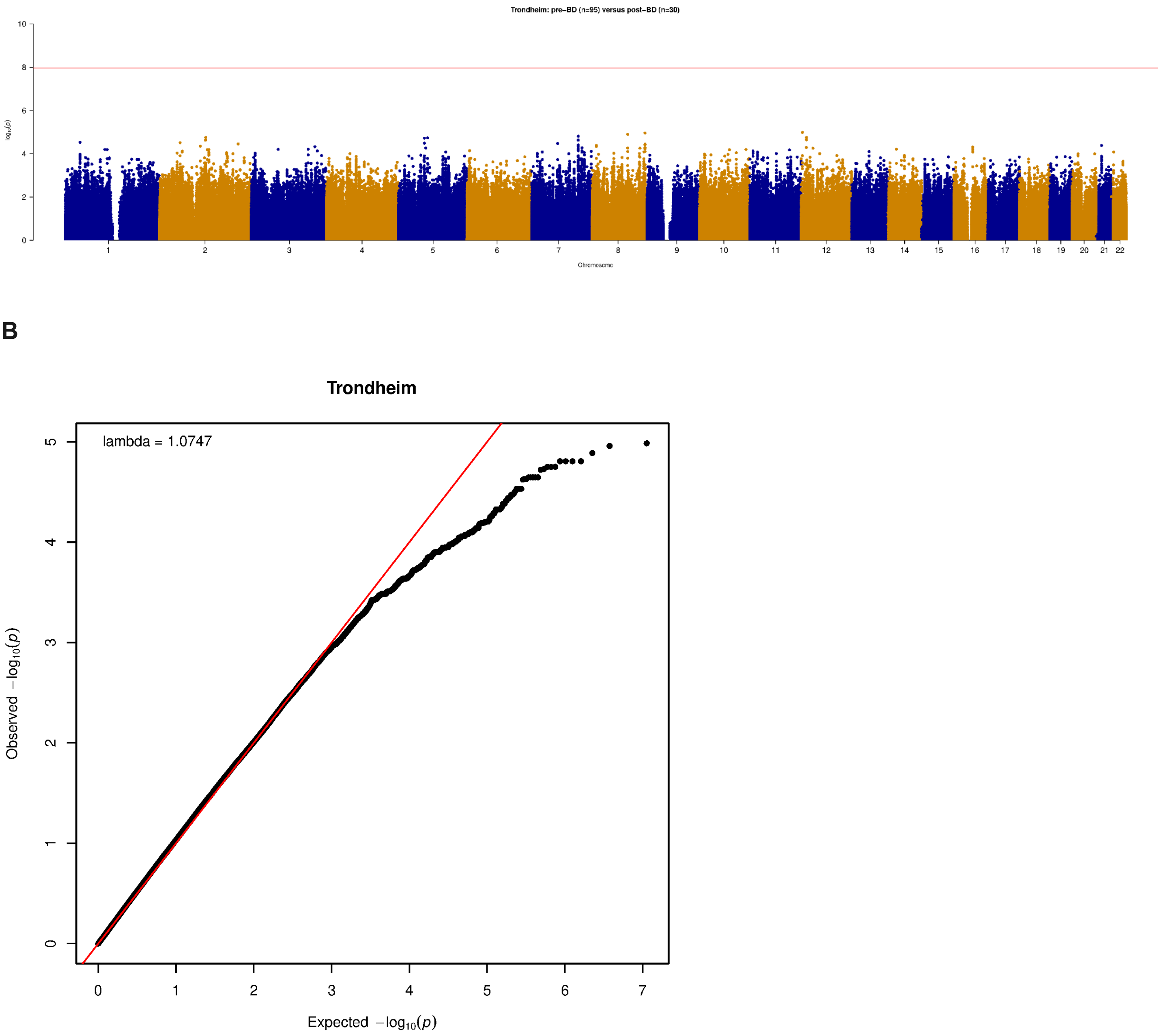


**Supplementary Figure 6. A**) Manhattan plot and **B**) Quantile-quantile (QQ) plot from a genome-wide scan comparing allele frequencies in pre-BD and post-BD individuals from Trondheim.

A

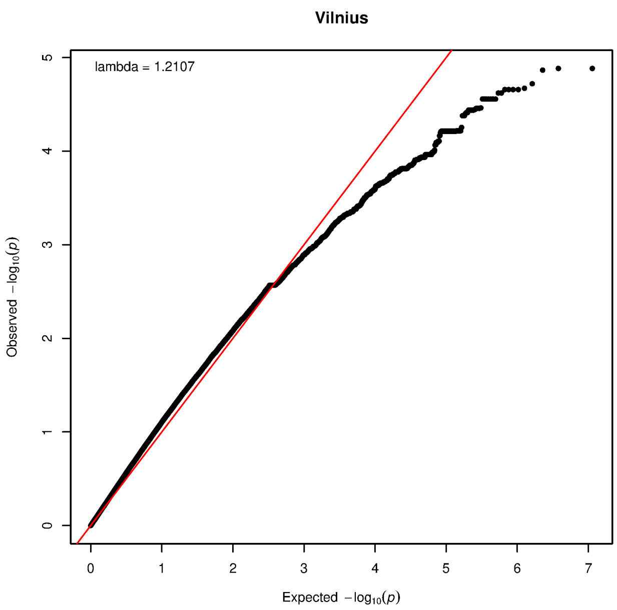


B

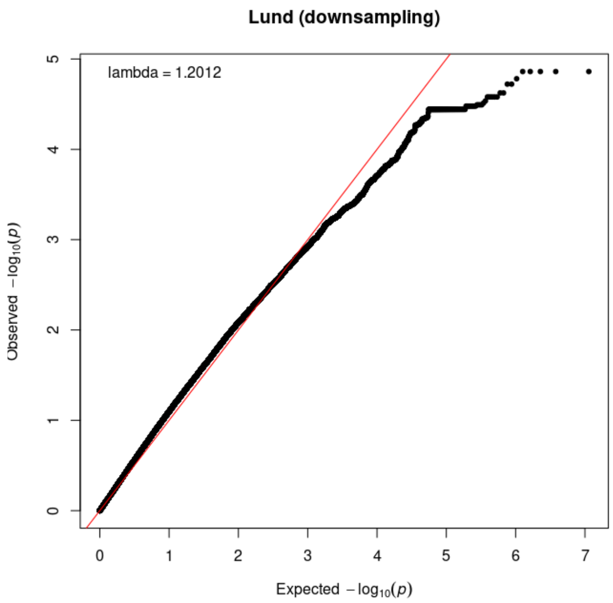


**Supplementary Figure 7. A)** Quantile-quantile (QQ) plot from a genome-wide scan comparing allele frequencies in pre-BD (N=19) and post-BD (N=38) samples from Vilnius, which shows significant genomic inflation (lambda=1.2107) and also deflation in the right tail. **B)** QQ plot from a genome-wide scan comparing Lund samples, downsampled to match the Vilnius cohort size ($N_{pre}$=19 and $N_{post}$=38), The similar pattern of inflation and deflation suggests the poor model fit is largely driven by small sample size.

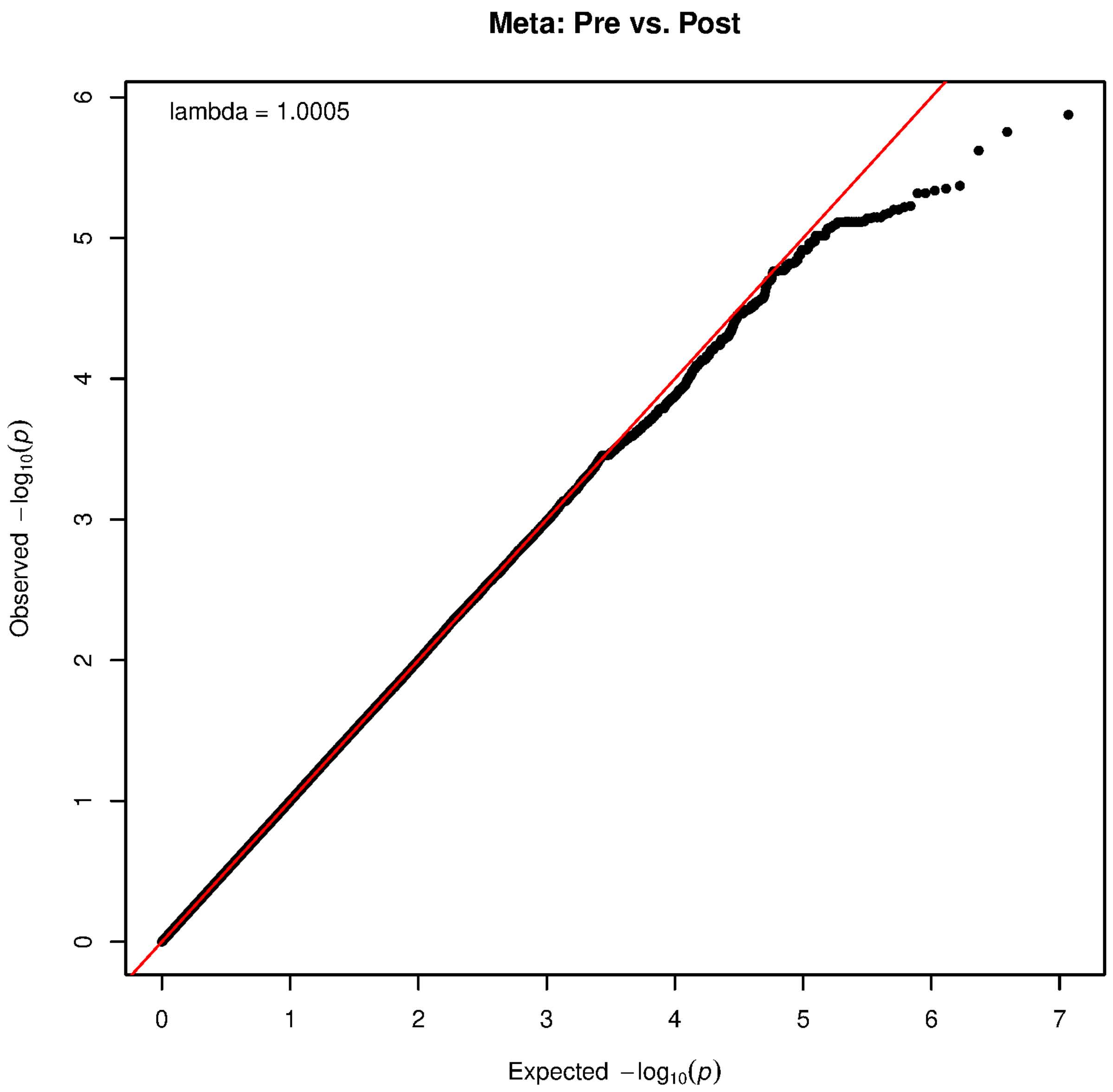


**Supplementary Figure 8.** Quantile-quantile (QQ) plot for the meta-analysis of allele frequency comparison in pre-BD versus post-BD individuals, combining the Lund and Trondheim cohorts.

**A**

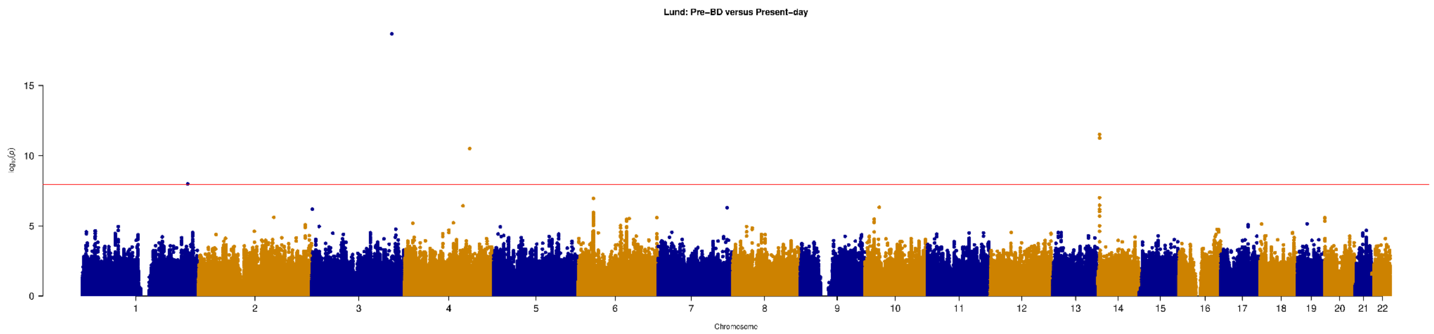


**B**

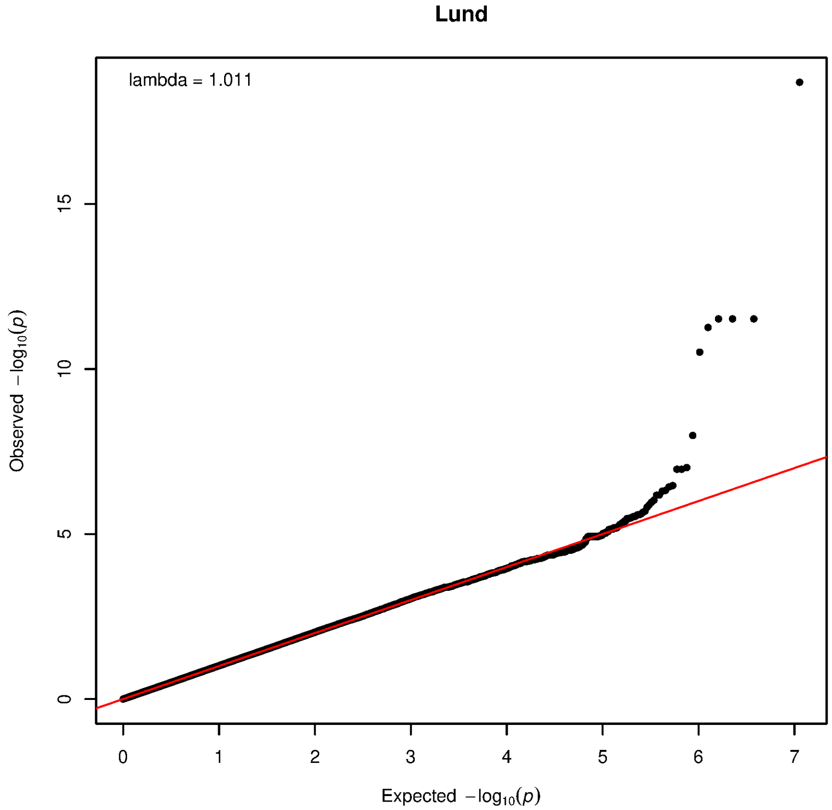


**Supplementary Figure 9. A**) Manhattan plot and **B**) Quantile-quantile (QQ) plot from a genome-wide scan comparing allele frequencies in the pre-BD individuals from Lund with present-day proxies individuals sampled from Denmark and Skåne of Sweden.

A

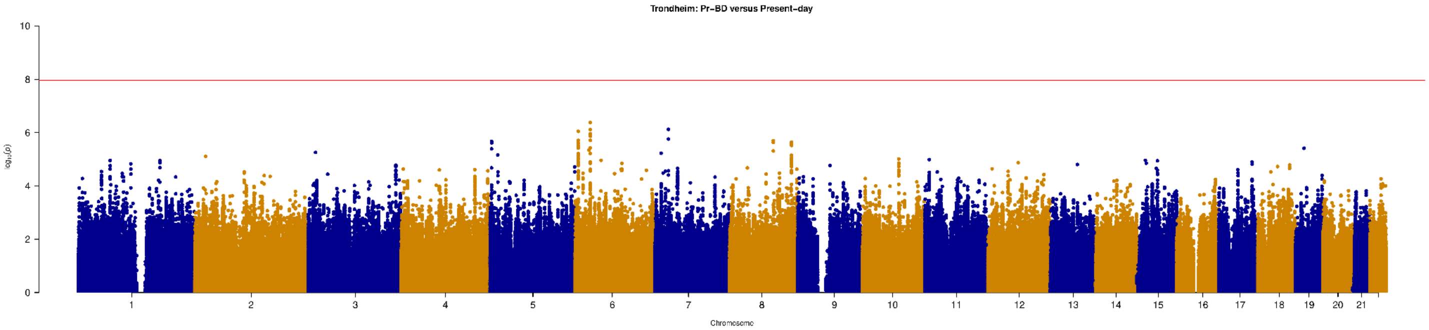


B

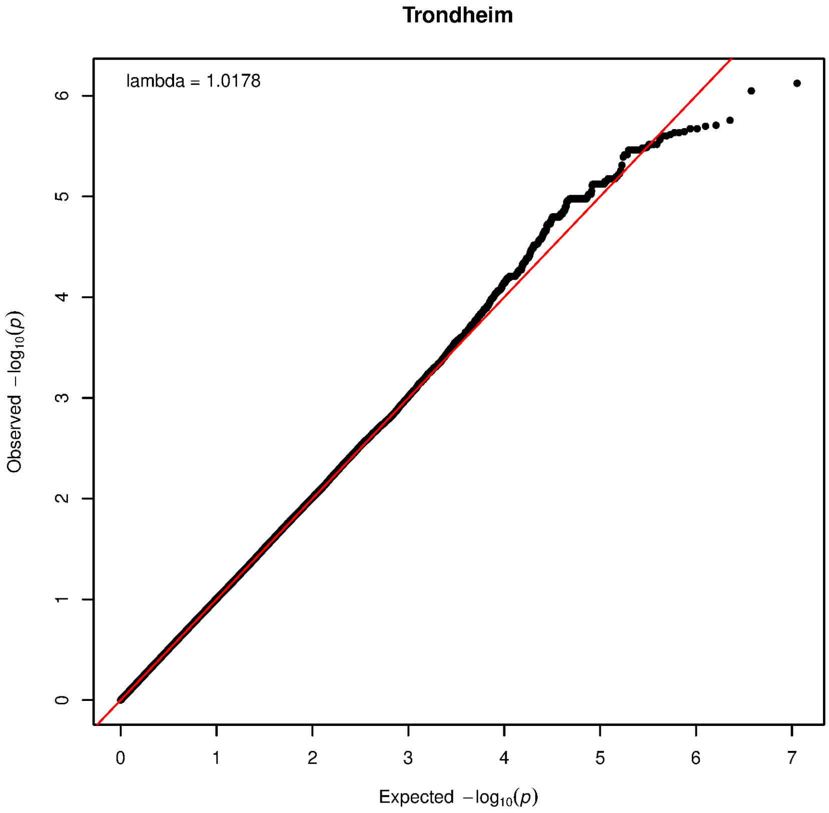


**Supplementary Figure 10. A)** Manhattan plot and **B)** Quantile-quantile (QQ) plot from a genome-wide scan comparing allele frequencies in the pre-BD individuals from Trondheim with present-day proxy individuals sampled from Norway.

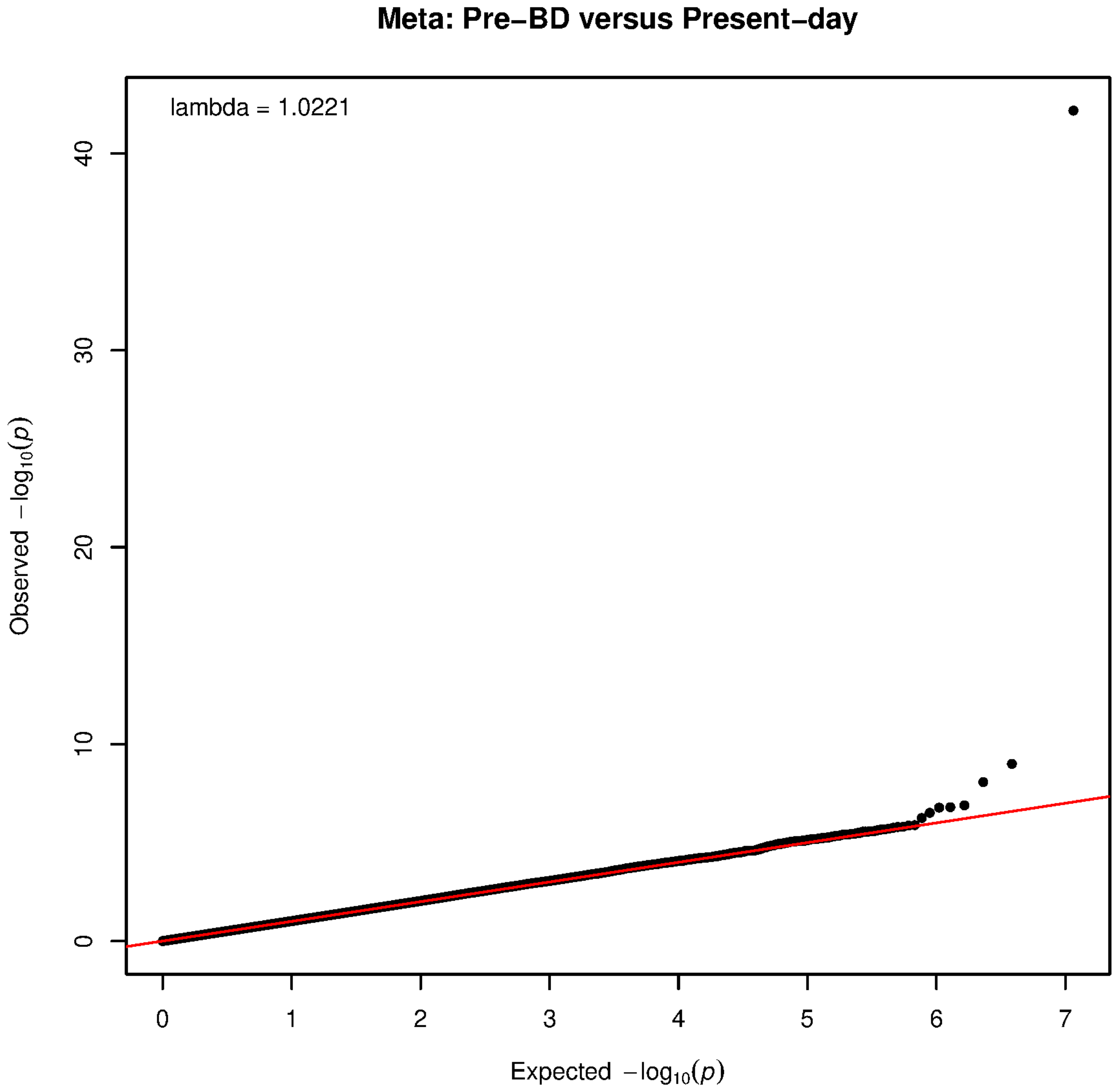


**Supplementary Figure 11.** Quantile-quantile (QQ) plot for the meta-analysis of allele frequencies in pre-BD versus present-day individuals, combining the Lund and Trondheim cohorts.

**A)**

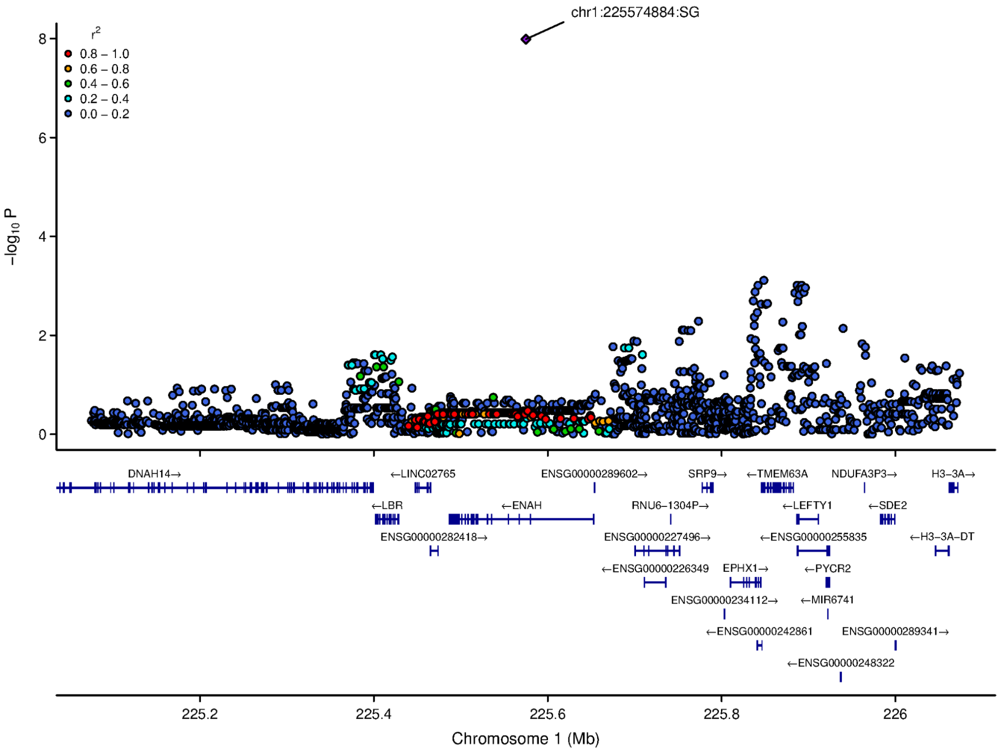

chr1:225574884:SG
r2
0.8 – 1.0
0.6 – 0.8
0.4 – 0.6
0.2 – 0.4
0.0 – 0.2
−log10 P
DNAH14→
←LINC02765
ENSG00000289602→
SRP9→
←TMEM63A
NDUFA3P3→
H3−3A→
←LBR
←ENAH
RNU6−1304P→
←LEFTY1
←SDE2
ENSG00000282418→
ENSG00000227496→
←ENSG00000255835
←H3−3A−DT
←ENSG00000226349
EPHX1→
←PYCR2
ENSG00000234112→
←MIR6741
←ENSG00000242861
ENSG00000289341→
←ENSG00000248322
Chromosome 1 (Mb)


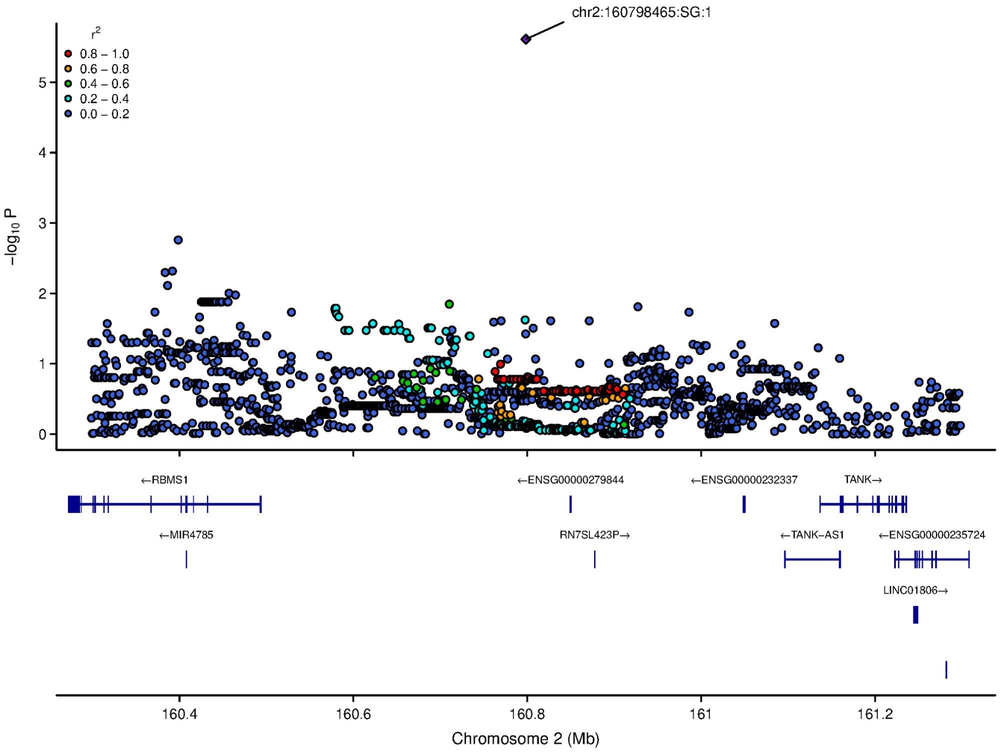

chr2:160798465:SG:1
r2
0.8 – 1.0
0.6 – 0.8
0.4 – 0.6
0.2 – 0.4
0.0 – 0.2
−log10 P
←RBMS1
←ENSG00000279844
←ENSG00000232337
TANK→
←MIR4785
RN7SL423P→
←TANK−AS1
←ENSG00000235724
LINC01806→
Chromosome 2 (Mb)

(continued)

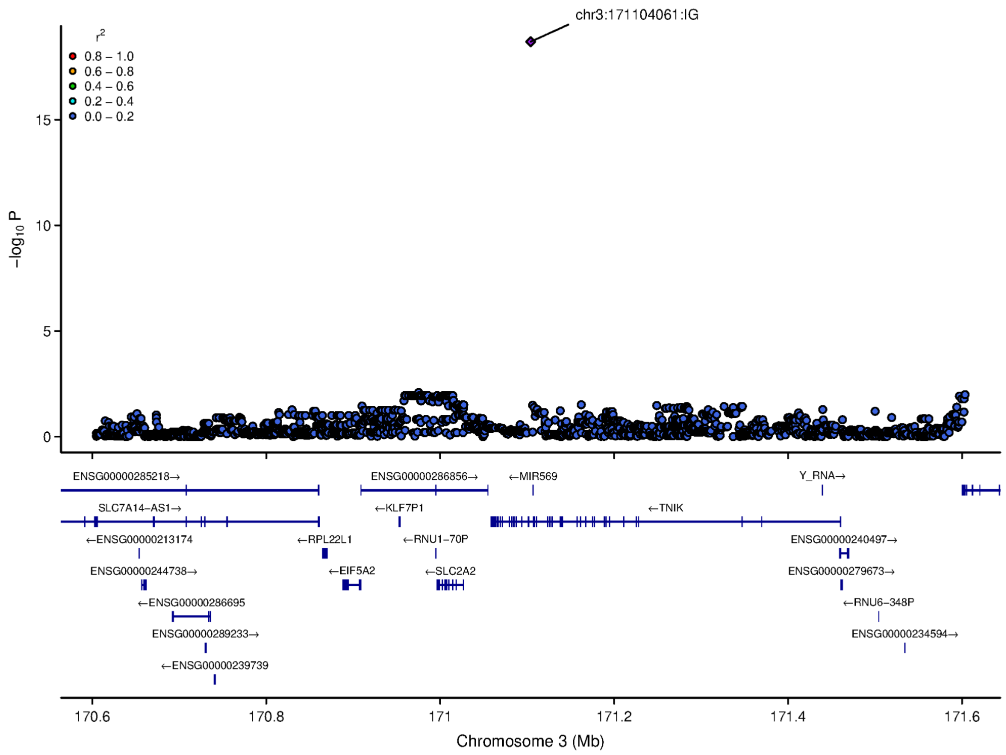
chr3:171104061:IG
r2
0.8 – 1.0
0.6 – 0.8
0.4 – 0.6
0.2 – 0.4
0.0 – 0.2
15
10
5
0
−log10 P
ENSG00000285218→
ENSG00000286856→
←MIR569
Y_RNA→
SLC7A14−AS1→
←KLF7P1
←TNIK
←ENSG00000213174
←RPL22L1
←RNU1−70P
ENSG00000240497→
ENSG00000244738→
←EIF5A2
←SLC2A2
ENSG00000279673→
←ENSG00000286695
←RNU6−348P
ENSG00000289233→
ENSG00000234594→
←ENSG00000239739
170.6
170.8
171
171.2
171.4
171.6
Chromosome 3 (Mb)

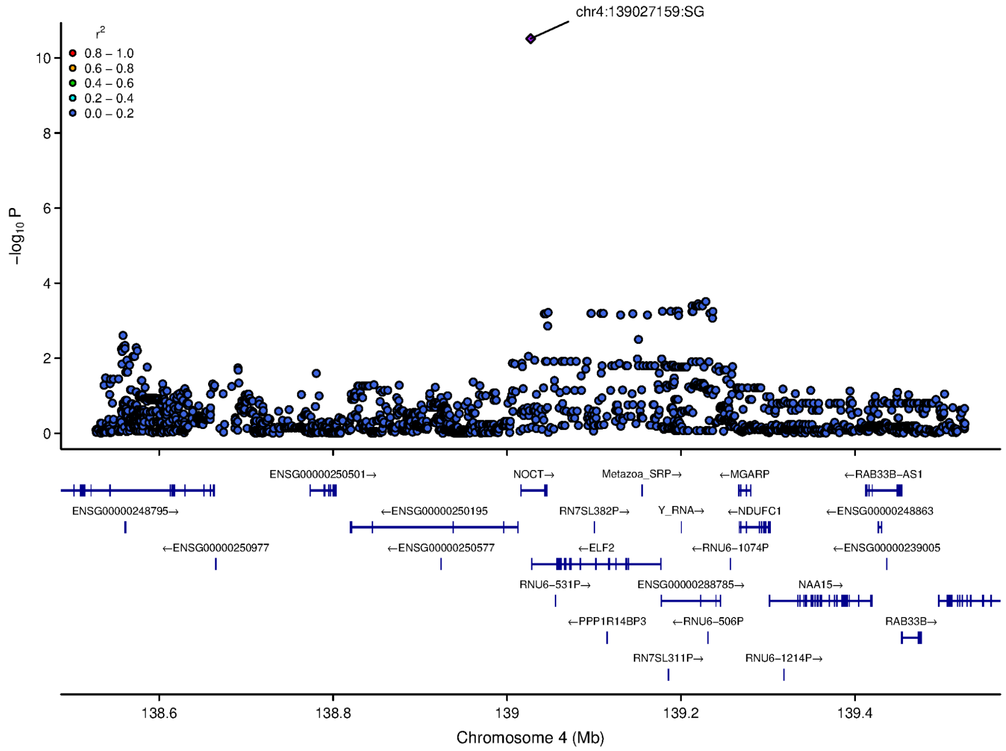
chr4:139027159:SG
r2
0.8 – 1.0
0.6 – 0.8
0.4 – 0.6
0.2 – 0.4
0.0 – 0.2
10
8
6
4
2
0
−log10 P
ENSG00000250501→
NOCT→
Metazoa_SRP→
←MGARP
←RAB33B−AS1
ENSG00000248795→
←ENSG00000250195
RN7SL382P→
Y_RNA→
←NDUFC1
←ENSG00000248863
←ENSG00000250977
←ENSG00000250577
←ELF2
←RNU6−1074P
←ENSG00000239005
RNU6−531P→
ENSG00000288785→
NAA15→
←PPP1R14BP3
←RNU6−506P
RAB33B→
RN7SL311P→
RNU6−1214P→
138.6
138.8
139
139.2
139.4
Chromosome 4 (Mb)

(continued)

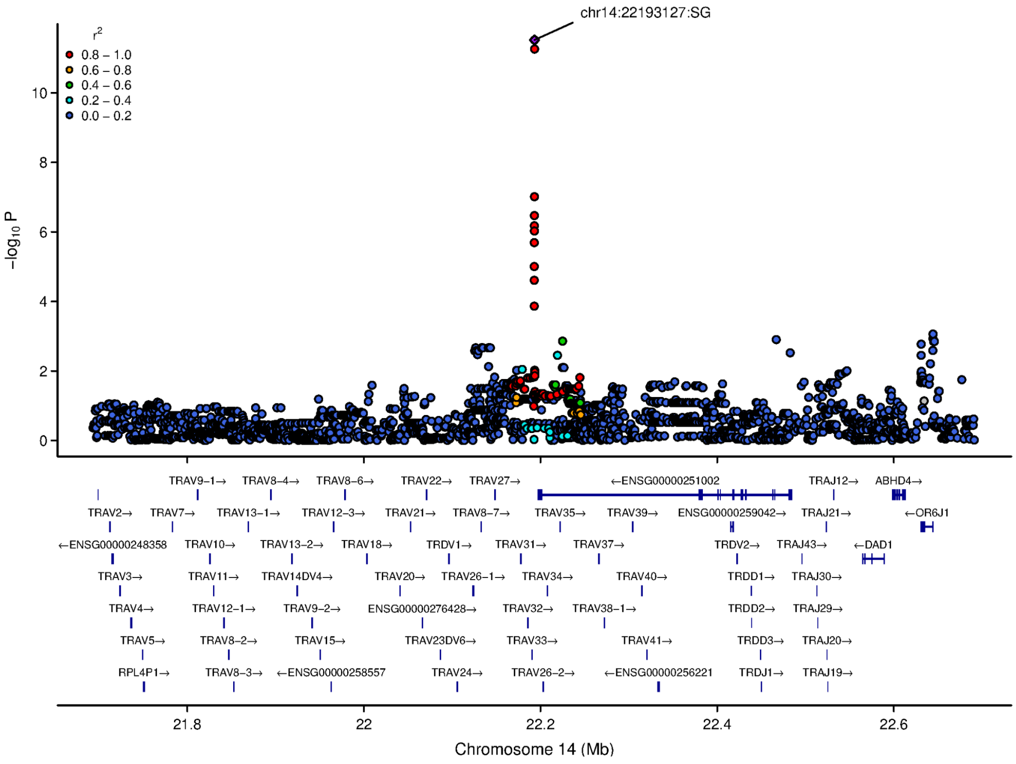
chr14:22193127:SG
r2
0.8 – 1.0
0.6 – 0.8
0.4 – 0.6
0.2 – 0.4
0.0 – 0.2
−log10 P
10
8
6
4
2
0
TRAV9−1→
TRAV8−4→
TRAV8−6→
TRAV22→
TRAV27→
←ENSG00000251002
TRAJ12→
ABHD4→
TRAV2→
TRAV7→
TRAV13−1→
TRAV12−3→
TRAV21→
TRAV8−7→
TRAV35→
TRAV39→
ENSG00000259042→
TRAJ21→
←OR6J1
←ENSG00000248358
TRAV10→
TRAV13−2→
TRAV18→
TRDV1→
TRAV31→
TRAV37→
TRDV2→
TRAJ43→
←DAD1
TRAV3→
TRAV11→
TRAV14DV4→
TRAV20→
TRAV26−1→
TRAV34→
TRAV40→
TRDD1→
TRAJ30→
TRAV4→
TRAV12−1→
TRAV9−2→
ENSG00000276428→
TRAV32→
TRAV38−1→
TRDD2→
TRAJ29→
TRAV5→
TRAV8−2→
TRAV15→
TRAV23DV6→
TRAV33→
TRAV41→
TRDD3→
TRAJ20→
RPL4P1→
TRAV8−3→
←ENSG00000258557
TRAV24→
TRAV26−2→
←ENSG00000256221
TRDJ1→
TRAJ19→
21.8
22
22.2
22.4
22.6
Chromosome 14 (Mb)

**B)**

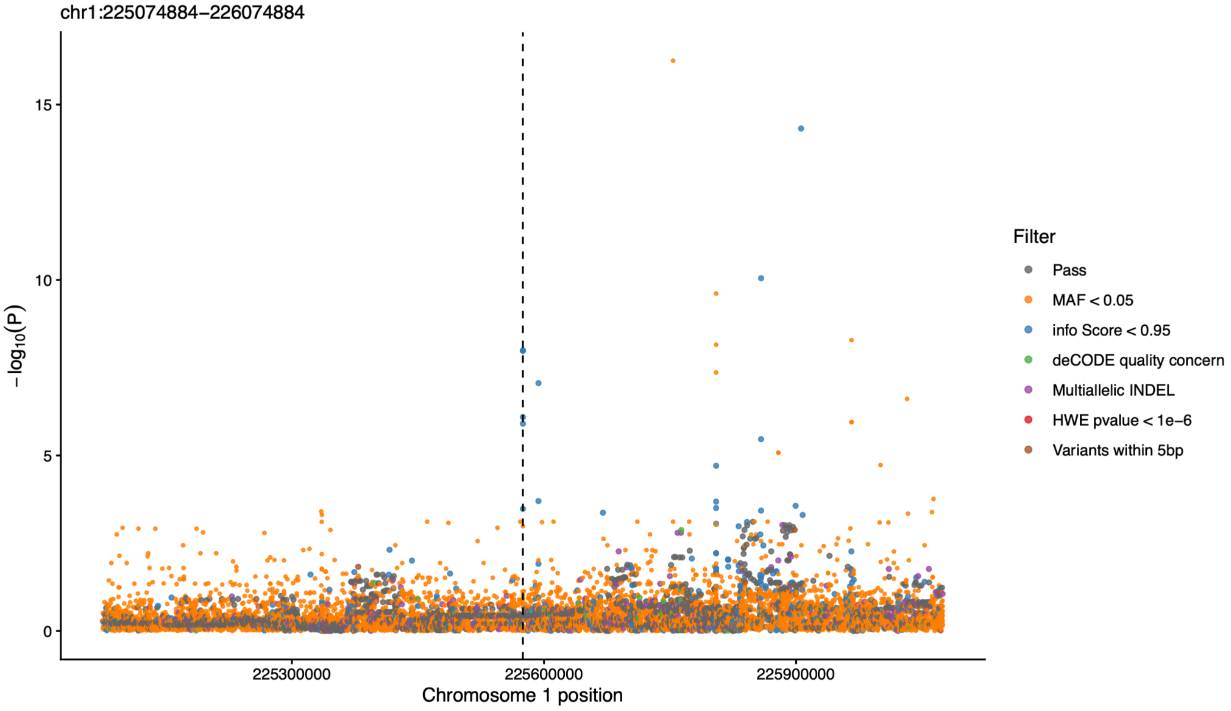
chr1:225074884–226074884
−log10(P)
15
10
5
0
225300000
225600000
225900000
Chromosome 1 position
Filter
Pass
MAF < 0.05
info Score < 0.95
deCODE quality concern
Multiallelic INDEL
HWE pvalue < 1e−6
Variants within 5bp

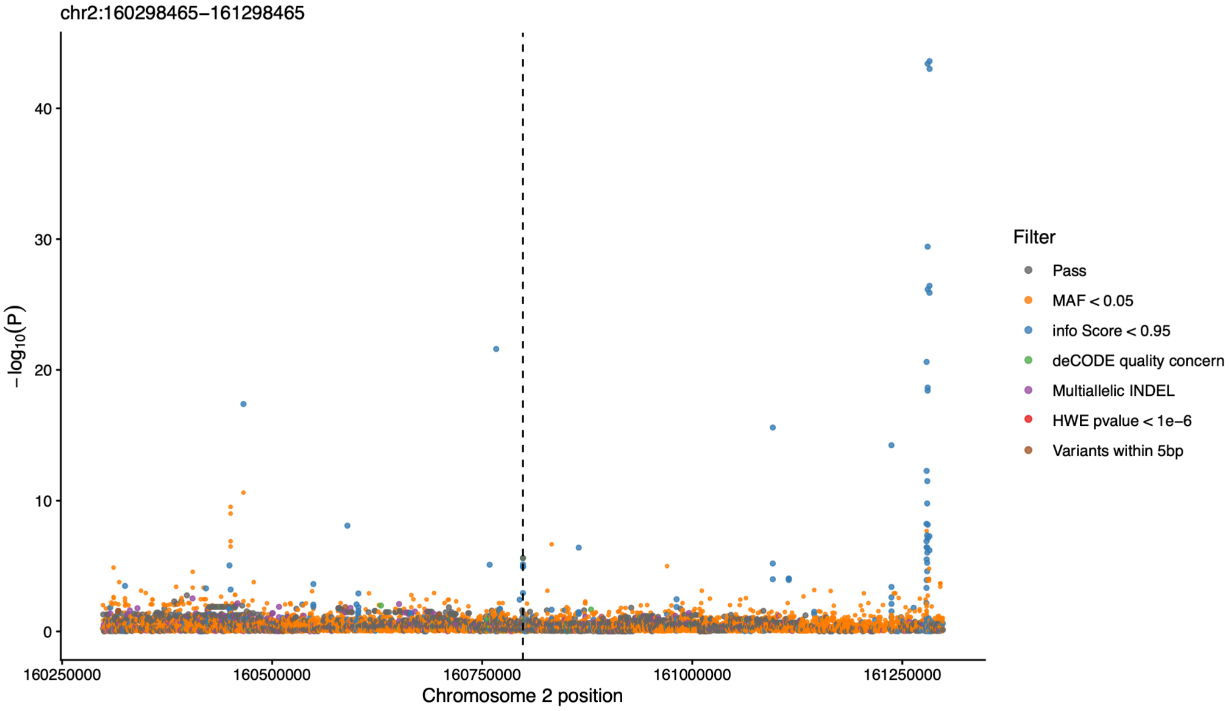
chr2:160298465−161298465
−log10(P)
0
10
20
30
40
160250000
160500000
160750000
161000000
161250000
Chromosome 2 position
Filter
Pass
MAF < 0.05
info Score < 0.95
deCODE quality concern
Multiallelic INDEL
HWE pvalue < 1e−6
Variants within 5bp

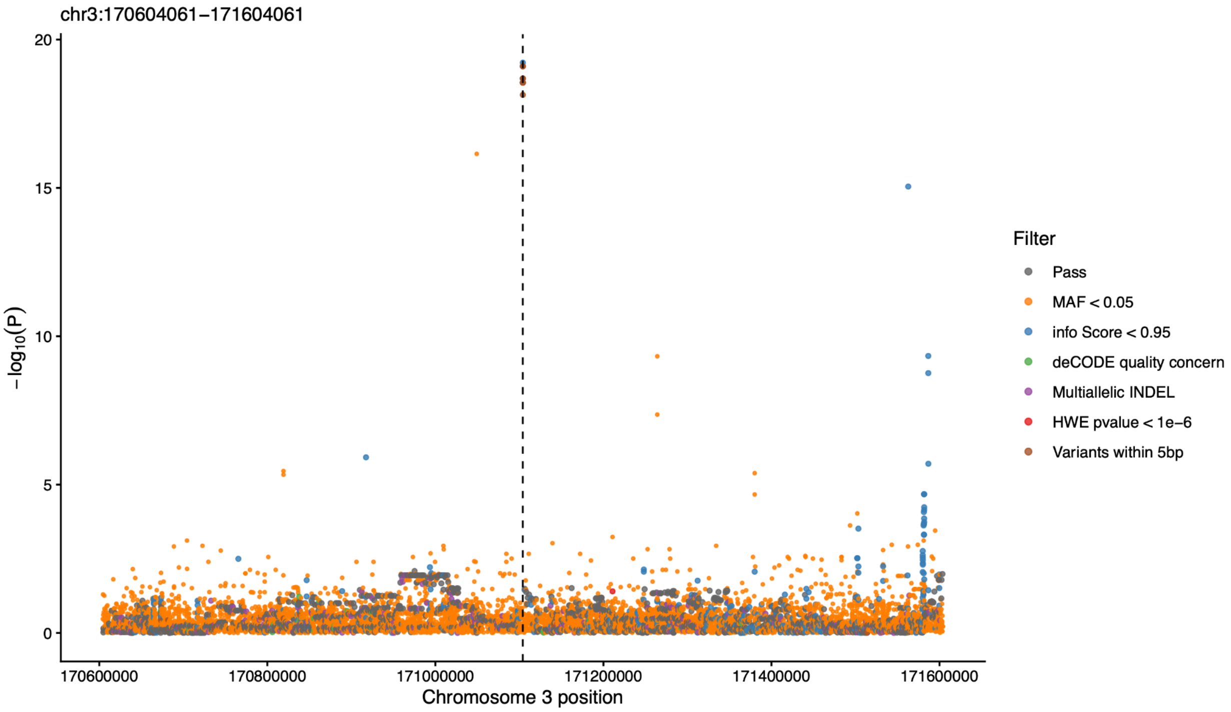
chr3:170604061−171604061
−log10(P)
0
5
10
15
20
170600000
170800000
171000000
171200000
171400000
171600000
Chromosome 3 position
Filter
Pass
MAF < 0.05
info Score < 0.95
deCODE quality concern
Multiallelic INDEL
HWE pvalue < 1e−6
Variants within 5bp

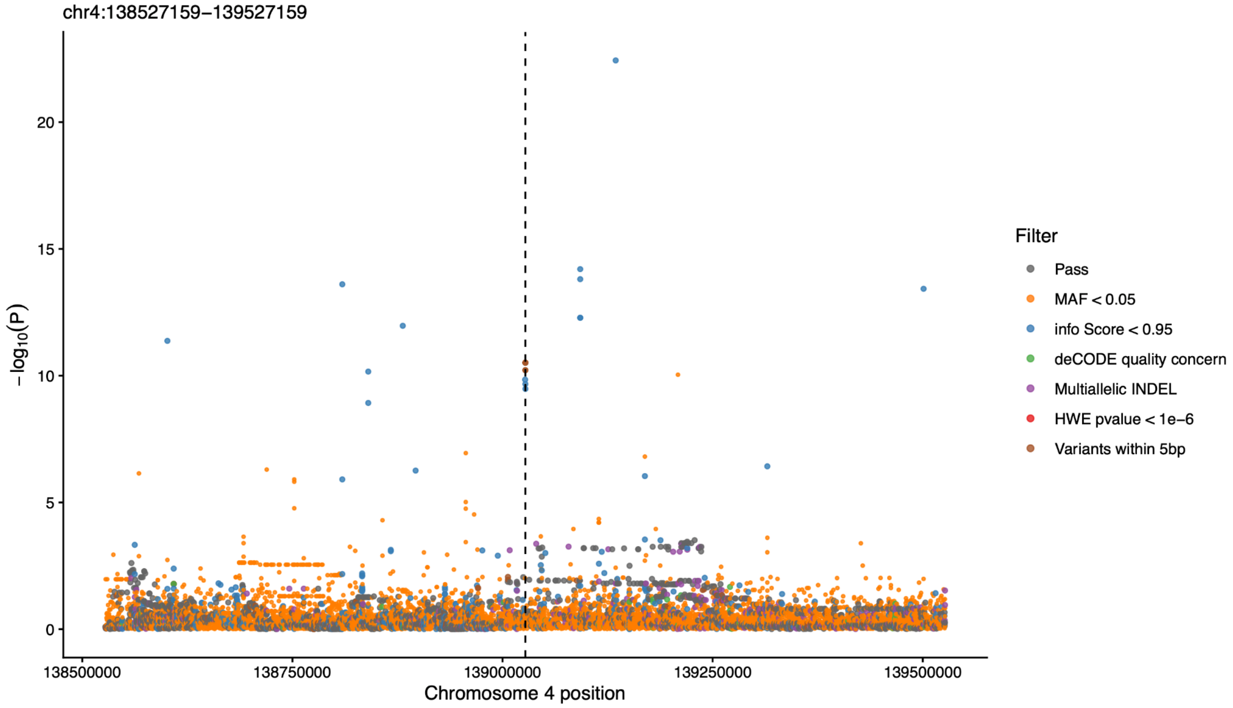


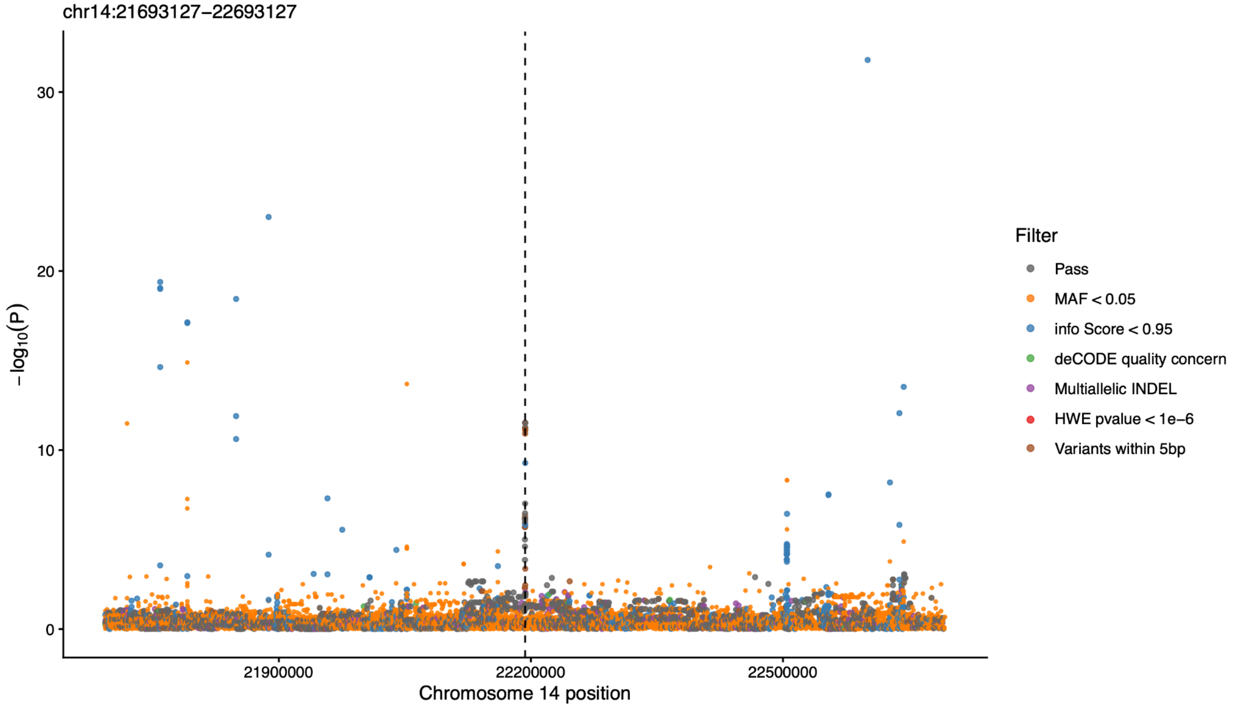


**Supplementary Figure 12.** LocusZoom plots of genome-wide significant loci identified by GWAS (Supplementary Table 6a). **A)** LocusZoom plots after applying stringent site-level filtering (see Methods). Each plot shows an independent locus, with the lead marker indicated for that locus. The x-axis indicates genomic positions, and the y-axis shows the association significance as -log10(*P*). For loci detected only in a site-specific GWAS, the plotted *P*-values are from the corresponding GWAS. For loci that are significant in both site-specific GWAS and the meta-analysis, the plotted *P*-values are from the GWAS comparing allele frequencies between pre-BD individuals from Lund and present-day proxies. Points represent variants in the locus and are colored according to their linkage disequilibrium (LD) with the lead marker. LD was calculated using present-day individuals from Denmark and Skåne (N=380). LD estimates based on present-day individuals from Norway (N=1182) showed highly similar patterns but are not shown. **B)** LocusZoom plots of the same loci before application of the filters. Markers are coloured according to the filter category.

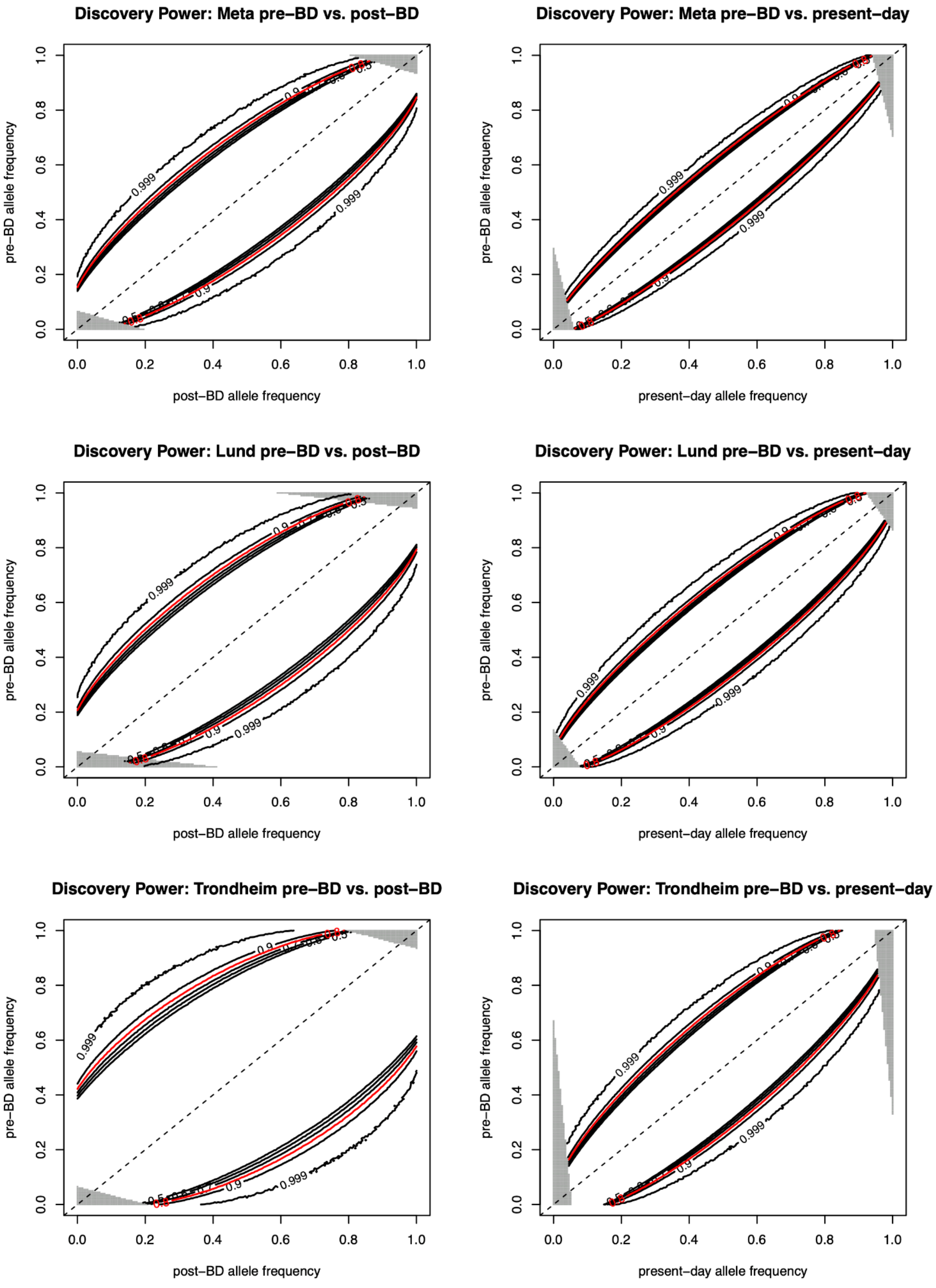


**Supplementary Figure 13.** Power calculations for detecting allele frequency differences at α = $1.1x10^{-8}$. Each panel shows the minimum allele frequency difference required to achieve a given statistical power. Curves represent power levels of 50%, 60%, 70%, 80%, 90% and 99.9%. The red curve indicates 80% power. The left column shows power for pre-BD versus post-BD comparisons, and the right column shows power for pre-BD versus present-day comparisons. Rows show results for the combined meta-analysis, Lund and Trondheim.

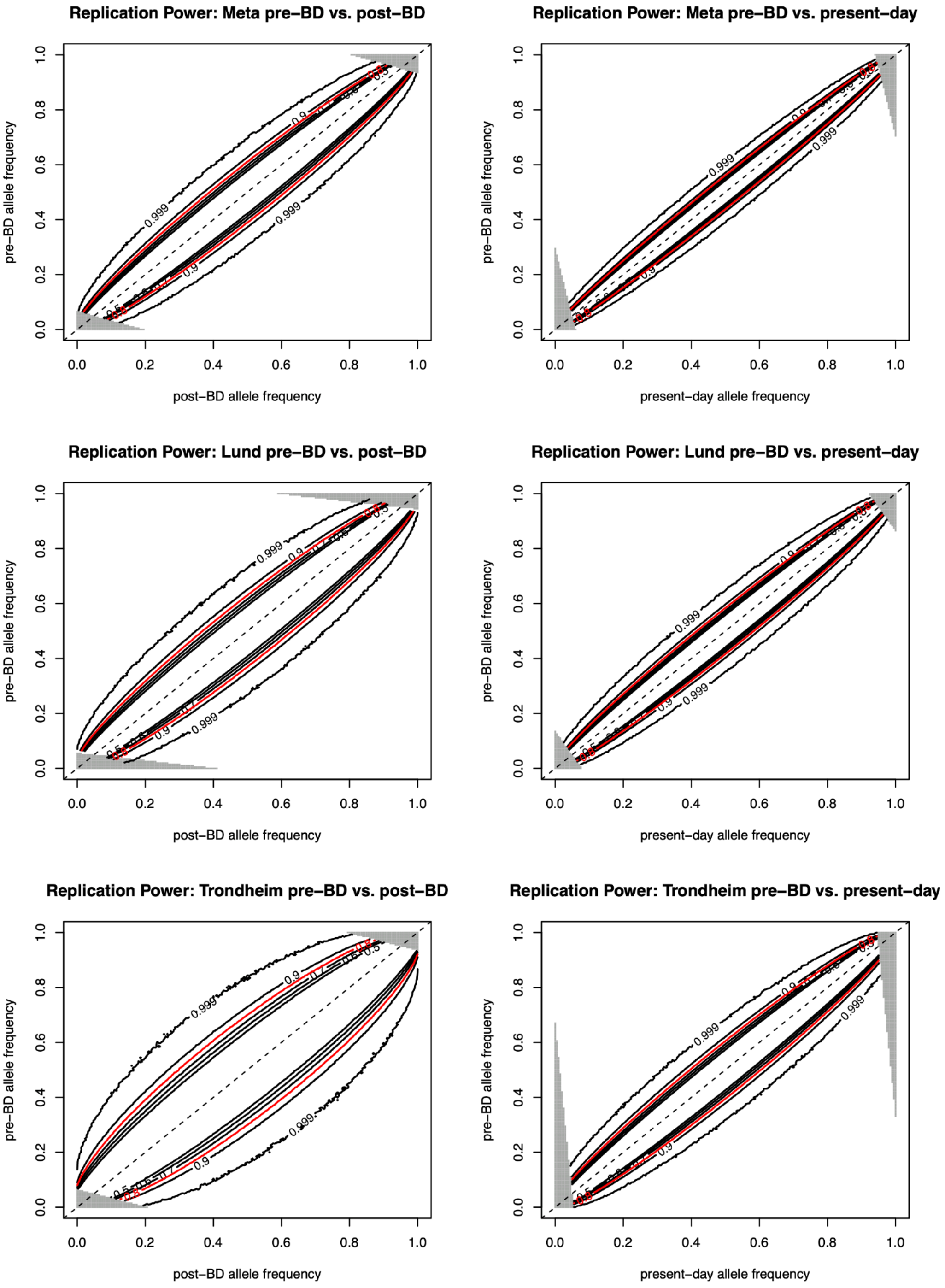


**Supplementary Figure 14.** Power calculations for detecting allele frequency differences at α = 0.05. Each panel shows the minimum allele frequency difference required to achieve a given statistical power. Curves represent power levels of 50%, 60%, 70%, 80%, 90% and 99.9%. The red curve indicates 80% power. The left column shows power for pre-BD versus post-BD comparisons, and the right column shows power for pre-BD versus present-day comparisons. Rows show results for the combined meta-analysis, Lund and Trondheim.

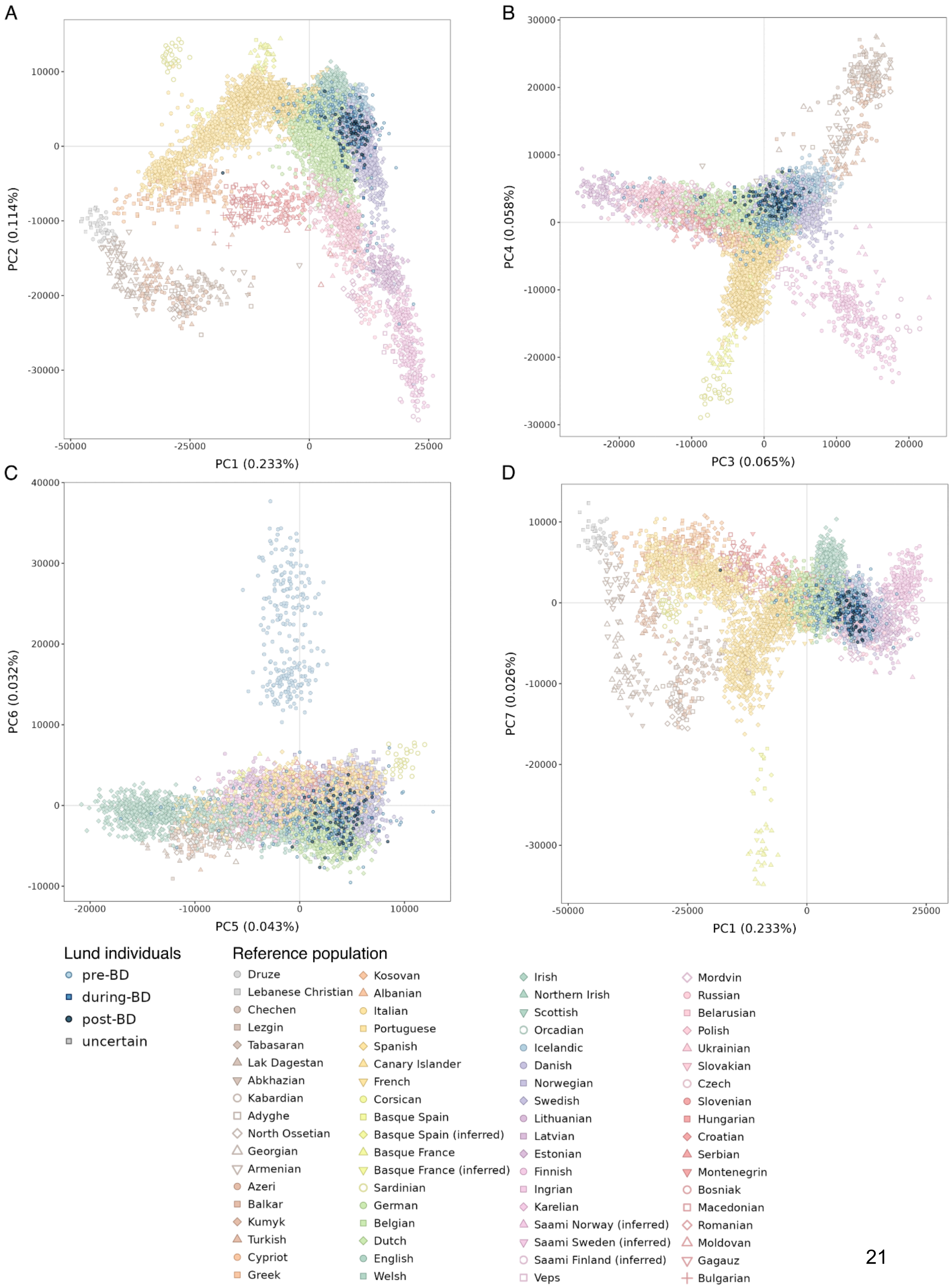
A
B
C
D
PC1 (0.233%)
PC2 (0.114%)
PC3 (0.065%)
PC4 (0.058%)
PC5 (0.043%)
PC6 (0.032%)
PC7 (0.026%)
Lund individuals
pre-BD
during-BD
post-BD
uncertain
Reference population
Druze
Lebanese Christian
Chechen
Lezgin
Tabasaran
Lak Dagestan
Abkhazian
Kabardian
Adyghe
North Ossetian
Georgian
Armenian
Azeri
Balkar
Kumyk
Turkish
Cypriot
Greek
Kosovan
Albanian
Italian
Portuguese
Spanish
Canary Islander
French
Corsican
Basque Spain
Basque Spain (inferred)
Basque France
Basque France (inferred)
Sardinian
German
Belgian
Dutch
English
Welsh
Irish
Northern Irish
Scottish
Orcadian
Icelandic
Danish
Norwegian
Swedish
Lithuanian
Latvian
Estonian
Finnish
Ingrian
Karelian
Saami Norway (inferred)
Saami Sweden (inferred)
Saami Finland (inferred)
Veps
Mordvin
Russian
Belarusian
Polish
Ukrainian
Slovakian
Czech
Slovenian
Hungarian
Croatian
Serbian
Montenegrin
Bosniak
Macedonian
Romanian
Moldovan
Gagauz
Bulgarian

(continued)

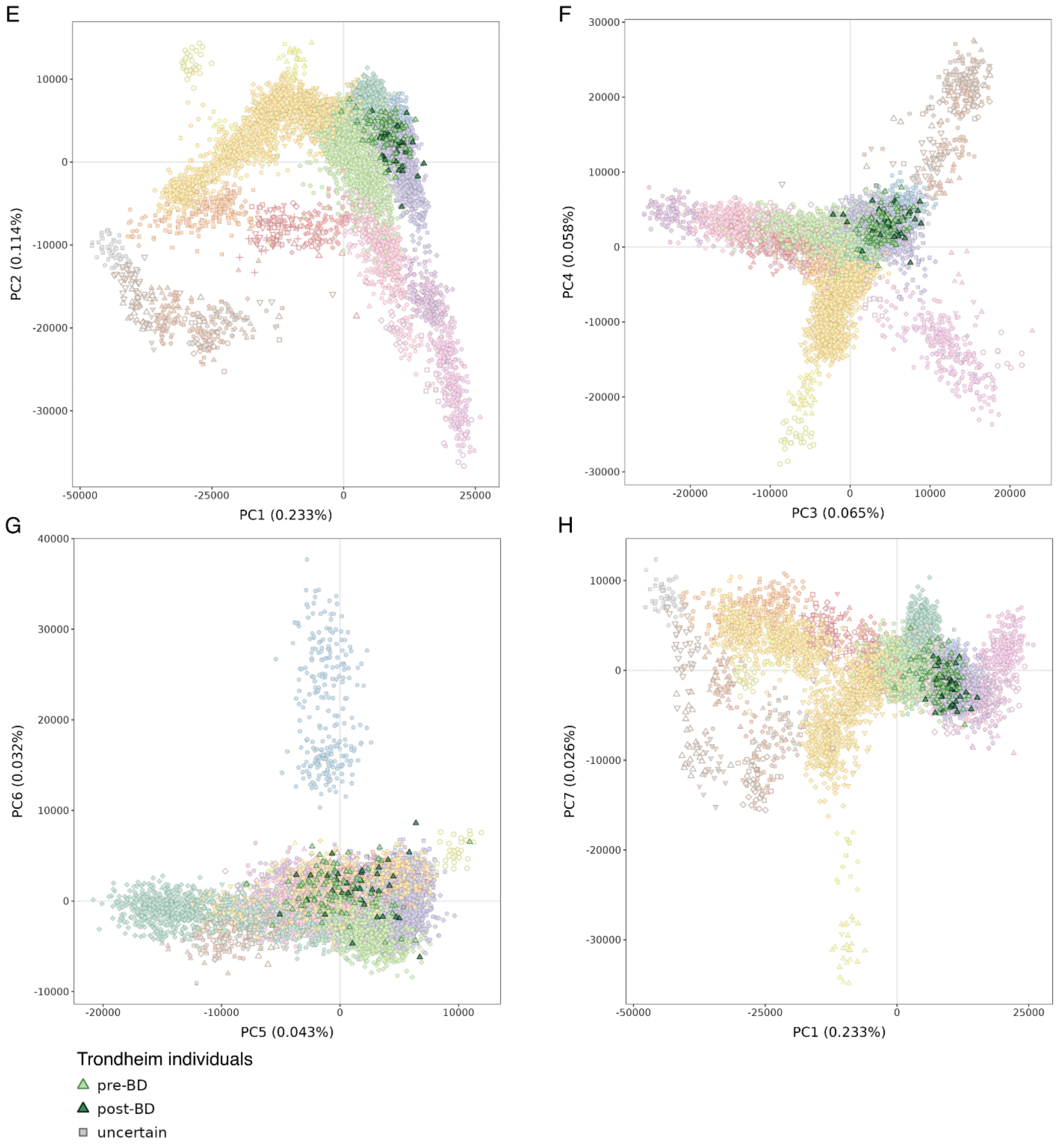
E
F
G
H
PC1 (0.233%)
PC2 (0.114%)
PC3 (0.065%)
PC4 (0.058%)
PC5 (0.043%)
PC6 (0.032%)
PC1 (0.233%)
PC7 (0.026%)
Trondheim individuals
pre-BD
post-BD
uncertain

(continued)

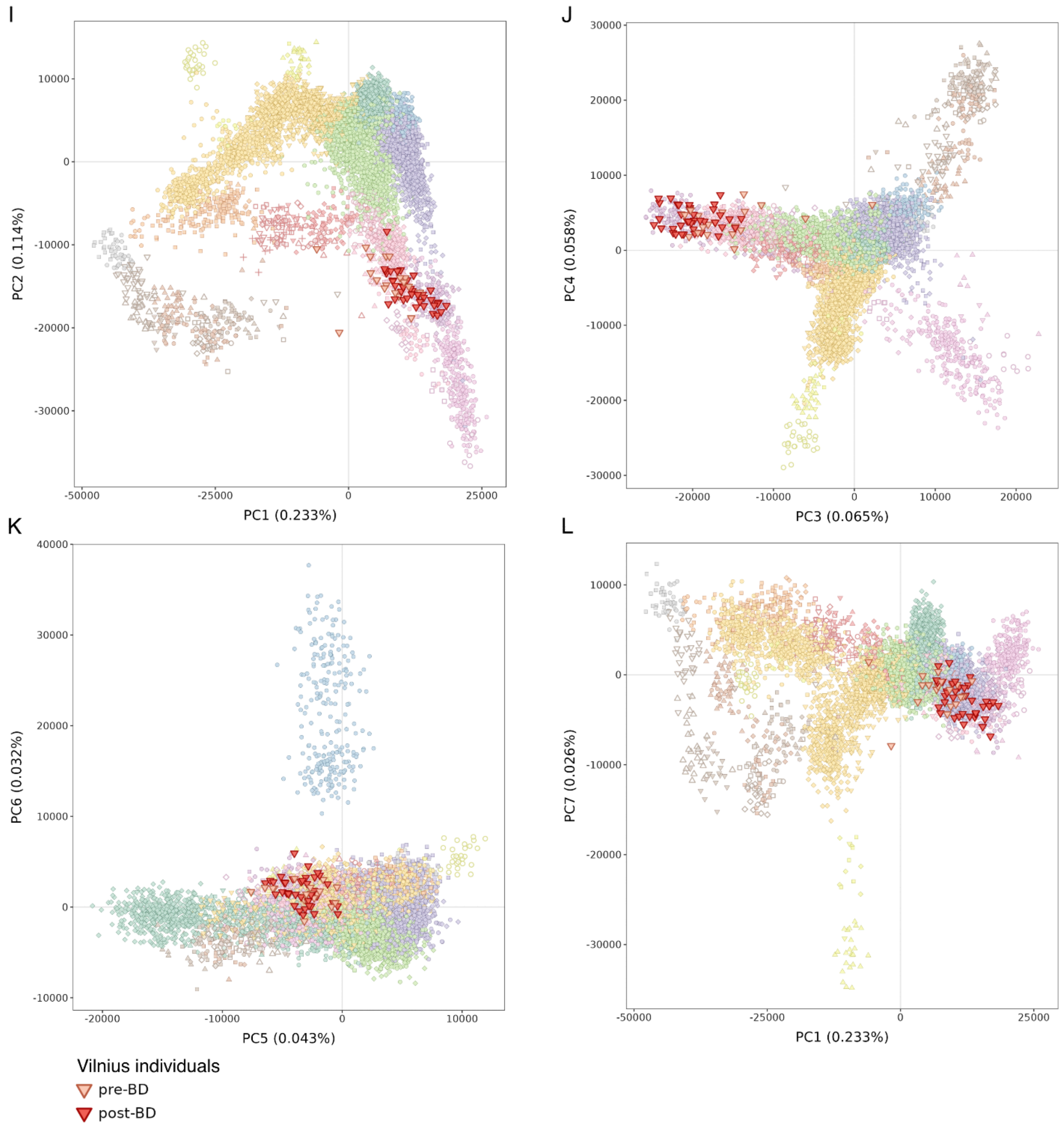


**Supplementary Figure 15.** PCA projection of ancient individuals onto present-day West Eurasian populations. Ancient individuals from Lund **(A-D)**, Trondheim **(E-H)**, and Vilnius **(I-L)** were projected onto a PCA constructed from 67 present-day West Eurasian populations (Supplementary Table 5). Ancient individuals are colored according to temporal group, and present-day reference populations are indicated by combinations of colour and shape.

**A)** Trondheim

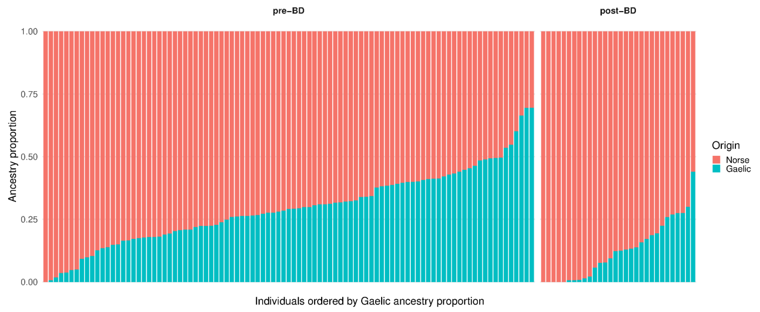


**B)** Lund

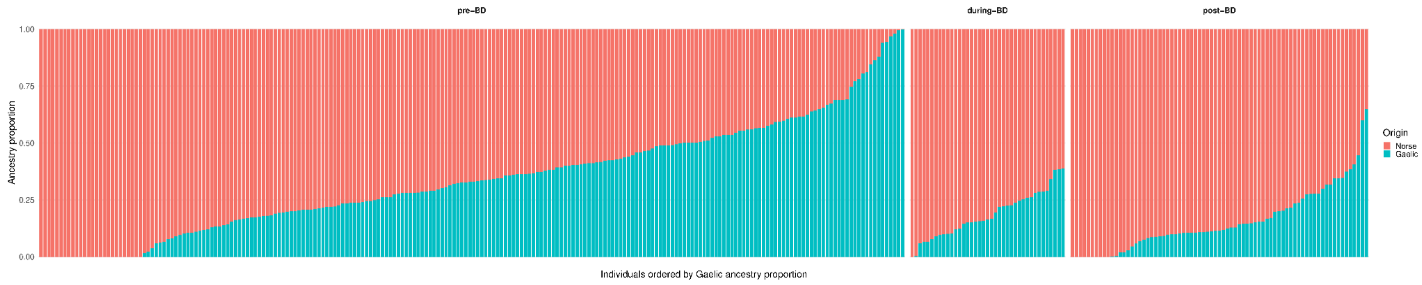


**Supplementary Figure 16.** Supervised ADMIXTURE analysis using present-day reference populations representative of Gaelic and Norse ancestry. ADMIXTURE proportions are shown for individuals from (**A**) Trondheim and (**B**) Lund using Gaelic and Norse as potential source populations (Supplementary Table 5). Individuals are ordered by their inferred Gaelic ancestry proportion within each site.

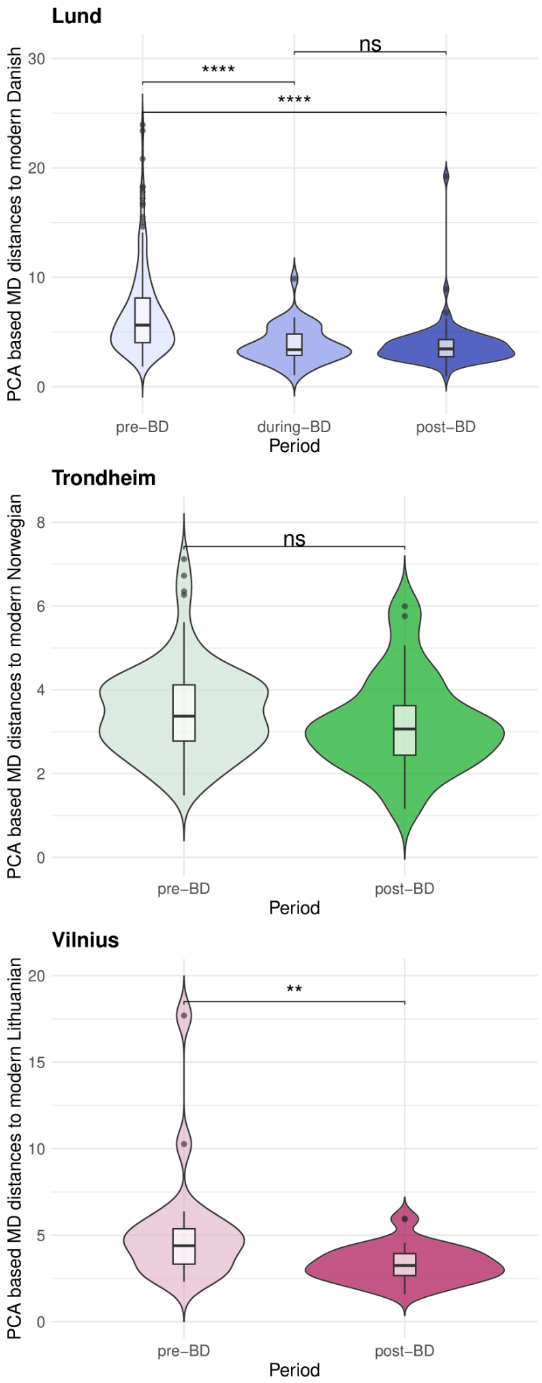


**Supplementary Figure 17.** Violin plots of Mahalanobis distances ($D_M$) across temporal groups. $D_M$ was calculated relative to the present-day reference population indicated on the y-axis using the first 7 PCs from the West Eurasian PCA projection, and compared among temporal groups for Lund (top), Trondheim (middle), and Vilnius (bottom). Statistical significance was assessed using Wilcoxon rank sum tests. Significance codes indicate *P*-value thresholds as follows: **** *P*≤0.0001, *** *P*≤0.001, ** *P*≤0.01, * *P*≤0.05, . *P*≤0.1, and *ns* >0.1.

**A)** IBD profiles for individuals classified as non-Scandinavian outliers in Lund

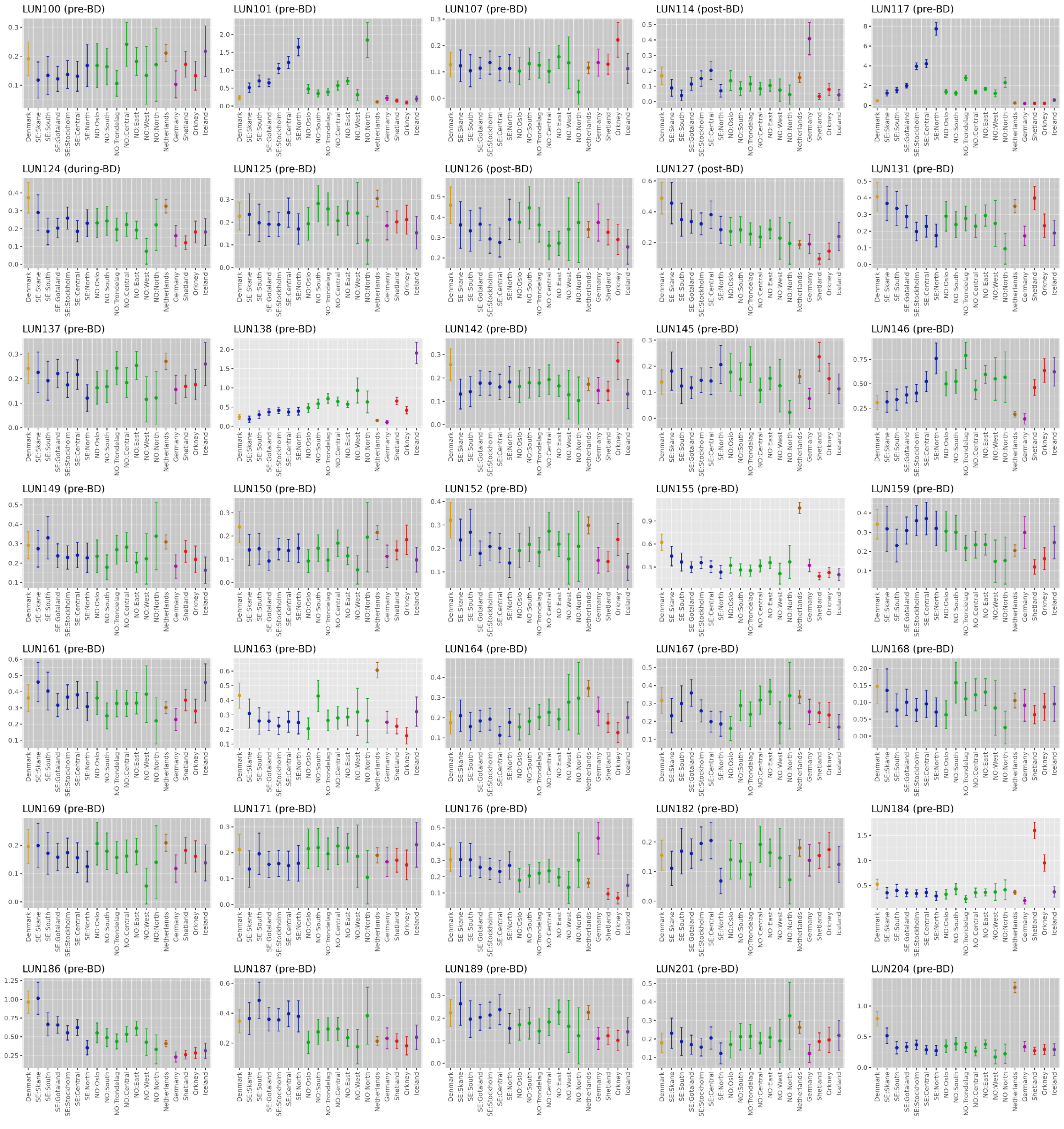

(continued)

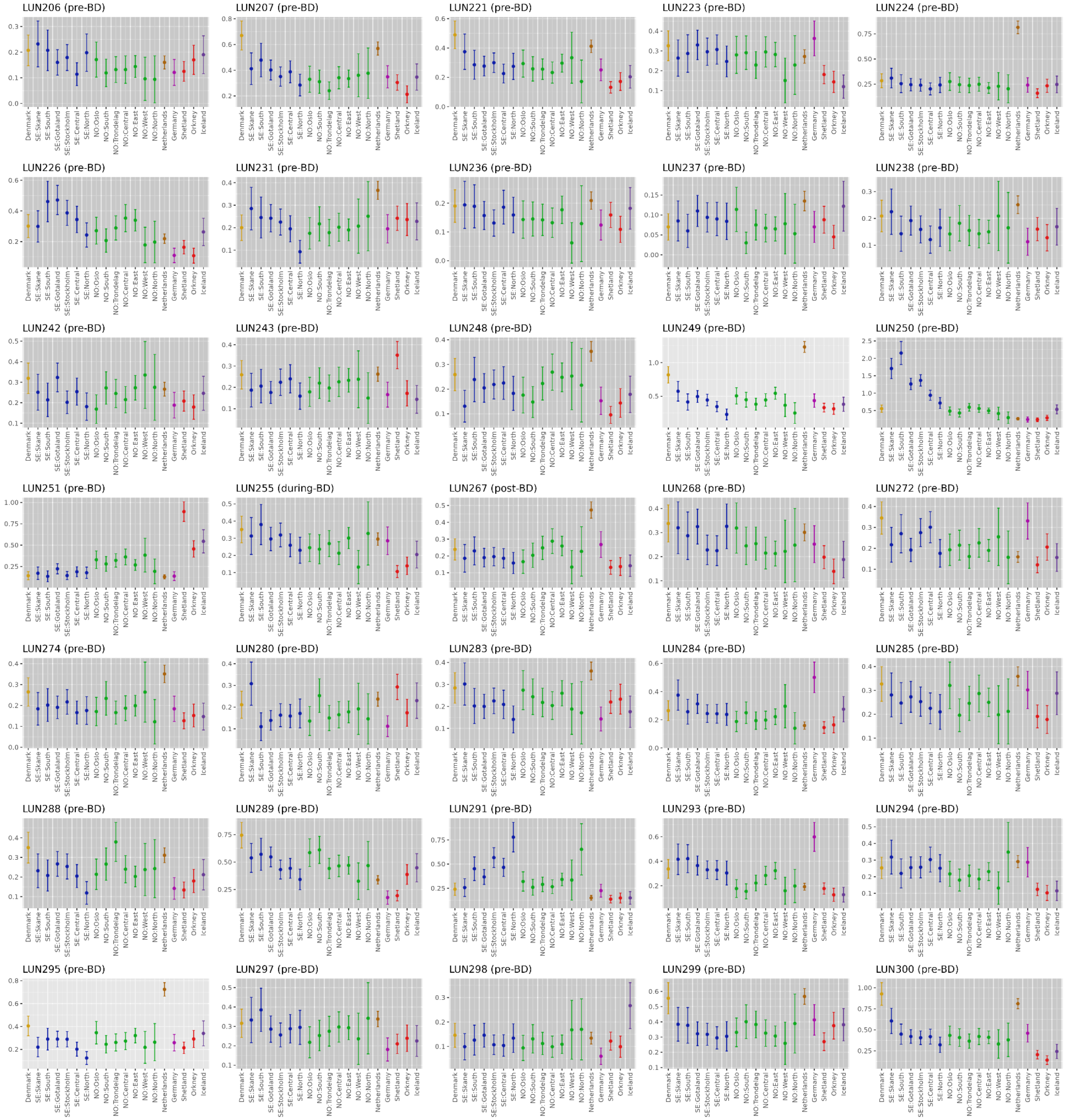
LUN206 (pre-BD)
LUN207 (pre-BD)
LUN221 (pre-BD)
LUN223 (pre-BD)
LUN224 (pre-BD)
LUN226 (pre-BD)
LUN231 (pre-BD)
LUN236 (pre-BD)
LUN237 (pre-BD)
LUN238 (pre-BD)
LUN242 (pre-BD)
LUN243 (pre-BD)
LUN248 (pre-BD)
LUN249 (pre-BD)
LUN250 (pre-BD)
LUN251 (pre-BD)
LUN255 (during-BD)
LUN267 (post-BD)
LUN268 (pre-BD)
LUN272 (pre-BD)
LUN274 (pre-BD)
LUN280 (pre-BD)
LUN283 (pre-BD)
LUN284 (pre-BD)
LUN285 (pre-BD)
LUN288 (pre-BD)
LUN289 (pre-BD)
LUN291 (pre-BD)
LUN293 (pre-BD)
LUN294 (pre-BD)
LUN295 (pre-BD)
LUN297 (pre-BD)
LUN298 (pre-BD)
LUN299 (pre-BD)
LUN300 (pre-BD)
Denmark
SE:Skane
SE:South
SE:Gotaland
SE:Stockholm
SE:Central
SE:North
NO:Oslo
NO:South
NO:Trondelag
NO:Central
NO:East
NO:West
NO:North
Netherlands
Germany
Shetland
Orkney
Iceland

(continued)

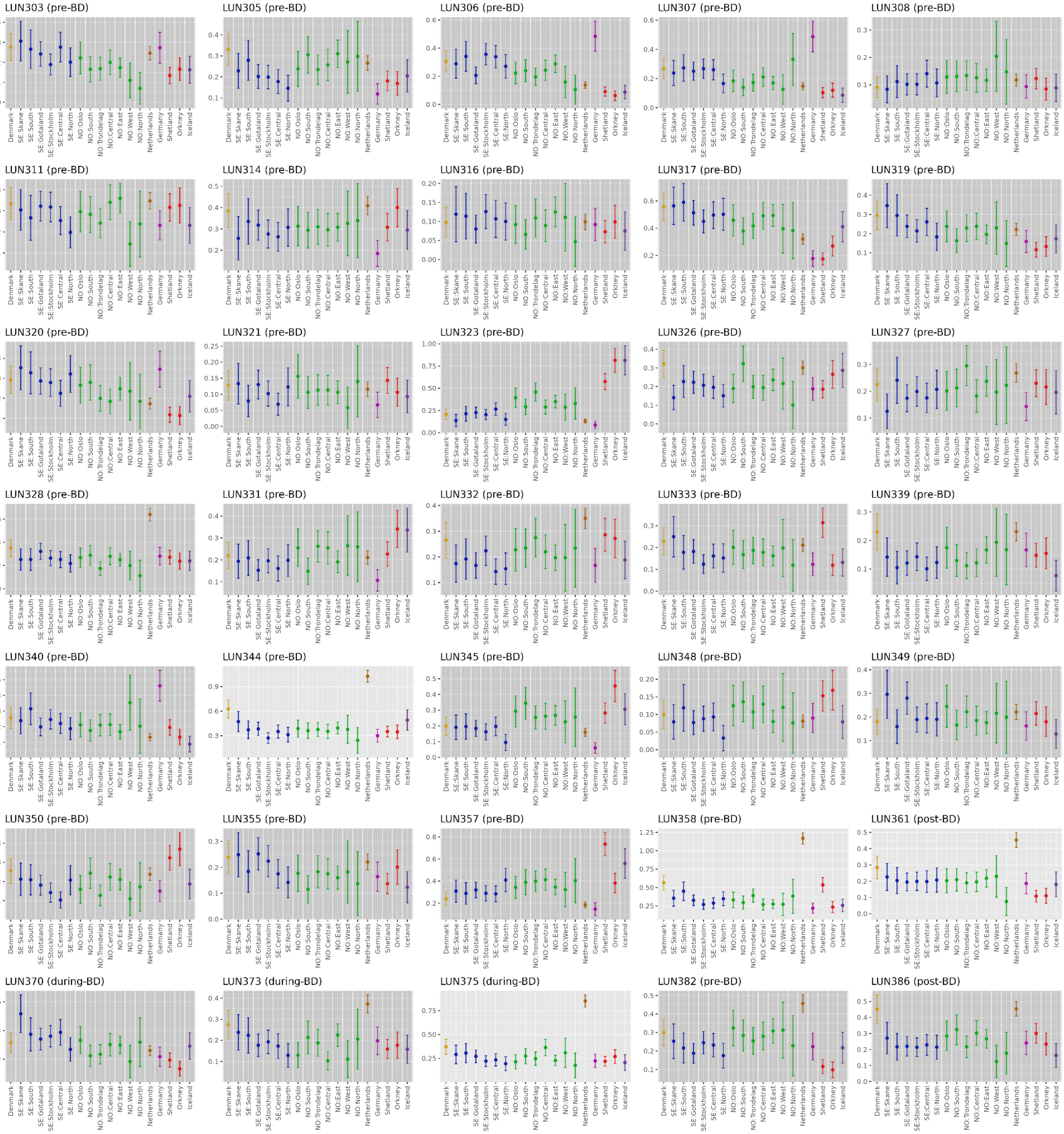
LUN303 (pre-BD)
LUN305 (pre-BD)
LUN306 (pre-BD)
LUN307 (pre-BD)
LUN308 (pre-BD)
LUN311 (pre-BD)
LUN314 (pre-BD)
LUN316 (pre-BD)
LUN317 (pre-BD)
LUN319 (pre-BD)
LUN320 (pre-BD)
LUN321 (pre-BD)
LUN323 (pre-BD)
LUN326 (pre-BD)
LUN327 (pre-BD)
LUN328 (pre-BD)
LUN331 (pre-BD)
LUN332 (pre-BD)
LUN333 (pre-BD)
LUN339 (pre-BD)
LUN340 (pre-BD)
LUN344 (pre-BD)
LUN345 (pre-BD)
LUN348 (pre-BD)
LUN349 (pre-BD)
LUN350 (pre-BD)
LUN355 (pre-BD)
LUN357 (pre-BD)
LUN358 (pre-BD)
LUN361 (post-BD)
LUN370 (during-BD)
LUN373 (during-BD)
LUN375 (during-BD)
LUN382 (pre-BD)
LUN386 (post-BD)
Denmark
SE:Skane
SE:South
SE:Gotaland
SE:Stockholm
SE:Central
SE:North
NO:Oslo
NO:South
NO:Trondelag
NO:Central
NO:East
NO:West
NO:North
Netherlands
Germany
Shetland
Orkney
Iceland

(continued)

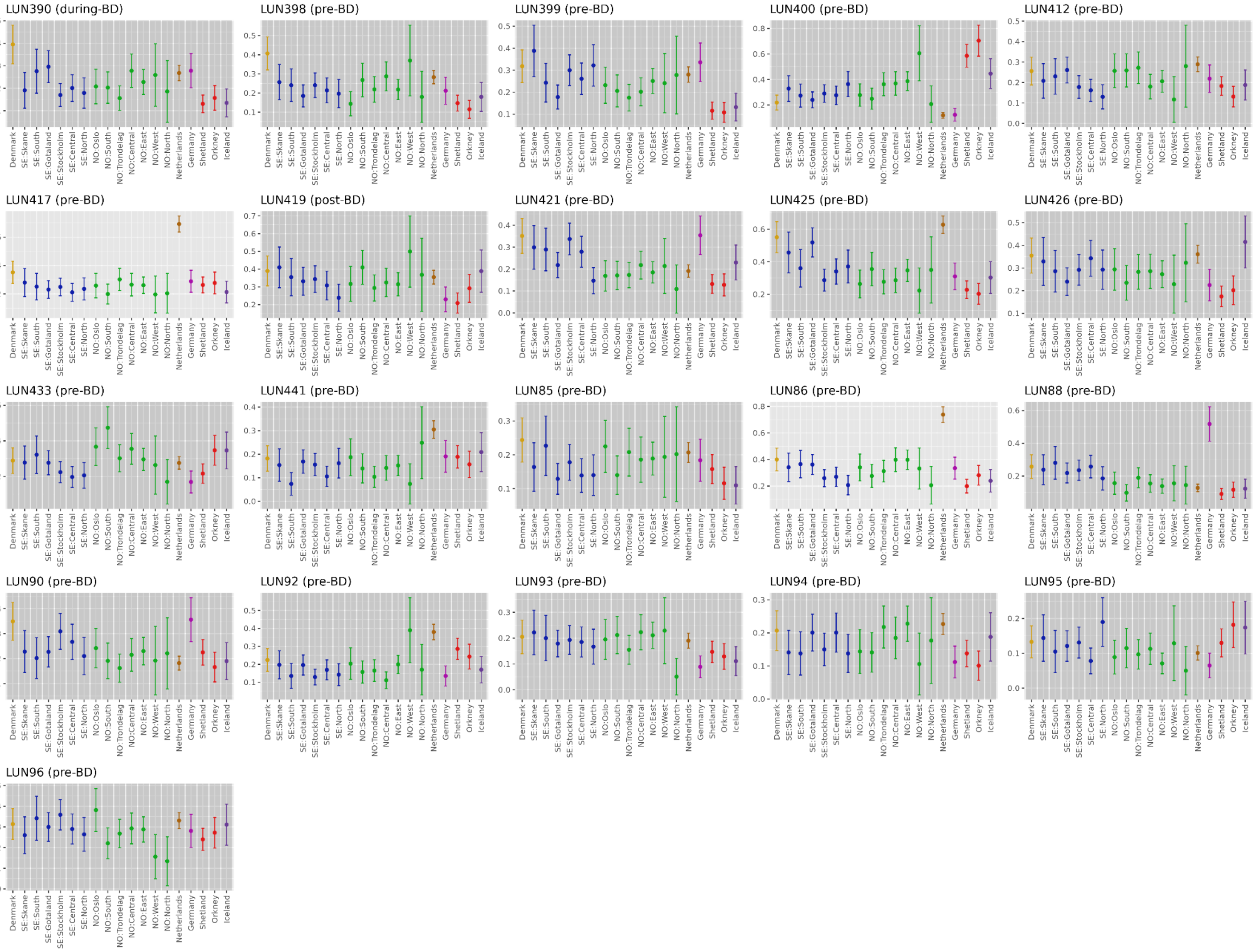

LUN390 (during-BD)
LUN398 (pre-BD)
LUN399 (pre-BD)
LUN400 (pre-BD)
LUN412 (pre-BD)
LUN417 (pre-BD)
LUN419 (post-BD)
LUN421 (pre-BD)
LUN425 (pre-BD)
LUN426 (pre-BD)
LUN433 (pre-BD)
LUN441 (pre-BD)
LUN85 (pre-BD)
LUN86 (pre-BD)
LUN88 (pre-BD)
LUN90 (pre-BD)
LUN92 (pre-BD)
LUN93 (pre-BD)
LUN94 (pre-BD)
LUN95 (pre-BD)
LUN96 (pre-BD)
Denmark
SE:Skane
SE:South
SE:Gotaland
SE:Stockholm
SE:Central
SE:North
NO:Oslo
NO:South
NO:Trondelag
NO:Central
NO:East
NO:West
NO:North
Netherlands
Germany
Shetland
Orkney
Iceland

**B)** IBD profiles for individuals classified as non-local in Lund

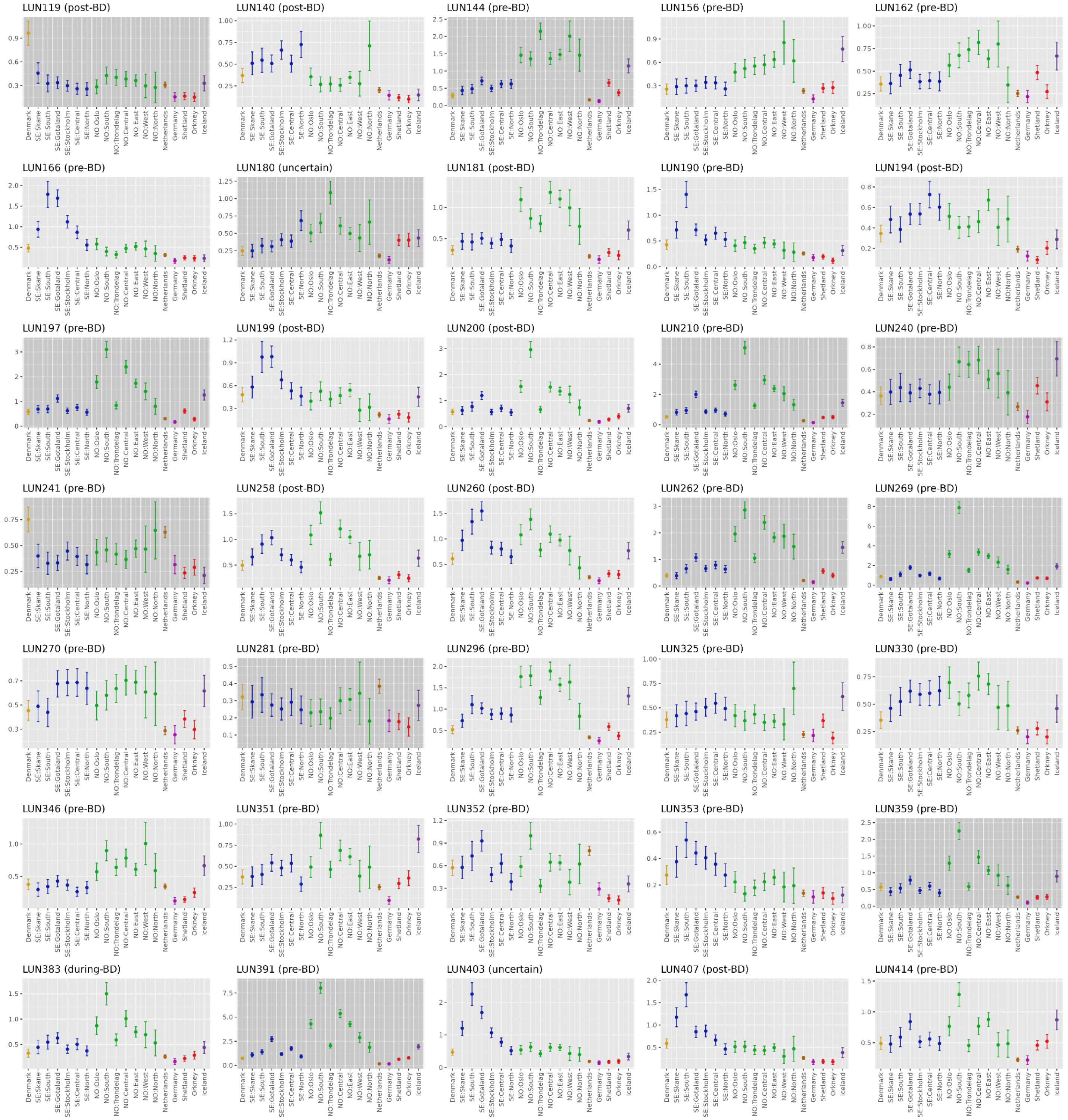

(continued)

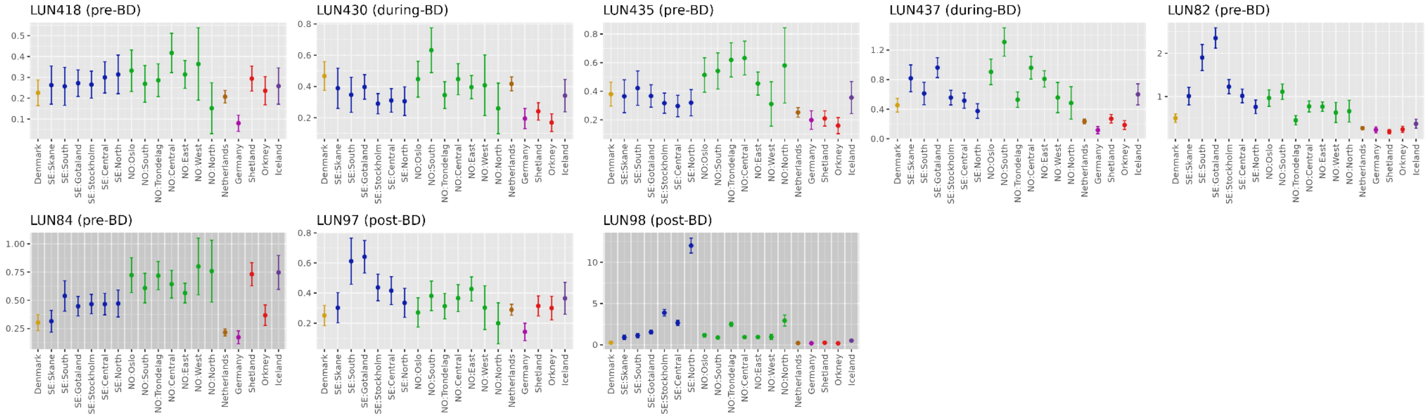
LUN418 (pre-BD)
LUN430 (during-BD)
LUN435 (pre-BD)
LUN437 (during-BD)
LUN82 (pre-BD)
LUN84 (pre-BD)
LUN97 (post-BD)
LUN98 (post-BD)
Denmark
SE:Skane
SE:South
SE:Gotaland
SE:Stockholm
SE:Central
SE:North
NO:Oslo
NO:South
NO:Trondelag
NO:Central
NO:East
NO:West
NO:North
Netherlands
Germany
Shetland
Orkney
Iceland

**C)** IBD profiles for individuals classified as local in Lund

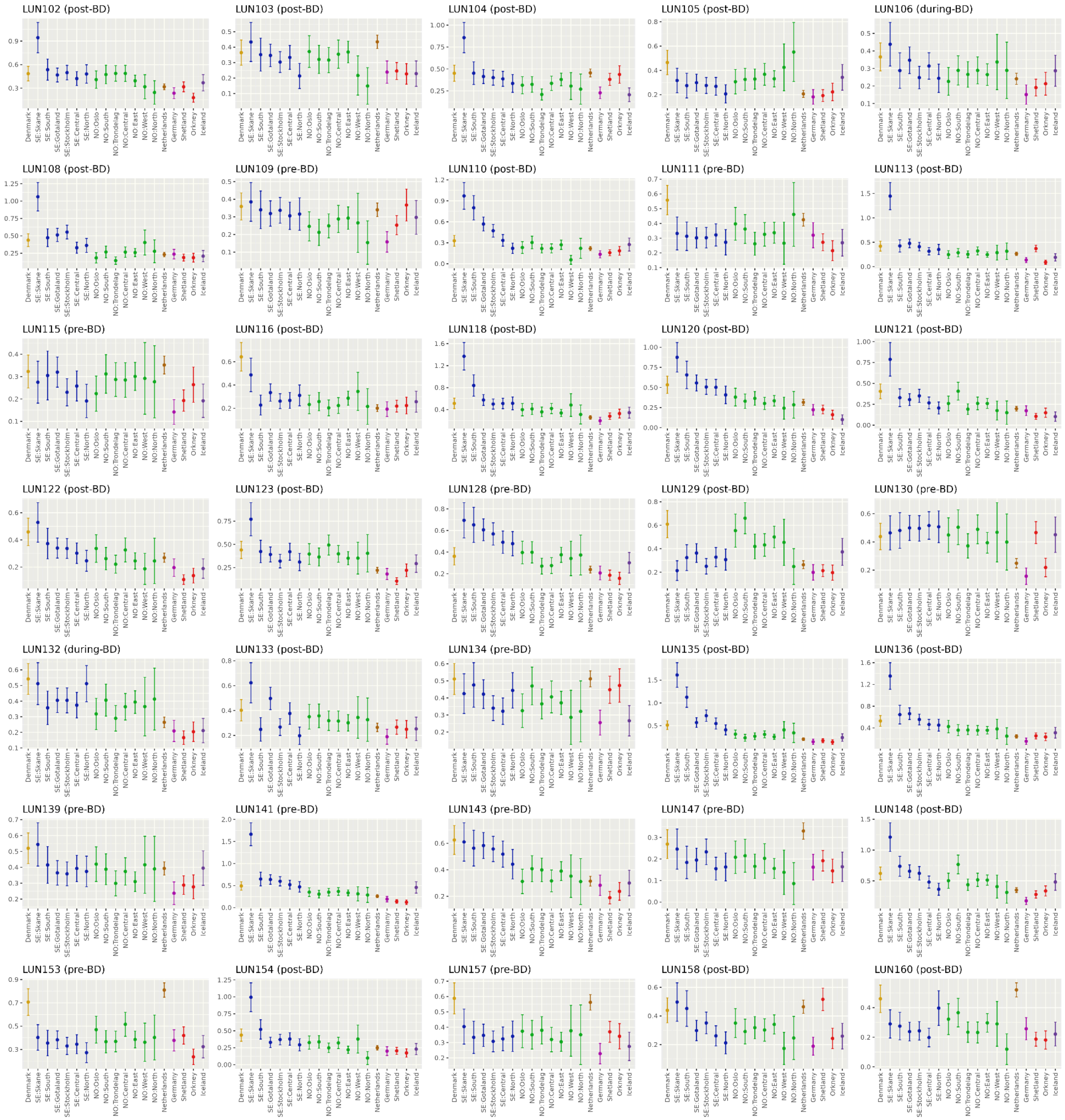

(continued)

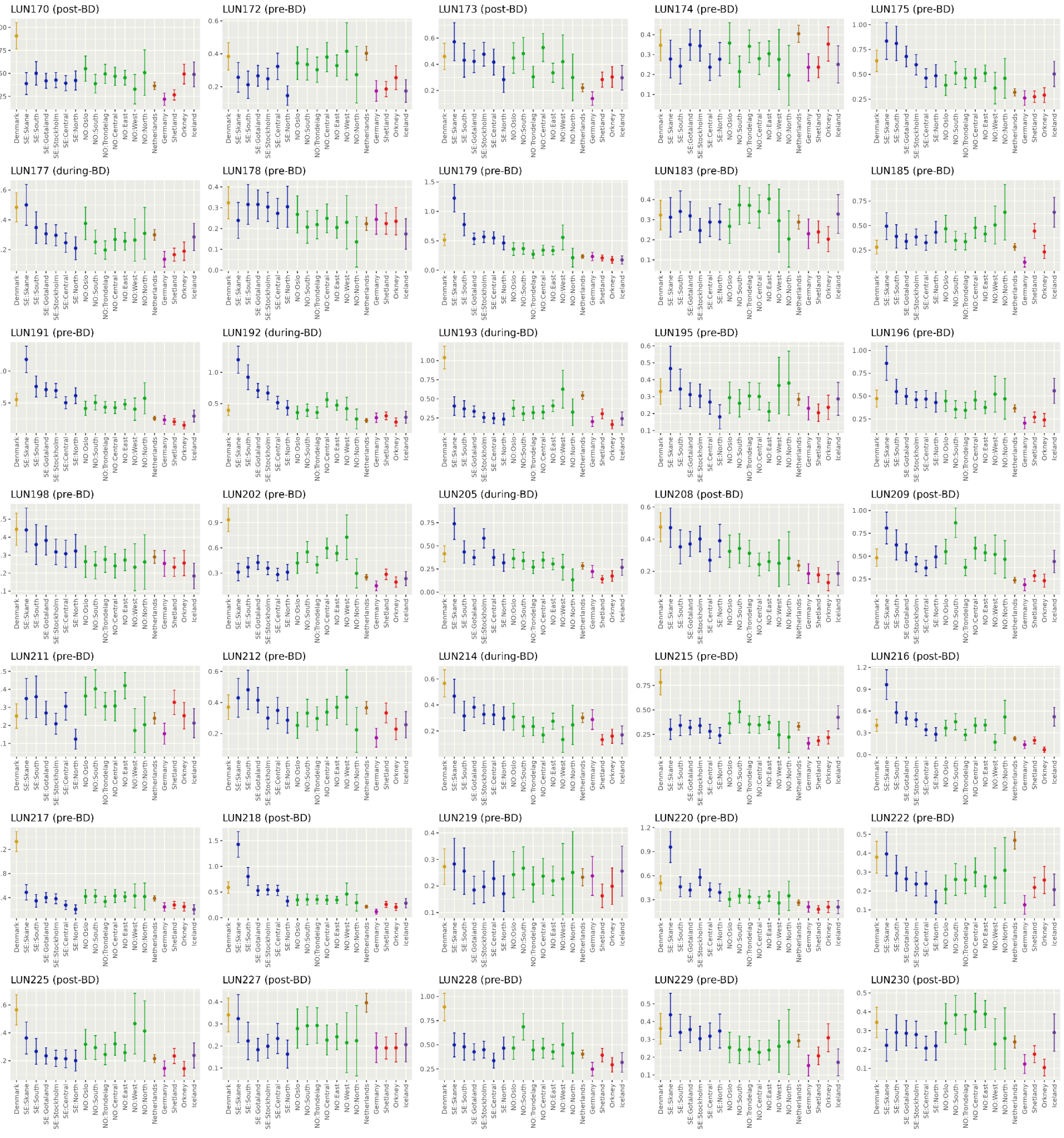
LUN170 (post-BD)
LUN172 (pre-BD)
LUN173 (post-BD)
LUN174 (pre-BD)
LUN175 (pre-BD)
LUN177 (during-BD)
LUN178 (pre-BD)
LUN179 (pre-BD)
LUN183 (pre-BD)
LUN185 (pre-BD)
LUN191 (pre-BD)
LUN192 (during-BD)
LUN193 (during-BD)
LUN195 (pre-BD)
LUN196 (pre-BD)
LUN198 (pre-BD)
LUN202 (pre-BD)
LUN205 (during-BD)
LUN208 (post-BD)
LUN209 (post-BD)
LUN211 (pre-BD)
LUN212 (pre-BD)
LUN214 (during-BD)
LUN215 (pre-BD)
LUN216 (post-BD)
LUN217 (pre-BD)
LUN218 (post-BD)
LUN219 (pre-BD)
LUN220 (pre-BD)
LUN222 (pre-BD)
LUN225 (post-BD)
LUN227 (post-BD)
LUN228 (pre-BD)
LUN229 (pre-BD)
LUN230 (post-BD)
Denmark
SE:Skane
SE:South
SE:Gotaland
SE:Stockholm
SE:Central
SE:North
NO:Oslo
NO:South
NO:Trondelag
NO:Central
NO:East
NO:West
NO:North
Netherlands
Germany
Shetland
Orkney
Iceland

(continued)

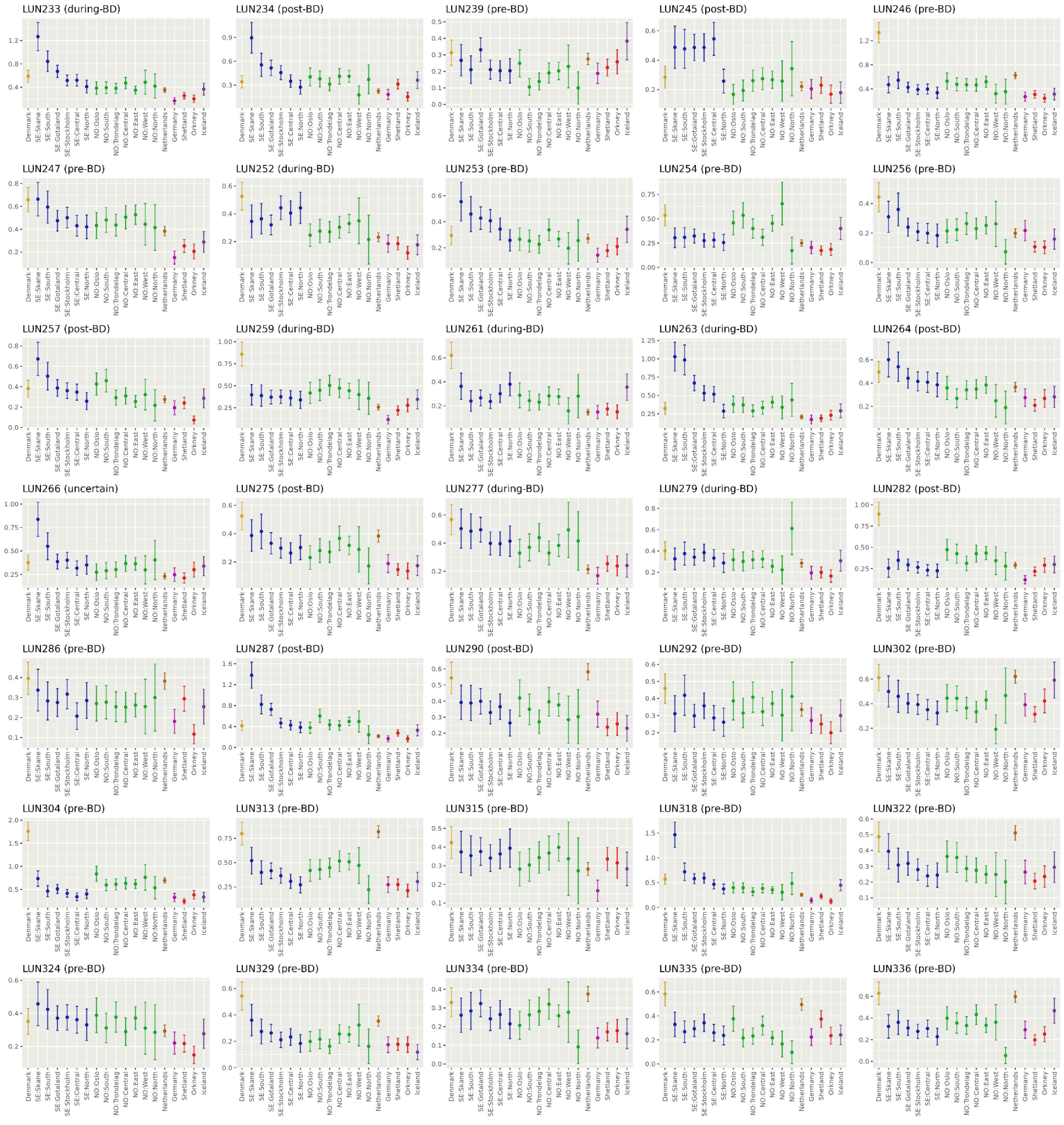
LUN233 (during-BD)
LUN234 (post-BD)
LUN239 (pre-BD)
LUN245 (post-BD)
LUN246 (pre-BD)
LUN247 (pre-BD)
LUN252 (during-BD)
LUN253 (pre-BD)
LUN254 (pre-BD)
LUN256 (pre-BD)
LUN257 (post-BD)
LUN259 (during-BD)
LUN261 (during-BD)
LUN263 (during-BD)
LUN264 (post-BD)
LUN266 (uncertain)
LUN275 (post-BD)
LUN277 (during-BD)
LUN279 (during-BD)
LUN282 (post-BD)
LUN286 (pre-BD)
LUN287 (post-BD)
LUN290 (post-BD)
LUN292 (pre-BD)
LUN302 (pre-BD)
LUN304 (pre-BD)
LUN313 (pre-BD)
LUN315 (pre-BD)
LUN318 (pre-BD)
LUN322 (pre-BD)
LUN324 (pre-BD)
LUN329 (pre-BD)
LUN334 (pre-BD)
LUN335 (pre-BD)
LUN336 (pre-BD)
Denmark
SE:Skane
SE:South
SE:Gotaland
SE:Stockholm
SE:Central
SE:North
NO:Oslo
NO:South
NO:Trondelag
NO:Central
NO:East
NO:West
NO:North
Netherlands
Germany
Shetland
Orkney
Iceland

(continued)

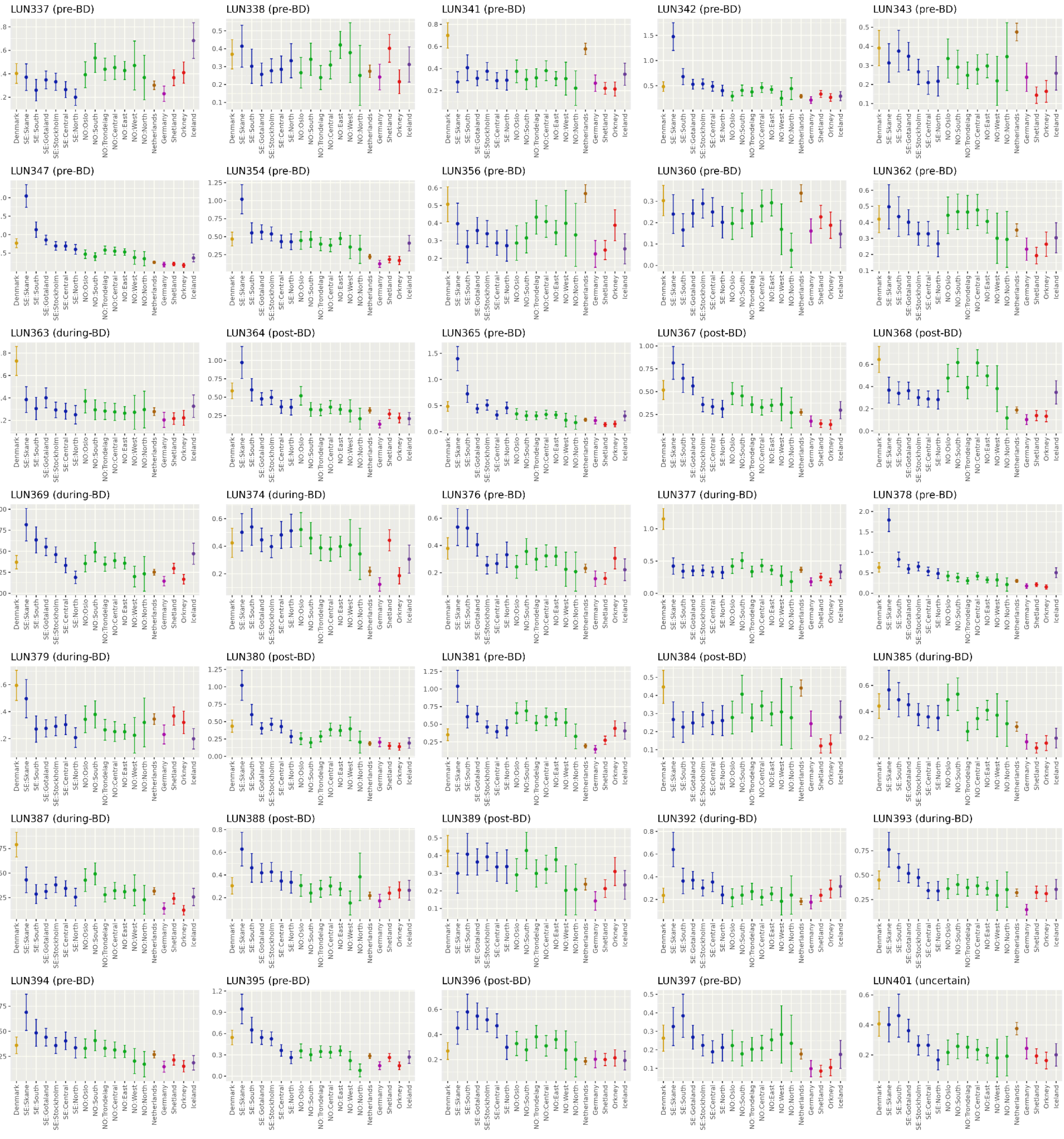
LUN337 (pre-BD)
LUN338 (pre-BD)
LUN341 (pre-BD)
LUN342 (pre-BD)
LUN343 (pre-BD)
LUN347 (pre-BD)
LUN354 (pre-BD)
LUN356 (pre-BD)
LUN360 (pre-BD)
LUN362 (pre-BD)
LUN363 (during-BD)
LUN364 (post-BD)
LUN365 (pre-BD)
LUN367 (post-BD)
LUN368 (post-BD)
LUN369 (during-BD)
LUN374 (during-BD)
LUN376 (pre-BD)
LUN377 (during-BD)
LUN378 (pre-BD)
LUN379 (during-BD)
LUN380 (post-BD)
LUN381 (pre-BD)
LUN384 (post-BD)
LUN385 (during-BD)
LUN387 (during-BD)
LUN388 (post-BD)
LUN389 (post-BD)
LUN392 (during-BD)
LUN393 (during-BD)
LUN394 (pre-BD)
LUN395 (pre-BD)
LUN396 (post-BD)
LUN397 (pre-BD)
LUN401 (uncertain)
Denmark
SE:Skane
SE:South
SE:Gotaland
SE:Stockholm
SE:Central
SE:North
NO:Oslo
NO:South
NO:Trondelag
NO:Central
NO:East
NO:West
NO:North
Netherlands
Germany
Shetland
Orkney
Iceland

(continued)

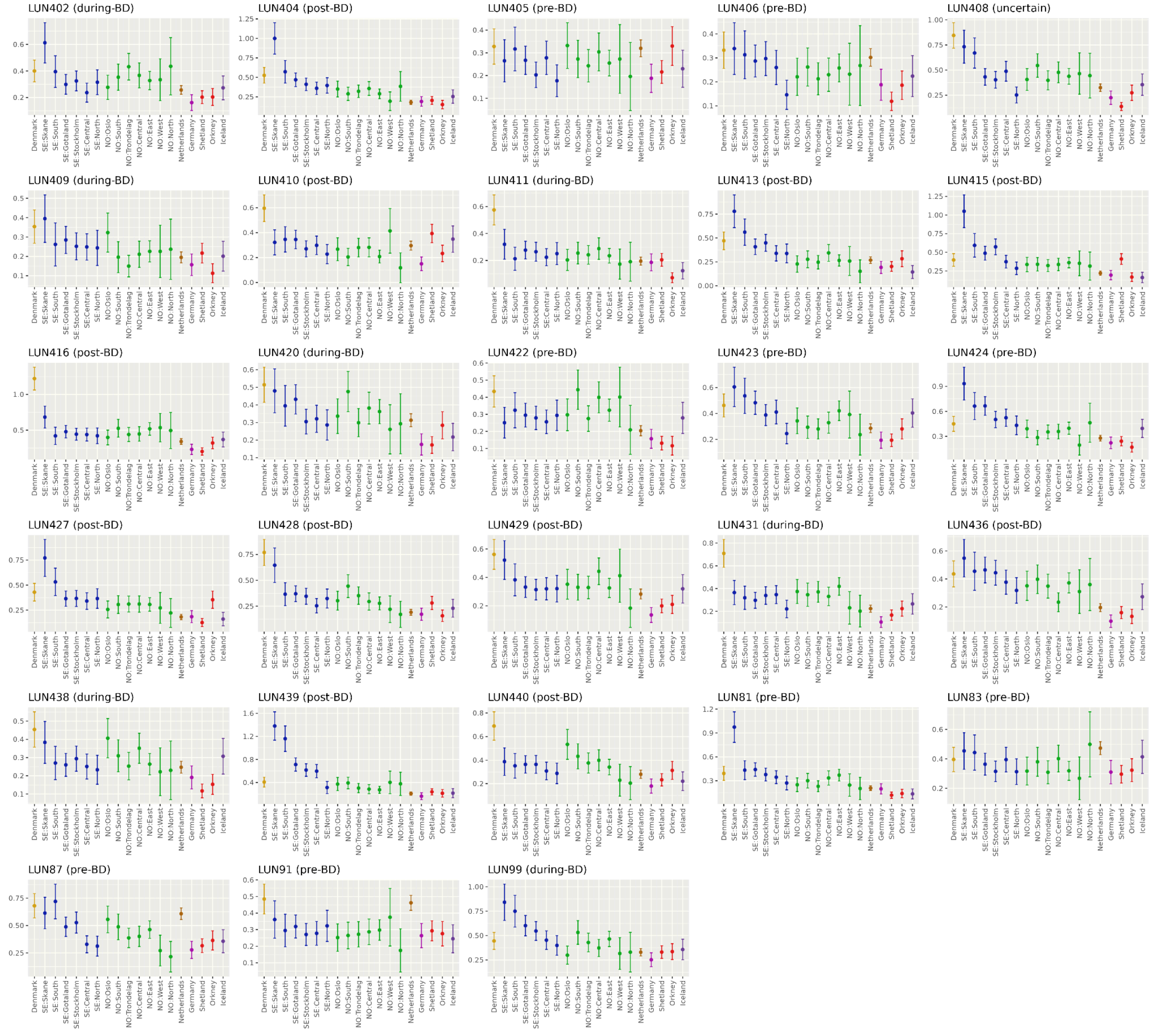


**Supplementary Figure 18.** IBD sharing profiles for ancient individuals from Lund categorized as **(A)** outliers (non-Scandinavian), **(B)** non-local Scandinavian, and **(C)** local Scandinavian based on ancestry classification (see Methods). Each panel shows average IBD between the corresponding ancient individual (indicated by the panel title) and present-day reference populations from the OmniExpress European panel (Supplementary Table 5). Error bars indicate 95% confidence intervals. Dark and light gray backgrounds indicate PCA- and IBD- based classifications, respectively, for outliers in **A** and non-local Scandinavian individuals in **B**. See Supplementary Figure 31 for a simplified summary.

**A)** IBD profiles for individuals classified as non-Scandinavian in Trondheim

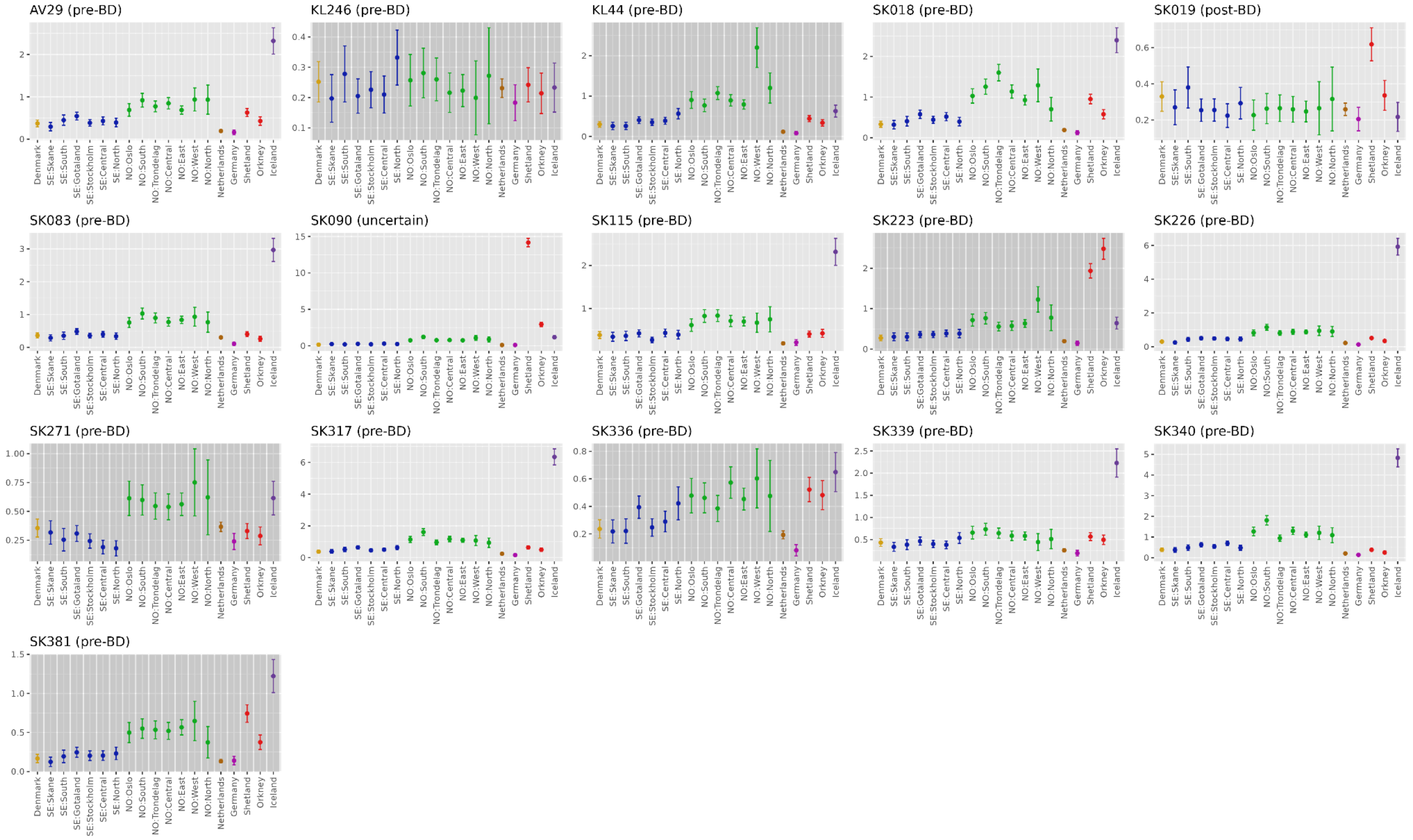

**B)** IBD profiles for individuals classified as non-local in Trondheim

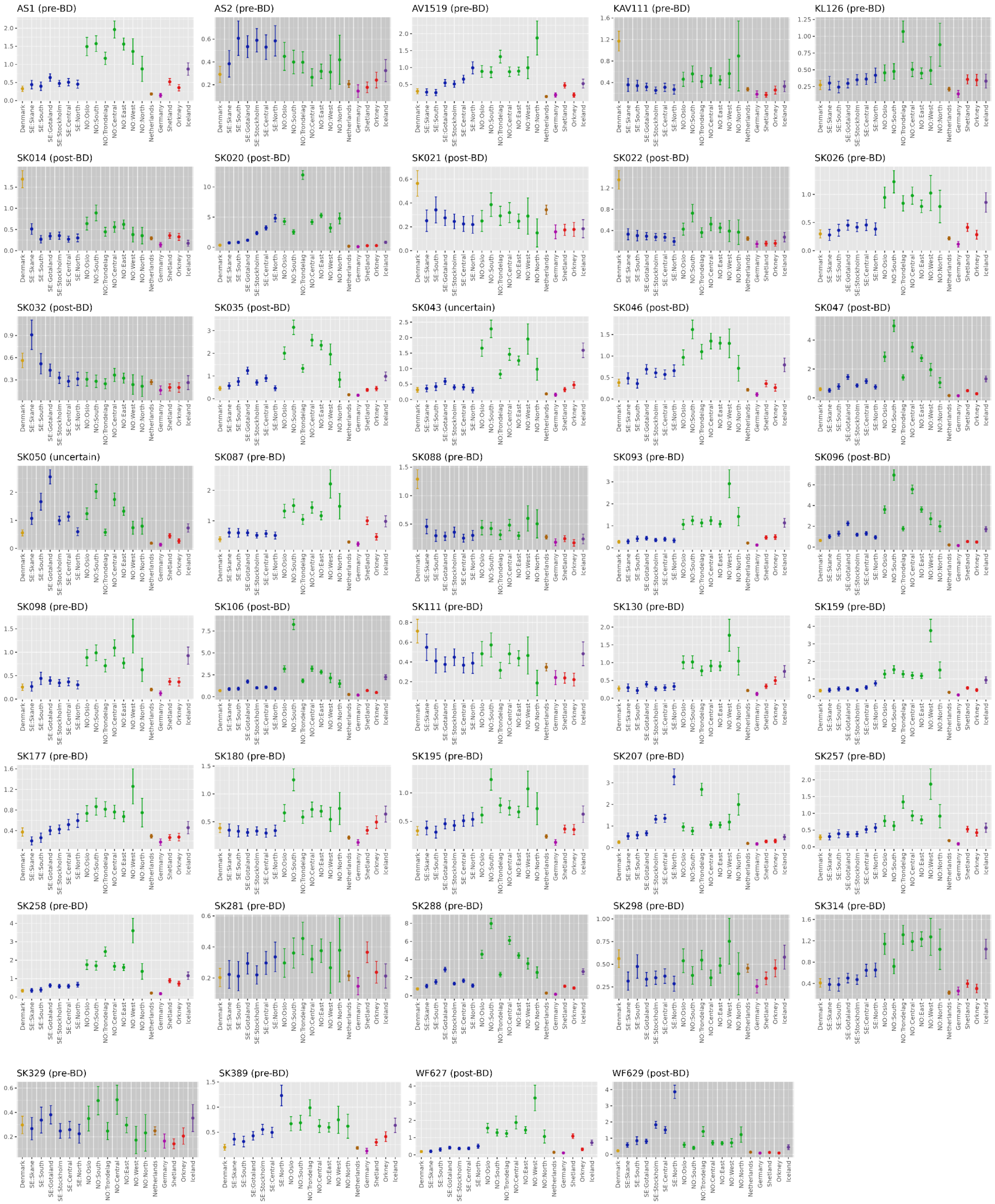

**C)** IBD profiles for individuals classified as local in Trondheim

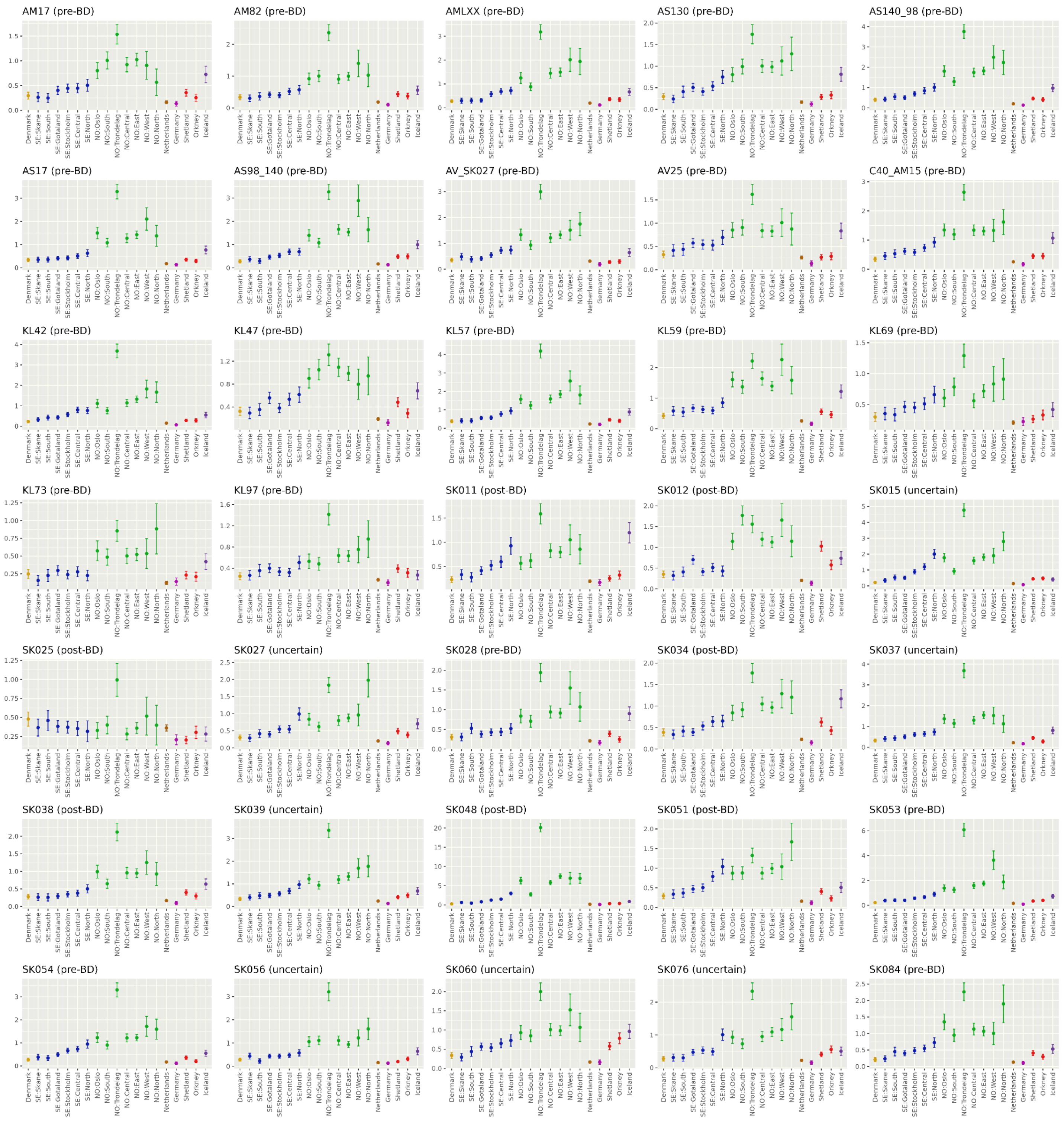

(continued)

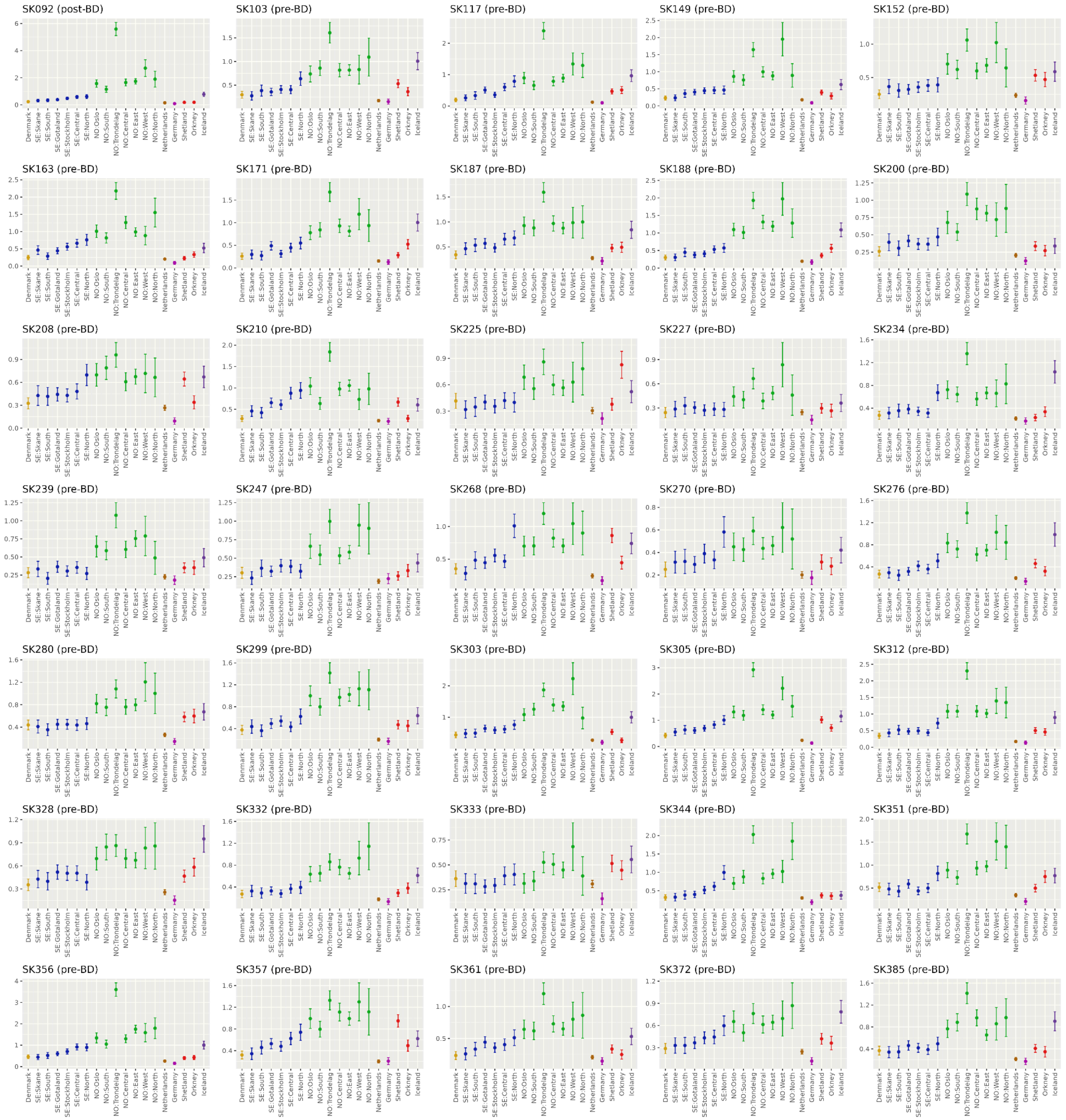

SK092 (post-BD)
SK103 (pre-BD)
SK117 (pre-BD)
SK149 (pre-BD)
SK152 (pre-BD)
SK163 (pre-BD)
SK171 (pre-BD)
SK187 (pre-BD)
SK188 (pre-BD)
SK200 (pre-BD)
SK208 (pre-BD)
SK210 (pre-BD)
SK225 (pre-BD)
SK227 (pre-BD)
SK234 (pre-BD)
SK239 (pre-BD)
SK247 (pre-BD)
SK268 (pre-BD)
SK270 (pre-BD)
SK276 (pre-BD)
SK280 (pre-BD)
SK299 (pre-BD)
SK303 (pre-BD)
SK305 (pre-BD)
SK312 (pre-BD)
SK328 (pre-BD)
SK332 (pre-BD)
SK333 (pre-BD)
SK344 (pre-BD)
SK351 (pre-BD)
SK356 (pre-BD)
SK357 (pre-BD)
SK361 (pre-BD)
SK372 (pre-BD)
SK385 (pre-BD)
Denmark
SE:Skane
SE:South
SE:Gotaland
SE:Stockholm
SE:Central
SE:North
NO:Oslo
NO:South
NO:Trondelag
NO:Central
NO:East
NO:West
NO:North
Netherlands
Germany
Shetland
Orkney
Iceland

(continued)

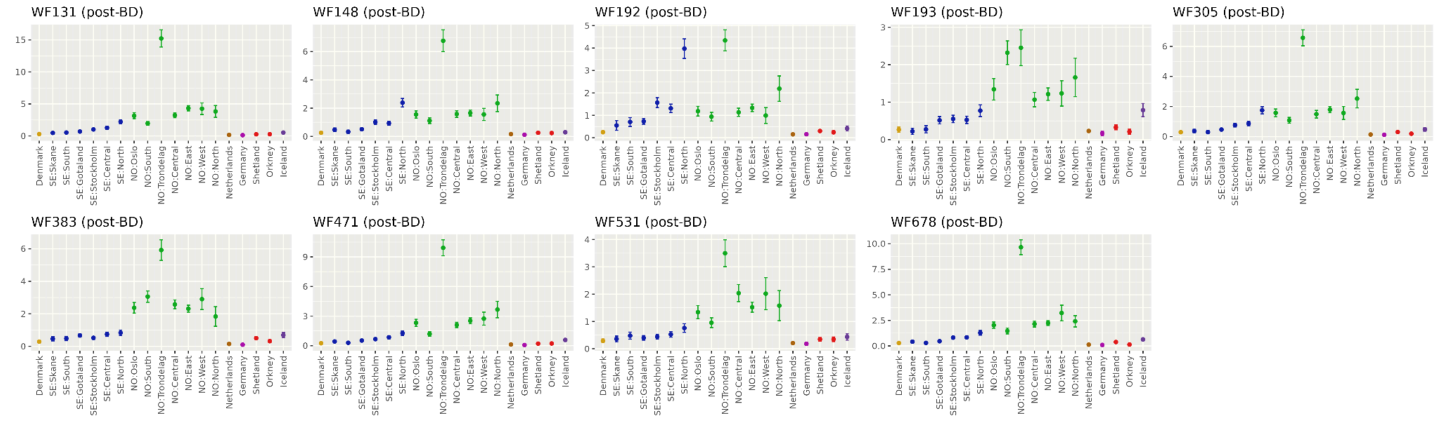


**Supplementary Figure 19.** IBD sharing profiles for ancient individuals from Trondheim categorized as **(A)** outliers (non-Scandinavian), **(B)** non-local Scandinavian, and **(C)** local Scandinavian based on ancestry classification (see Methods). Each panel shows average IBD between the corresponding ancient individual (indicated by the panel title) and present-day reference populations from the OmniExpress European panel (Supplementary Table 5). Error bars indicate 95% confidence intervals. Dark and light gray backgrounds indicate PCA- and IBD- based classifications, respectively, for outliers in **A** and non-local Scandinavian individuals in **B**. See Supplementary Figure 31 for a simplified summary.

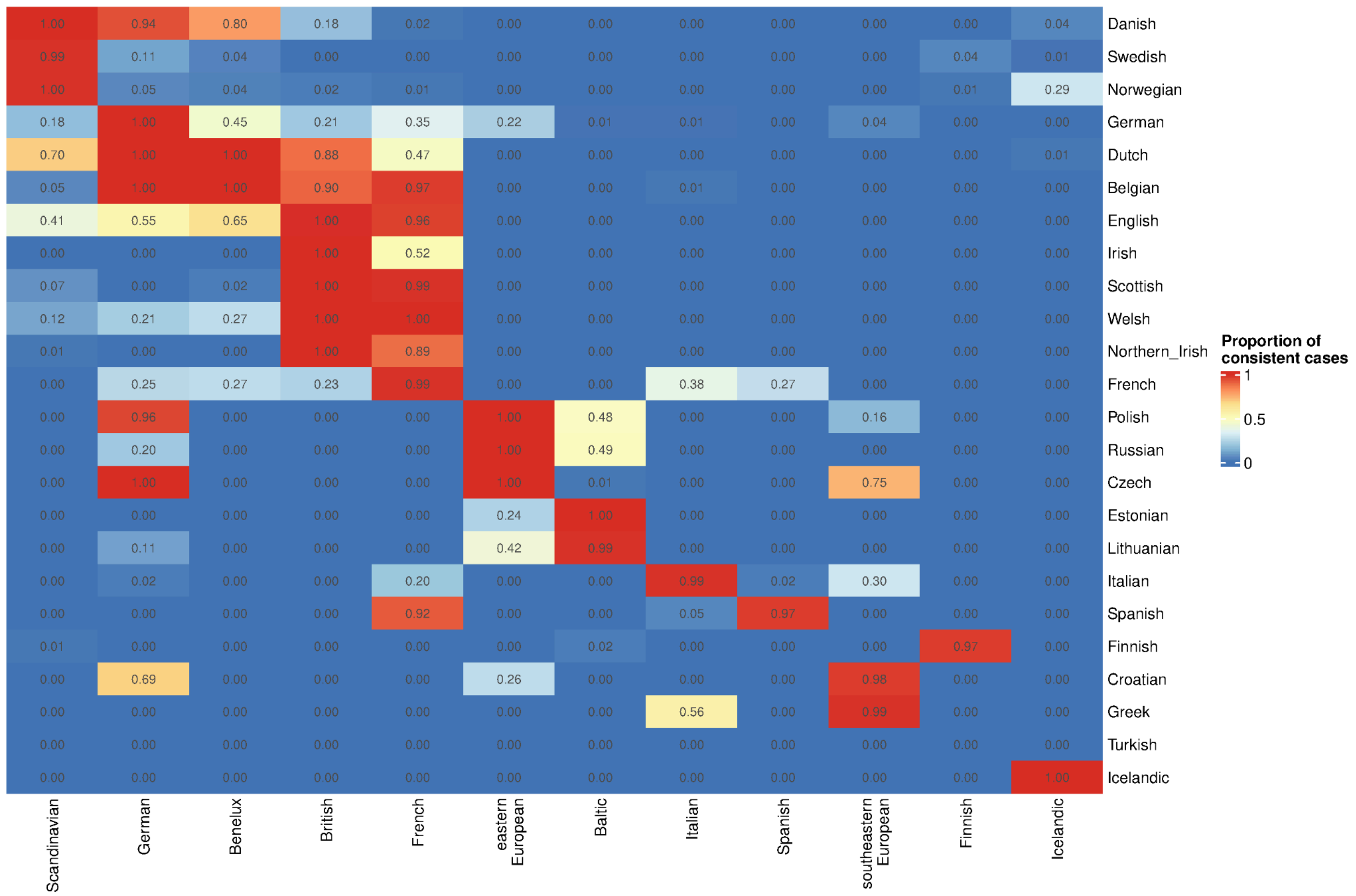


**Supplementary Figure 20.** Summary of leave-one-out test results based on Mahalanobis distances ($D_M$) calculated using the first seven PCs after projection onto the PCA constructed from 67 West Eurasian populations. Rows represent individual source populations, whereas columns represent broad reference groups. These broad groups were used to mimic the PCA-based outlier detection procedures applied to ancient individuals, including tests for non-Scandinavian outliers in the Lund and Trondheim datasets and non-Baltic outliers in the Vilnius dataset. Each cell shows the proportion of individuals from the row population that are consistent with at least one population within the corresponding column group according to a chi-square test (df=7, $P$>0.001). Higher values indicate greater genetic similarity between the row population and the column group. Column groups are defined as follows: Scandinavian, Danish/Swedish/Norwegian; German, German; Benelux, Dutch/Belgian; British, English/Irish/Scottish/Welsh/Northern Irish; French, French; eastern European, Polish/Russian/Czech; Baltic, Lithuanian/Estonian; Italian, Italian; Spanish, Spanish; southeastern European, Croatian/Greek; Finnish, Finnish; and Icelandic, Icelandic.

#### A) IBD profiles for non-Baltic samples in Vilnius

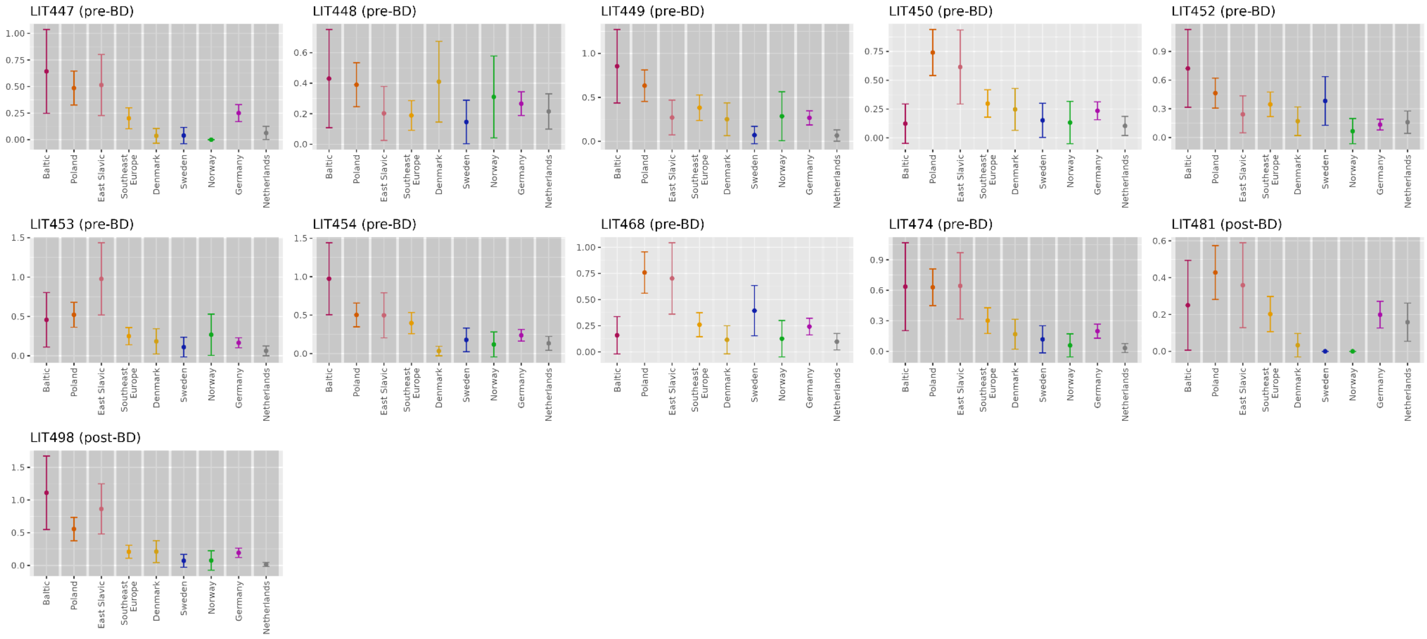


#### B) IBD profiles for Baltic samples in Vilnius

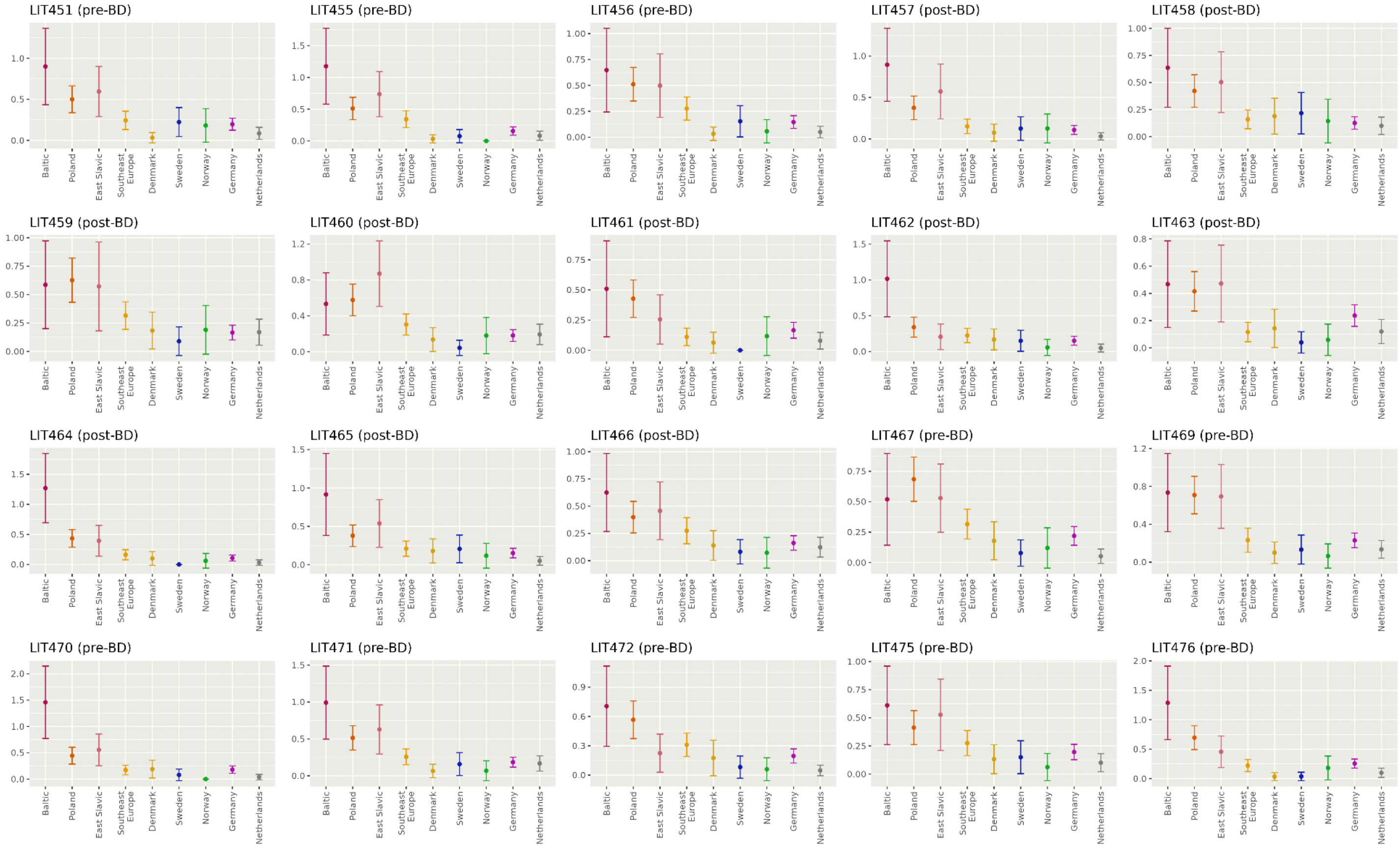

(continued)

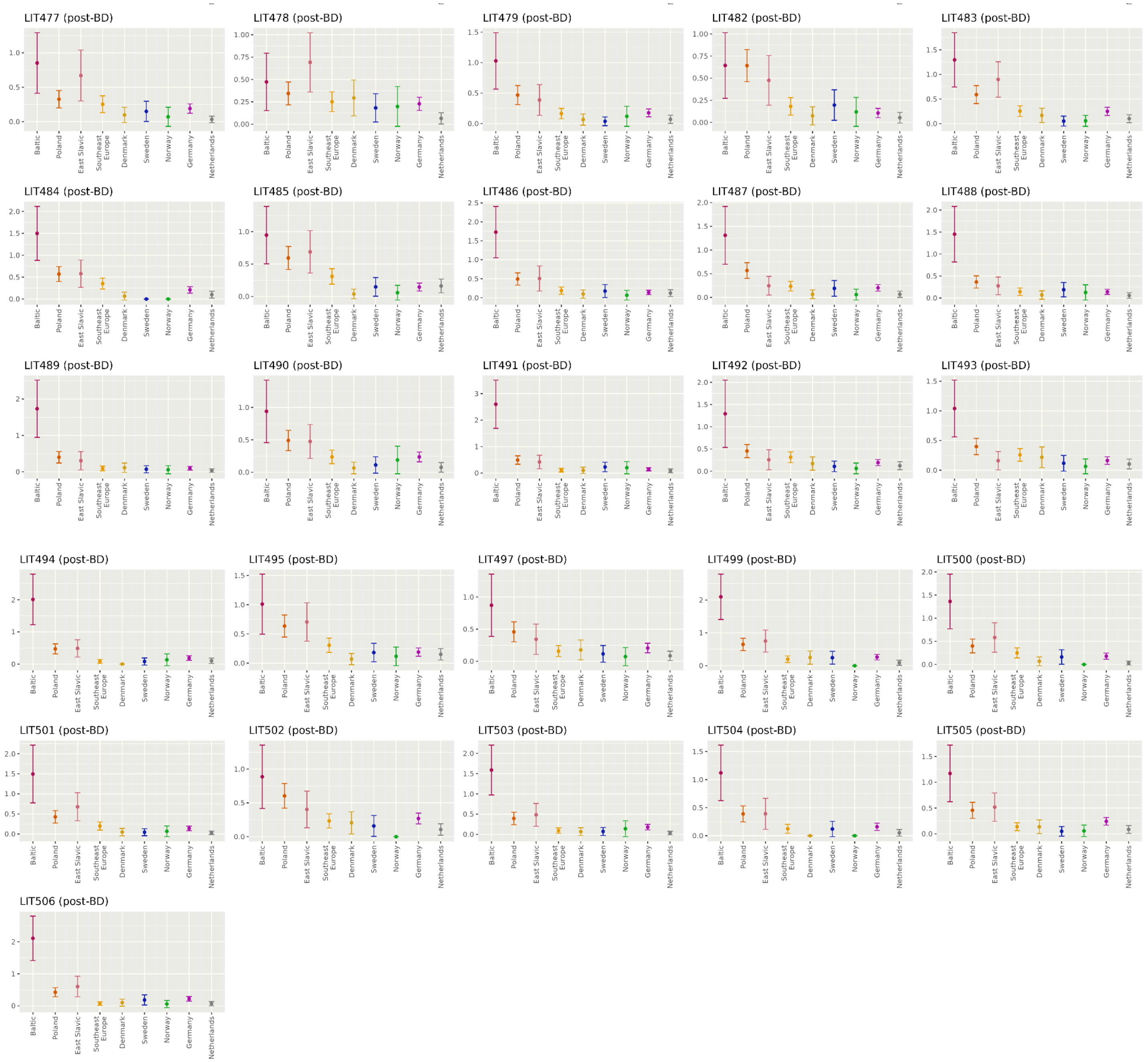


**Supplementary Figure 21.** IBD sharing profiles for ancient individuals from Vilnius categorized as **(A)** outliers (non-Baltic) and **(B)** Baltic based on ancestry classification (see Methods). Each panel shows average IBD between the corresponding ancient individual (indicated by the panel title) and present-day reference populations from the UK Biobank panel (Supplementary Table 5). Error bars indicate 95% confidence intervals. In **A**, dark gray background denotes individuals classified as outliers by PCA, and light gray background denotes individuals classified as outliers by IBD-sharing. See Supplementary Figure 31 for a simplified summary.

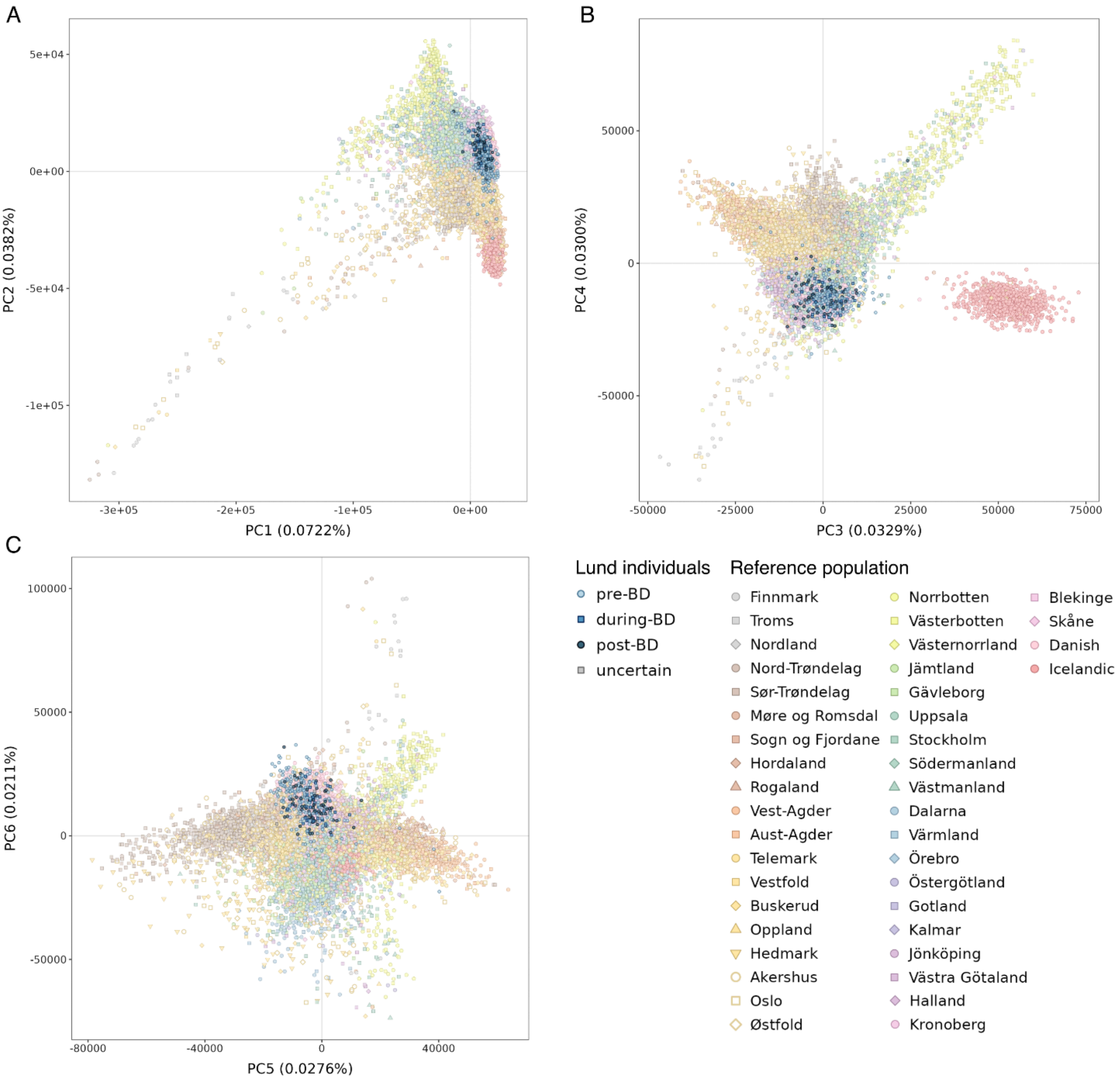
A
B
C
5e+04
0e+00
-5e+04
-1e+05
-3e+05
-2e+05
-1e+05
0e+00
PC1 (0.0722%)
PC2 (0.0382%)
50000
0
-50000
-25000
25000
75000
PC3 (0.0329%)
PC4 (0.0300%)
100000
-80000
-40000
40000
PC5 (0.0276%)
PC6 (0.0211%)
Lund individuals
pre-BD
during-BD
post-BD
uncertain
Reference population
Finnmark
Troms
Nordland
Nord-Trøndelag
Sør-Trøndelag
Møre og Romsdal
Sogn og Fjordane
Hordaland
Rogaland
Vest-Agder
Aust-Agder
Telemark
Vestfold
Buskerud
Oppland
Hedmark
Akershus
Oslo
Østfold
Norrbotten
Västerbotten
Västernorrland
Jämtland
Gävleborg
Uppsala
Stockholm
Södermanland
Västmanland
Dalarna
Värmland
Örebro
Östergötland
Gotland
Kalmar
Jönköping
Västra Götaland
Halland
Kronoberg
Blekinge
Skåne
Danish
Icelandic

(continued)

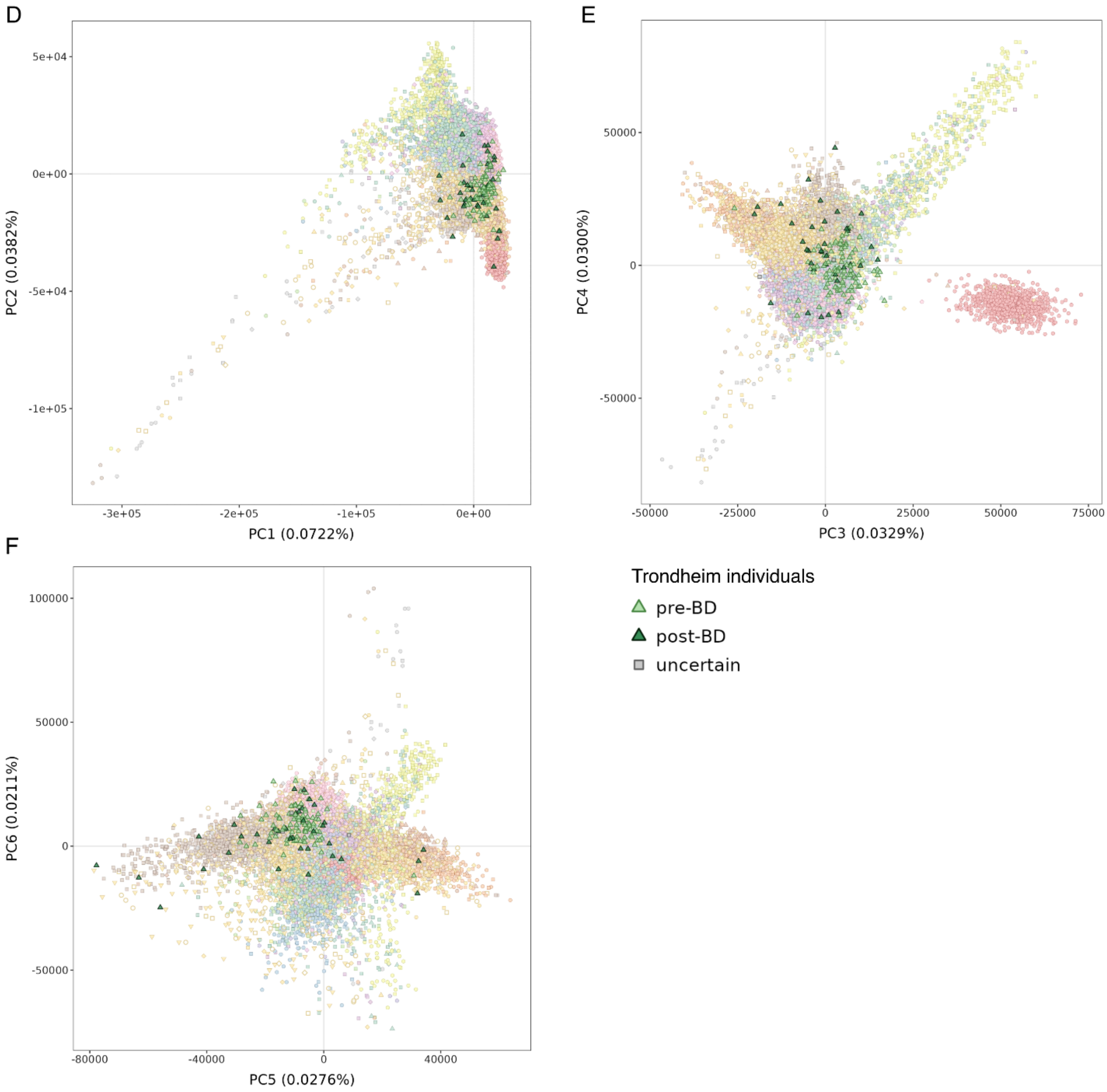
D
E
F
PC1 (0.0722%)
PC2 (0.0382%)
PC3 (0.0329%)
PC4 (0.0300%)
PC5 (0.0276%)
PC6 (0.0211%)
Trondheim individuals
pre-BD
post-BD
uncertain

(continued)

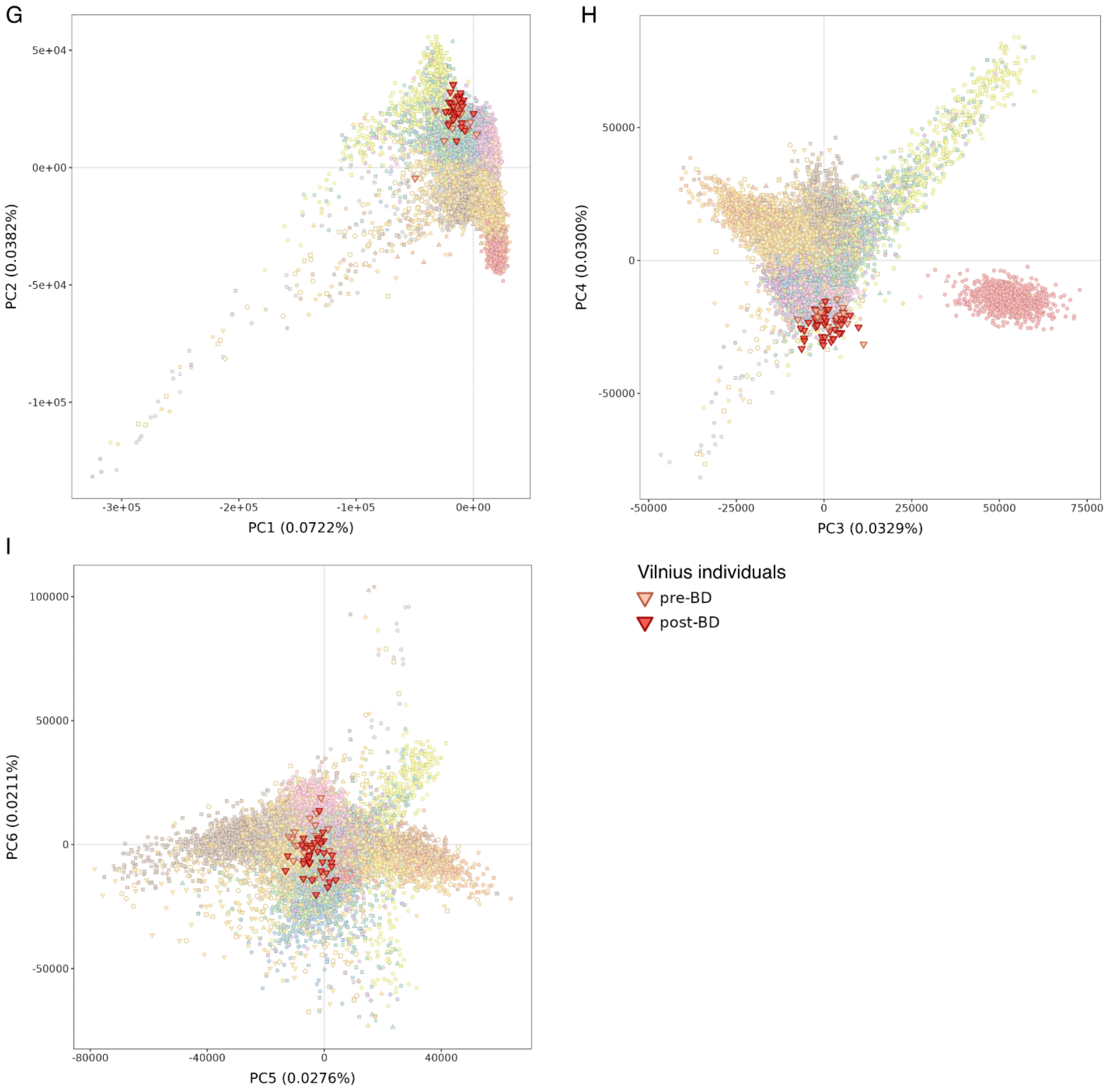


**Supplementary Figure 22.** PCA projection of ancient individuals onto present-day Scandinavian populations (subset of OmniExpress reference panel). Ancient individuals from Lund (A-D), Trondheim (E-H), and Vilnius (I-L) were projected onto a PCA constructed from 15 present-day Scandinavian groups (Supplementary Table 5). Ancient individuals are colored according to temporal group, and present-day reference populations are indicated by combinations of colour and shape.

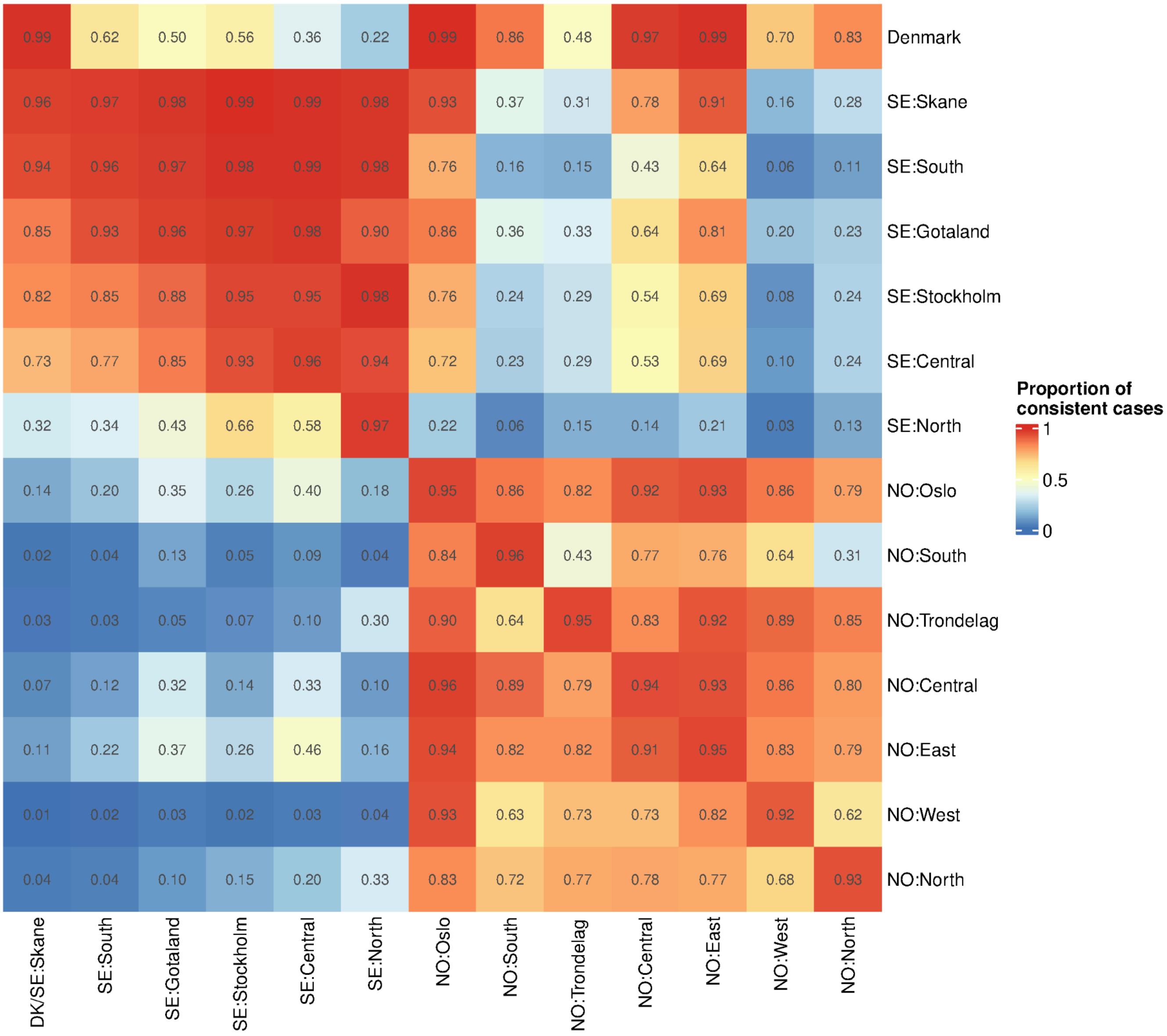


**Supplementary Figure 23.** Summary of leave-one-out test results based on Mahalanobis distances ($D_M$) calculated using the first 15 PCs after projection onto the PCA constructed from Scandinavian populations in the OmniExpress dataset. Each cell shows the proportion of individuals from the row population that are consistent with the corresponding column population according to a chi-square test (df=15, $P$>0.001). Higher values indicate greater genetic similarity between the row population and the column population. To make these tests reflect our PCA-based analysis of identifying non-local in Lund, we combined Denmark and Skåne of Sweden into a single column category (‘DK/SE:Skåne’).

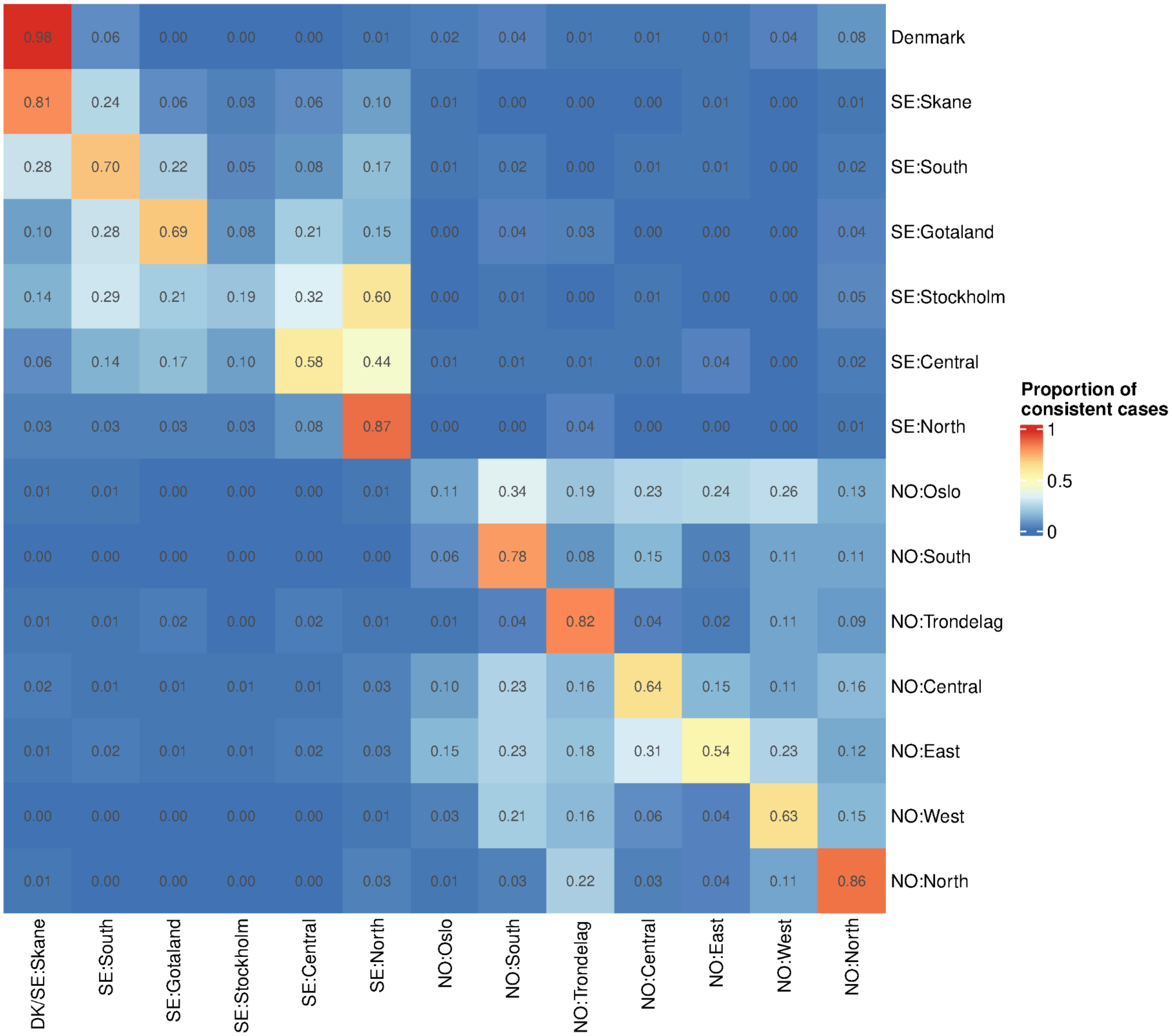


**Supplementary Figure 24.** Summary of leave-one-out results based on IBD-sharing summary metrics with Scandinavian populations in the OmniExpress dataset. Each cell represents the proportion of present-day individuals from the row population that are consistent with the population in the corresponding column. An individual was considered inconsistent with a given population (column) if it shared significantly more IBD with any other population than with that population ($P$<0.05, one-sided Welch's t-test); otherwise, it was considered consistent. Higher values indicate greater genetic similarity between the row population and the column population. To make these tests reflect our IBD-based analysis of identifying non-local individuals in Lund, we combined Denmark and Skåne of Sweden into a single column category ('DK/SE:Skåne '). For a concrete example, the cell in the first row and the first column shows that 98% of tested present-day individuals from Denmark were consistent with'DK/SE:Skåne '.

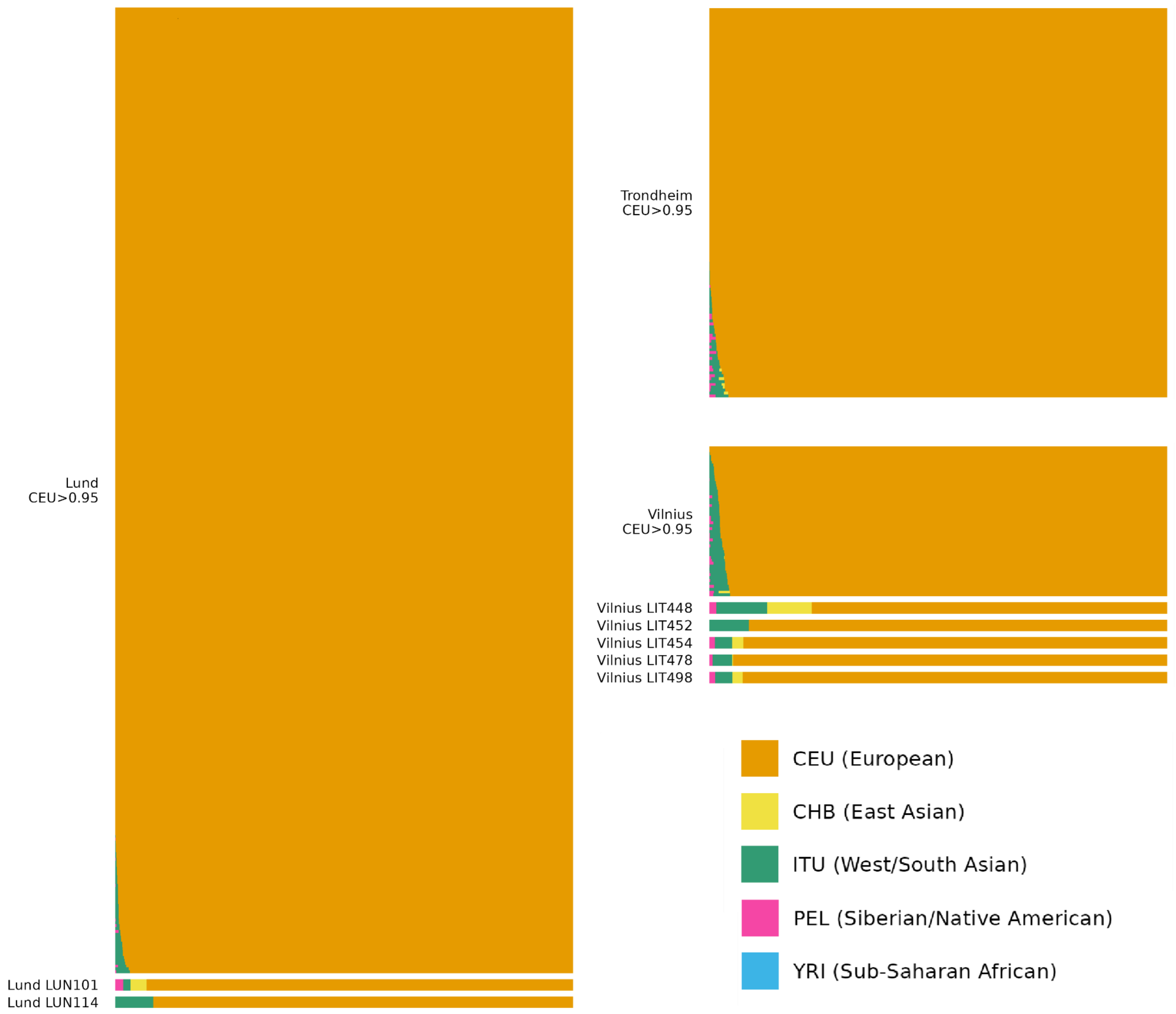


**Supplementary Figure 25.** Supervised ADMIXTURE results for ancient individuals from Lund, Trondheim, and Vilnius using K=5 continental training populations from 1000G (Supplementary Table 5). Samples assigned ≤95% CEU ancestry shown separately (none for Trondheim).

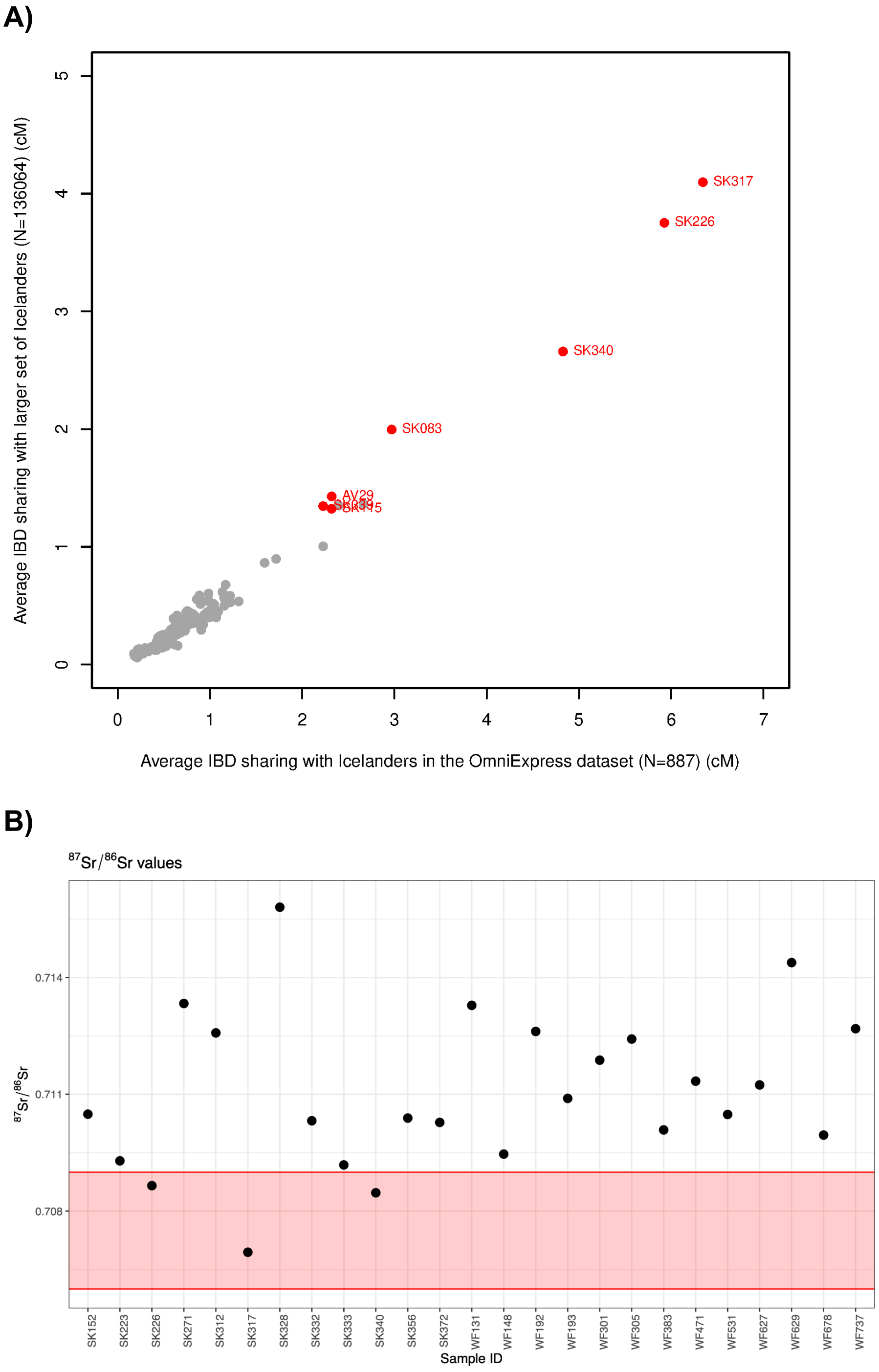


**Supplementary Figure 26.** Evidence for Icelandic origins among ancient individuals from Trondheim. **A**) Comparison of average IBD sharing , measured in cM, with two present-day Icelandic reference datasets: the OmniExpress dataset (N=887, x-axis) and a larger dataset (N=136,064, y-axis). Individuals inferred to be of Icelandic origin based on their IBD sharing profiles (see Methods) are highlighted and labelled. **B**) Strontium isotope ratio ($^{87}Sr/^{86}Sr$) for the subset of ancient individuals from Trondheim for whom strontium isotope data are available. The Icelandic local strontium isotope range is indicated by the red rectangle. The strontium isotope results for SK226, SK317 and SK340 fall within this range.

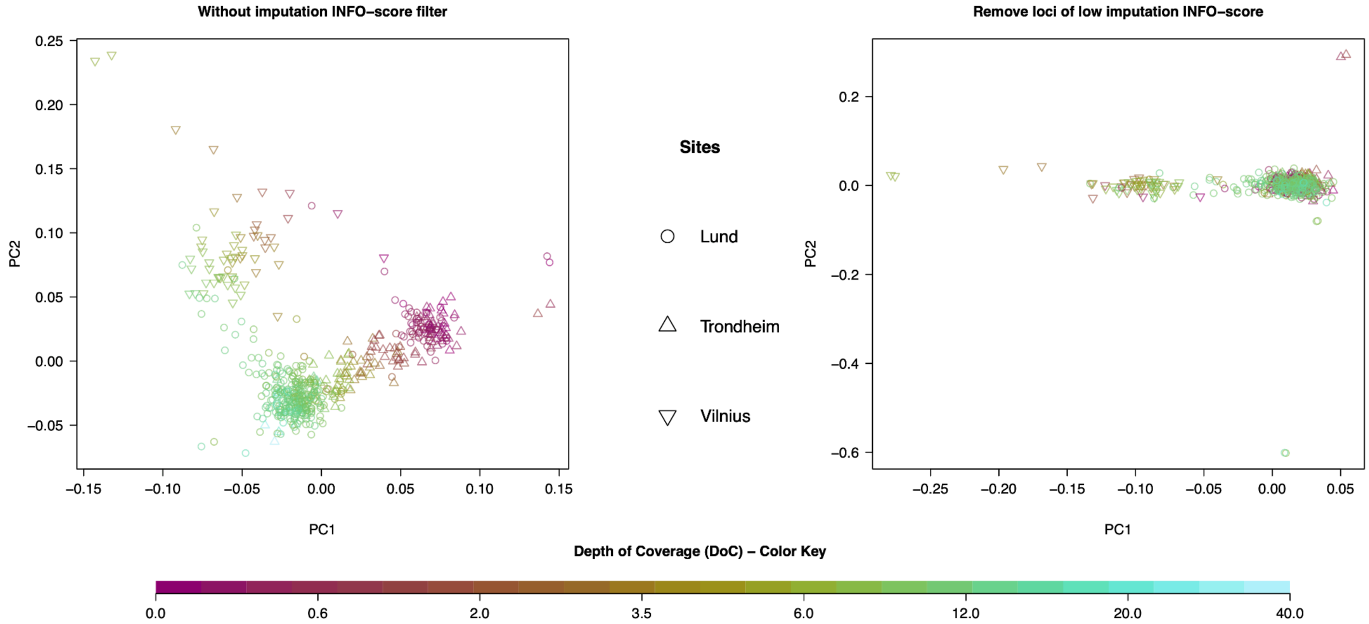


**Supplementary Figure 27.** Assessment of sequencing depth bias in principal component analysis (PCA). PCA of ancient individuals from Lund, Trondheim, and Vilnius was performed using common autosomal variants with minor allele frequency > 0.05. Shapes indicate sampling sites and colours indicate sequencing depth. **Left**: PCA performed without additional site filtering. PC1 primarily reflects differences in sequencing depth, indicating a depth-related bias that obscures population structure. **Right**: PCA after excluding variants with low imputation INFO scores (<0.95). This filtering eliminates the depth-driven effects and PC1 separates Vilnius individuals from individuals from the two Scandinavian sites: Lund and Trondheim.

### Supplementary Text

#### Supplementary Note 1: Contextual information

##### *Lund*

Although today located in southwestern Sweden, during the Middle Ages Lund was in the prosperous eastern part of the Danish kingdom. Founded in the 970s CE, it served as the archiepiscopal see of the Nordic countries from 1103. As Denmark's largest mint, Lund was also a hub of trade and commerce. However, the town's prominence began to wane in the 15th century as the ports of Malmö and Copenhagen expanded. Its decline was exacerbated by the Reformation in 1536, when most monasteries and churches were demolished.

Given Lund's establishment as a political and ecclesiastical center[1], it attracted clergy, royal officials, coiners, artisans, merchants, and both free and unfree labourers. Although written sources from its earliest period are scarce, extensive archaeological excavations since 1890 have revealed well-preserved cultural layers, offering detailed insights into settlement patterns and daily life. The archaeological record indicates that Lund did not evolve from an earlier village or trading post but was a planned foundation: plots were deliberately staked out, and the site was systematically populated. This provides a unique opportunity to study the city's earliest settlers.

The samples analysed in this study are principally teeth, obtained following ethical approval from the skeletal collection held at the museum of Kulturen in Lund. The skeletons were all excavated from the cemetery associated, over various periods, with the former Trinity Church, Trinity Abbey, and Trinity Parish Church, and which represents the largest medieval burial ground in Lund, in use ca 990-1536 CE[2]. The churches and large parts of the churchyard were excavated and archaeologically documented in connection with foundation excavations for new construction projects in 1974-75, and 1982-84[2]. Based on excavated areas and grave density, the churchyard is estimated to have contained approximately 5,700 individuals[3]. Archaeological evidence indicates that the cemetery was established following the construction in ca. 990 CE of the first wooden church in the middle of the site. This church was subsequently replaced by a large stone church at the southern part of the cemetery, which was consecrated to both 'Sanct Salvator' and 'Sancte Trinitas' in the 1050/60s CE[2]. This Trinity church is believed to have initially been the bishop's church, before the consecration of the current Cathedral in Lund at the beginning of the 12th century. In the second quarter of the 12th century the church was transformed into being both a parish as well as abbey church for the Order of Premonstratensians[4]. Already in the beginning of the 13th century the monastery was closed down[4], but the church maintained the status as a parish church. After a final rebuilding in the 13th century the Trinity was the largest church with an aisle-less nave in Scandinavia and the second most lavish of the city's churches during the High and Late Middle Ages[2]. At the Reformation in 1536 CE, the Trinity and most of the other churches were pulled down and the building stones were reused in the fortification of Malmo castle[5].

The establishment of the Trinitatis churchyard is mainly dated based on dendrochronological analysis of wooden coffins. The oldest coffin planks originated from a tree felled between 985 and 997 CE, and during the reign of King Sweyn Forkbeard (986-1014 CE)[1,5]. Combined evidence from stratigraphy, dendrochronology, radiocarbon dating, and characteristic arm positions of the deceased has enabled the division of graves into chronological groups: T1 (990–1050/60 CE) T2&3 (1050/60–1100 CE), T4 (1100- 1300 CE), and T5 (1300–1536 CE)[2]. Maria Cinthio[1] examined the latter period (T5) in greater stratigraphical detail and noted a general increase in funerary activity. This "red layer" also contained additional double and triple burials. This concentration is interpreted as evidence of mortality during the Black Death and subsequent plague outbreaks[3]. That study further highlights environmental and social changes following the pandemic: notably, an increase in average stature after 1350 CE, suggesting improved living standards and better access to nutritious food[3]. For this study, 61 individuals from T1, 101 from T2&3, 71 from T4, and 120 from T5 were sequenced. Of the T5 individuals, 42 derived from the layer associated with mortality during the Black Death and the subsequent plague outbreaks.

##### *Trondheim*

Trondheim, historically known as Nidaros for most of the Middle Ages, is situated on a peninsula encircled by the Nidelven River, with the exception of the western side, where it links to the mainland by a tiny isthmus. The Nidelven River emanates from the resource-abundant hinterlands near the Swedish border, providing access to furs, antlers, stone, and iron. The origins of the town's establishment are ambiguous; nevertheless, between 960-970 CE, perhaps under the influence of Ladejarlen (Earl of Lade) Håkon Sigurdsson, a regional commercial hub with a permanent population developed a little distance upstream from the river mouth[6]. At the start of the century, it was called Kaupangen (which translates to "trading place"), as referenced in the sagas, and had roughly two hundred permanent inhabitants; both the region and the population expanded swiftly to approximately 1200-1400 permanent people by circa 1100 CE. The foundation for this extensive development was the establishment of early Christian royal authority in the town, which included a residence along with administrative and economic functions, while the trading town emerged as a significant ecclesiastical centre with approximately 18 churches, following the canonisation of King Olav Haraldsson, who, as recounted in Snorri Sturluson's Olav Saga (from Heimskringla), perished for his Christian faith at the Battle of Stiklestad.

By around 1100 CE, the trade town experienced growth in size, population, productivity, and commercial development. The city subsequently saw the establishment of a new royal house and four to five churches, two of which were constructed of stone by King Harald Hardråde in the mid-11th century. Olav Kyrre (1067-1093 CE) initiated the establishment of bishoprics in Norway and constructed the inaugural bishop's church in Trondheim about 1070 CE. Norway's national saint and "eternal king," Olav the Holy Haraldsson, was interred near the high altar, attracting a growing influx of pilgrims. In 1153 CE, the city, then known as Nidaros, established itself as the centre of its own Norwegian ecclesiastical province, one of the largest in the Nordic nations throughout the Middle Ages. The entity had 10 dioceses, encompassing 5 located on Norwegian territory, as well as Greenland, the Orkney Islands, the Southern Isles including Man, and the two Icelandic dioceses of Skálholt and Holar; in 1202 CE, the Faroe Islands were also integrated.

The selected skeletons from Trondheim were excavated in four distinct churchyards, each reflecting a different environment and time era, ranging from medieval (FB, SG7-11, and SG4) to post-medieval/early modern Trondheim (West Front churchyard).

The FB (Folkebibliotekstomten) churchyard[7], from which 98 individuals were included in this study, surrounds an early stone church, not named but historically known as the St. Olav's church, constructed in the first half of the 12th century and renovated about 1200 CE. Located along Merchant's Street, in the oldest section of Trondheim, the church served the town's socioeconomic elite. It was likely preceded by an earlier wooden church. The churchyard has been excavated archaeologically several times, the most recent being in 1984. A total of 200 square meters were unearthed, and 389 skeletons identified. The churchyard is divided into three distinct phases: 1100/1150-1175 CE, followed by 1175-1500 CE and 1500-1600 CE, although each of these phases can be subdivided. During the second phase, the churchyard was extended northward, replacing an earlier town yard. Around the second half of the 13th century, the church was given over to the Franciscans. While some of the latest burials in Phase 2, and most of the burials in Phase 3, are believed to be Franciscan monks, wealthier urban residents may still have been buried here. Late in Phase 3 (1500-1699 CE), the graveyard was separated from the church itself, most likely due to a shift to private ownership after the Reformation in 1536 CE, and the land put to other uses.

The SG4 (Søndre gate 4) churchyard[8], from which 16 individuals were included in this study, surrounds a stone church that was built at some point during the early 12th century. The church was previously believed to have been the St. Gregory's Church referred to in saga literature, although it is now debated whether it was instead dedicated to St. Olav. Upon completion, it stood on the outskirts of the town, on Long Street, an area dominated by craftspeople and workshops. It might have been built for ceremonial reasons relating to the devotion of St. Olav as a station church, with ties to the royal family and the bishop, but it likely also served as a parish church. The building had both a narthex and a crypt, uncommon architectural elements in medieval Norwegian church architecture. The choir was rebuilt ca.1200 CE. The church burned down in the first part of the 16th century and was thereafter utilized as a stone quarry.

The SG7-11 (Søndre gate 7-11) church[9], from which 10 individuals were included in this study, was located along Merchant's Street, in the medieval town's elite quarter (similar to the FB church). Built around 1100 CE, its wooden construction was unusual for its time, and it may have served an important symbolic role in the cult of St. Olav. The church was rebuilt four times before being demolished in the second half of the 14th century. A large number of extremely young infants were buried near the altar, which remained undisturbed under an open shelter after the destruction.

The West Front churchyard[10] from which 13 individuals were included in this study, contains burials that were located in the courtyard on the west side of Nidaros Cathedral and excavated in 1996 in advance of improvements to the area. Historical evidence indicates that the site was in use as a churchyard by 1663 CE. However, in the wake of the Reformation, Nidaros had been downgraded to parish church status and the churchyard may have been in use as early as 1585 CE. The churchyard remained in use until 1897 CE, when it was officially closed. Six phases of burial activity were identified. The West Front individuals included in this study represent five of these phases. Phase 21, the earliest phase of activity, is difficult to date. The excavators suggest the early 18th century, although radiocarbon results for the single individual in this study associated with this phase (WF737), may suggest an earlier date. Phase 22 dates to the late 18th/early 19th century and represents the most intense period of activity. Phase 23 dates to the early 19th century, seemingly after 1806 CE. Phase 25 represents reinterment of individuals previously interred in burial vaults within the church building. As such, these likely represent higher status individuals than those in the other phases of the West Front churchyard. Burials within church buildings were outlawed in Norway in 1805 CE, so while the Phase 25 burials may represent a wider time range than the other churchyard phases, they must all pre-date 1805 CE. Phase 26 represents the final phase of activity and must pre-date 1897 CE.

*Vilnius*

Christianization and urbanisation occurred later in Lithuania than in other European regions, shaping distinct political, technological, and social developments. Positioned between Catholic and Orthodox Europe, the Grand Duchy of Lithuania (GDL) developed a unique medieval path: from the 13th century onward, it incorporated new territories, and Vilnius, situated on the south-east margin of present-day Lithuania, emerged as a multicultural centre of a much larger territory, attracting people from across Europe[11]. The presence of early Christian settlements in a predominantly pagan, 13th century town marked a key phase in Vilnius' premodern history. Subsequent milestones – Christianization and state growth in the 15th century, prosperity during the 16th-century (the "Golden Age"), and decline in the mid-17th-century Muscovite invasion ("the Deluge") – shaped urban development as a multiethnic and multiconfessional capital city (mostly Roman Catholic and Eastern Orthodox, but with significant presence of other Christian confessions, as well as Judaism and Islam)[12]. It served as an administrative centre (one of two capitals of the Polish-Lithuanian Commonwealth) and as the centre of crafts and trade.

The samples from Vilnius are principally temporal bones, supplemented by a small number of teeth, collected after formal approval from the specialised storage facility for bioarcheological materials at the Department of Anatomy, Histology and Anthropology, of the Faculty of Medicine, Vilnius University. Samples derive from three distinct sites.

Bokšto str. 6 is one of the oldest cemeteries in Vilnius, having been radiocarbon dated to between the beginning of 13th – early 15th centuries. From this site, 19 individuals were sequenced. The burial site, located in the eastern half of Vilnius Old Town, within the area historically known as *Civitas Rutenica* (the Ruthenian town), represents the earliest known burial ground in Vilnius. This burial site is unique in the context of Vilnius's early urban history, not only for its chronological primacy, but also due to the Christian character of the burials during a time when Vilnius was still predominantly pagan. Archaeological and historical evidence suggests that the cemetery served the Orthodox Christian community, most likely composed of immigrants and their descendants, possibly from Ruthenian or other East Slavic backgrounds, as well as local converts. Extensive excavations were conducted during multiple field seasons (2006–2007, 2009–2011, 2012, and 2014), resulting in the discovery of 533 inhumations[13].

Aguonų str.10. Historical records and old maps of Vilnius do not indicate the presence of a cemetery at this site, that is situated outside the medieval city walls, and was found unexpectedly during construction work. Salvage excavations took place in 2006 and 2007. During both seasons the burial site revealed a total of 119 individual and group graves, containing the remains of approximately 220 individuals. Only a very small number of burials were found within coffins, and numerous graves contained individuals buried in unusual positions, such as upside down or with limbs positioned erratically, and in different orientations (including northwest, south, southeast, northeast, north, and east). Moreover, 36 group graves were found, containing up to 15 individuals per pit. Based on grave goods recovered, this burial location's use has been dated to the 15th to 16th centuries[14,15], subsequently confirmed by radiocarbon dating[16]. Anthropological analysis found no skeletal indicators of trauma or specific disease. However, ancient DNA (aDNA) analysis has detected convincing signals of ancient *Y. pestis* and presence of *Treponema pallidum pertenue*, the bacterium responsible for yaws[16], demonstrating that aDNA can identify infectious diseases in cases where skeletal evidence alone cannot. 20 individuals from Aguonų str.10 were sequenced in this study.

The Subačiaus str.7 burial ground. Under a ruler's privilege granted in 1592 CE, the Orthodox community of Vilnius established a church in the southern part of the city in 1597. The first structure on the site was a wooden church, which was later replaced by a masonry building (the Church of the Holy Spirit) approximately 27 meters uphill from the cemetery. Archaeological investigations conducted in 1998 uncovered a cemetery dating to the 17th century. A total of 131 burials were documented. The cemetery was situated on the ruins of a building of the fifteenth-sixteenth centuries. The burials were arranged in an orderly fashion—grouped in rows that may have been influenced by existing paths, fences, or buildings. Coins (Polish shillings) from the first half of the 17th century were found in several graves, along with two Orthodox bone rosaries[17]. For this study, 19 individuals from the Subačiaus str.7 burial ground were sequenced.

##### Supplementary Note 2: Genome-wide signals in GWAS

In the meta-analysis comparing allele frequencies between pre-BD and present-day individuals, we found two genome-wide significant SNP signals. The first SNP signal (chr2:160798465) is located in an intergenic region on chromosome 2 (meta-analysis $P$=8.50×$10^{-9}$, Lund $P$=2.45×$10^{-6}$, Trondheim $P$=4.41×$10^{-5}$, with the allele frequency changing in the same direction in Lund and Trondheim). The second SNP signal (chr4:139027159) lies within intron 1 of the gene *NOCT* (meta-analysis $P$=6.74×$10^{-43}$, Lund $P$=3.08×$10^{-11}$, Trondheim $P$=3.74×$10^{-28}$, the same direction in Lund and Trondheim). However, both variants were also highly significant when comparing post-BD to present-day individuals (chr2 meta-analysis $P$=5.68×$10^{-9}$, chr4 meta-analysis P=7.76×$10^{-75}$), and showed no evidence of significant allele frequency change between pre-BD and post-BD individuals even using a nominal significance threshold of 0.05 (chr2 meta-analysis $P$=0.48, chr4 meta-analysis $P$=0.82). In addition, we detected a genome-wide significant indel variant on chromosome 3 (chr3:171104061), corresponding to a deletion in the *TNIK* gene (meta-analysis $P$=9.99×$10^{-10}$, Lund $P$=2.05×$10^{-19}$, Trondheim $P$=0.84, opposite directions in Lund and Trondheim), with the signal largely driven by Lund. However, similar to the two SNP signals, this variant was also highly significant when comparing post-BD to present-day individuals in Lund ($P$=2.84×$10^{-20}$), and showed no evidence of significant allele frequency change between pre-BD and post-BD individuals using a nominal significance threshold of 0.05 (meta-analysis $P$=0.85, Lund $P$=0.85). Importantly, for all three loci, we observed associations for only a single variant after our stringent site-level filtering (see Methods) in the surrounding ±500 kb region (Supplementary Fig. 12). At the chromosome 2 locus, many nearby variants in high LD with the lead variant in present-day individuals exhibit almost no frequency change. At the chromosome 3 and chromosome 4 loci, lead variants are in very weak LD with nearby variants. Hence, there is no indication that other variants on shared haplotype backgrounds at these three loci also rose in frequency. Together, the significant frequency differences between the post-BD and present-day individuals, combined with the lack of significant signals in the pre-BD vs post-BD comparison and the lack of neighboring SNPs with a signal, make it unlikely that the observed allele frequency changes can be attributed to the Black Death.

In addition to the three genome-wide significant signals in the meta-analysis, several genome-wide significant loci were detected in the single-site GWAS analyses comparing pre-BD to present-day individuals. Two were found in Lund: an intergenic variant on chromosome 1 (chr1:225574884, $P$=1.03×$10^{-8}$), and variants within the T-cell receptor alpha gene segments in the q11.2 region on chromosome 14 (*TCRA*, chr14:22193177, $P$=3.02×$10^{-12}$). The signal on chromosome 1 exhibited the same issues as the meta-analysis-based signals: we observed a significant association when we compared post-BD to present-day (chr1:225574884 $P$=1.42×$10^{-6}$); it showed no evidence of significant allele frequency change between pre-BD and post-BD individuals (Lund $P$=0.85); and we observed an association for only one SNP in the region after quality filtering when comparing pre-BD to present-day (Supplementary Fig. 12). Hence, it is also unlikely that the observed allele frequency change in this locus can be attributed to the Black Death. The signal on chromosome 14 appeared more convincing: after quality filtering it consisted of multiple neighbouring markers that showed similarly low $P$-values (Supplementary Fig. 12) and the $P$ value for pre-BD vs post-BD comparison was marginally significant (Lund $P$= 0.03). Furthermore, while there was a significant association signal when comparing post-BD vs present-day ($P$=3.70×$10^{-4}$), that association was less strong than for the other signals, and this signal might reflect other episodes of selection given the established role of the T-cell gene segments in the region in adaptive immunity and immune response to many bacterial pathogens. Recurrent or long-term selection on immune-related variation in this region might have resulted in allele-frequency differences between medieval and present-day individuals. Finally, while the Trondheim individuals did not show a significant change in allele frequency when comparing pre-BD to present-day ($P$=0.07), the estimated allele frequencies at that site are in the same direction (Supplementary Table 6a): pre-BD, 0.18, (95% CI: 0.13-0.24); present-day, 0.27 (95% CI: 0.25-0.29). In light of this, this signal merits further follow up. However, we caution that this locus is in a region that is difficult to map to and therefore we do not consider it a strong candidate.

To summarise, neither the meta-analyses nor the single-site GWAS produced convincing signals of selection. We note that all statistically significant associations are from comparisons between ancient and present-day individuals. Furthermore, with the exception of the *TCRA* locus, they consist of lone hits without any tagging variants, and are not replicated in the pre-BD versus post-BD GWAS. As such, they are more likely to reflect technical differences in the data generation of ancient versus present-day genotypes rather than genuine allele frequency changes driven by the Black Death. Mismapping of short, damaged ancient DNA reads and imputation errors at individual loci in the imputed ancient genotypes are plausible explanations. To assess this, we calculated quality metrics on the aligned BAM files and imputed genotypes of securely dated ancient individuals from Lund and Trondheim. The results (Supplementary Table 6b) suggest potential mismapping around all of the significant loci: the chromosome 1, 2, and 4 loci are affected by low average mapping quality, the chromosome 3 locus has low average depth, and the chromosome 14 loci are characterised by Hardy-Weinberg deviations in imputed ancient genotypes at $P{\sim}4\times10^{-5}$. Uncertainty between pipelines in mapping and calling at some of these regions is also suggested by gnomAD v4.1.1[18], where the chromosome 1 and 4 variants are assigned a frequency of zero due to the removal of many low-quality genotypes, and the chromosome 3 locus carries two quality flags and is described as having a much lower alternative allele frequency in non-Finnish Europeans (51%) than the frequency that we observe in our present-day Scandinavians (100%). Despite our stringent quality control of variants and genotypes (Methods), it is possible that some problematic loci were not excluded and aDNA imputation was not able to rectify these underlying issues. Inspection of the regional association plots before site-level filtering further supports this possibility (Supplementary Fig. 12B): most of the significant signals are within narrow clusters of variants that failed our quality filters. Additional details and annotations, together with allele frequencies in the different cohorts, can be found in Supplementary Table 6.

#### Supplementary Note 3: Evaluation of ancestry assignment methods based on leave-one-out tests

*Method*

To assess the robustness of our ancestry classification framework, which integrates PCA and IBD analyses, we performed leave-one-out cross-validation on the present-day populations from the reference data.

For the West Eurasian PCA, only populations with a minimum sample size of 40 were included. In each iteration, one individual from each included population was held out and tested against all included reference populations using chi-square tests based on PCA-derived Mahalanobis distances ($D_M$) (see Methods). We then assessed whether the held-out individual was statistically inconsistent with the distributions of the targeted reference populations ($P<0.001$). To specifically evaluate the performance of the PCA-based classification for ancient individuals from Lund and Trondheim, we used Danish, Swedish and Norwegian populations as Scandinavian references. For the ancient individuals from Vilnius, Lithuanian and Estonian populations were used as Baltic references; Latvia was excluded due to small sample size (N=7). We then calculated the proportion of held-out individuals not rejected as Scandinavian or Baltic, respectively for each tested population.

We performed the same leave-one-out analysis for the Scandinavian PCA, which is constructed using present-day individuals (N=17,526) from 14 regions across Scandinavia in the OmniExpress dataset. In this analysis, we used individuals from the populations of Denmark and Skåne (Sweden) as local references for ancient individuals from Lund, and used individuals from the population of Trøndelag (Norway) as local references for ancient individuals from Trondheim. For each tested population, we calculated the proportion of held-out individuals consistent with present-day populations of Denmark or Skåne, and consistent with the present-day population of Trøndelag, respectively.

We next evaluated performance of the IBD-based analysis using a leave-one-out procedure conducted at two levels. First, using the OmniExpress European reference dataset, we assessed the ability of IBD sharing to distinguish Scandinavian from non-Scandinavian ancestry. We included the same 14 Scandinavian populations used in the Scandinavian PCA and several non-Scandinavian populations from neighbouring regions: Icelandic, German, Dutch, Orcadian, and Shetlandic, with the latter two used as proxies for British-Irish ancestry. Due to the computational cost of IBD fragment calling for all pairs of individuals from this large reference set of present-day individuals, we randomly selected 100 individuals from each population and called IBD fragments against all remaining samples for the leave-one-out test. Individuals were classified as ‘non-Scandinavian’ when they showed significantly greater IBD sharing with any non-Scandinavian population than with all Scandinavian populations using Welch’s t-test ($P$-value $< 0.05$, see Methods). For each tested population, we reported the proportion of individuals flagged as ‘Scandinavian’.

At the second level, we assessed the resolution of IBD sharing for distinguishing ancestry within Scandinavia. The test was restricted to the 14 Scandinavian populations, using the same randomly selected 100 individuals per population as above. As in the Scandinavian PCA, the populations of Denmark and Skåne County (Sweden) were used to define the local reference for the ancient individuals from Lund, and the population of Trøndelag (Norway) was used as the local reference for ancient individuals from Trondheim. Individuals were classified as 'non-local' when sharing significantly greater IBD with any reference population other than the local. For each tested population, we calculated the proportion of individuals flagged as 'local'.

| **Tested population** | **Proportion not rejected as Scandinavian** | **Proportion not rejected as Baltic** |
|---|---:|---:|
| Danish | 1.00 | 0.00 |
| Swedish | 0.99 | 0.00 |
| Norwegian | 1.00 | 0.00 |
| German | 0.18 | 0.01 |
| Dutch | 0.70 | 0.00 |
| Belgian | 0.05 | 0.00 |
| English | 0.41 | 0.00 |
| Irish | 0.00 | 0.00 |
| Scottish | 0.07 | 0.00 |
| Welsh | 0.12 | 0.00 |
| Northern_Irish | 0.01 | 0.00 |
| French | 0.00 | 0.00 |
| Polish | 0.00 | 0.48 |
| Russian | 0.00 | 0.49 |
| Czech | 0.00 | 0.0 |
| Estonian | 0.00 | 1.00 |
| Lithuanian | 0.00 | 0.99 |
| Italian | 0.00 | 0.00 |
| Spanish | 0.00 | 0.00 |
| Finnish | 0.01 | 0.02 |
| Croatian | 0.00 | 0.00 |
| Greek | 0.00 | 0.00 |
| Turkish | 0.00 | 0.00 |
| Icelandic | 0.00 | 0.00 |

**Supplementary Table 11.** Rates of non-rejection for present-day populations in the West Eurasian reference set against Scandinavian and Baltic distributions in leave-one-out tests of the $D_M$ method for PCA.

*Results*

The leave-one-out tests indicate that the West Eurasian PCA-based analysis has good power to distinguish Scandinavian populations from other European populations (Table S11). The three Scandinavian populations show very high classification accuracy, with more than 99% of individuals correctly classified as Scandinavian. However, several populations are difficult to distinguish from Scandinavians, with non-rejection rates greater than 10%, including Dutch,

German and British populations (English and Welsh). Of these, the Dutch population is the most challenging to differentiate, with approximately 70% of individuals not rejected as Scandinavian.

The West Eurasian PCA-based analysis also provides good power to differentiate Baltic populations and most other European populations, with more than 99% of Lithuanian and Estonian individuals correctly classified as Baltic. While most other European populations show low non-rejection rates for Baltic ancestry (<2%), Eastern European populations of Slavic ancestry, namely, Polish and Russian are difficult to distinguish from Baltic populations, with non-rejection rates of 48-49%.

The Scandinavian PCA-based analysis provides limited power to differentiate the more fine-scale differences in ancestry within Scandinavia (Table S12). For the Lund local reference, 98.7% of Danish individuals and 95.6% of Skåne individuals are not rejected by the Denmark-Skåne reference. Norwegian individuals are generally distinguishable from this reference, with low non-rejection rates ranging from 1.2% to 13.7%. In contrast, individuals from other Swedish regions have much higher non-rejection rates against the Denmark-Skåne reference, ranging from 31.8% in northern Sweden to 94.4% in southern Sweden. For the Trondheim local reference, 94.8% of Trøndelag individuals are not rejected by the Trøndelag reference. Individuals from the Danish and Swedish populations have lower non-rejection rates against this reference, ranging from 14.9% in southern Sweden to 47.6% in Denmark. In contrast, individuals from other Norwegian regions have consistently higher non-rejection rates, ranging from 42.9% in southern Norway to 82.3% in Oslo. Not surprisingly, these results indicate that, although the Scandinavian PCA can capture broad genetic structure between countries, it has more limited resolution for distinguishing fine-scale regional ancestry variation within countries.

| **Tested population** | | **Proportion not rejected as local (Denmark + SE:Skåne)** | **Proportion not rejected as local (NO:Trondelag)** |
|---|---|---|---|
| **Country** | **Region** | | |
| Denmark | Denmark | 0.99 | 0.48 |
| Sweden | Skåne | 0.96 | 0.31 |
| Sweden | South | 0.94 | 0.15 |
| Sweden | Gotaland | 0.85 | 0.33 |
| Sweden | Stockholm | 0.82 | 0.29 |
| Sweden | Central | 0.73 | 0.29 |
| Sweden | North | 0.32 | 0.15 |
| Norway | Oslo | 0.14 | 0.82 |
| Norway | South | 0.02 | 0.43 |
| Norway | West | 0.01 | 0.73 |
| Norway | Trøndelag | 0.03 | 0.95 |
| Norway | East | 0.11 | 0.82 |
| Norway | Central | 0.07 | 0.79 |
| Norway | North | 0.04 | 0.77 |

**Supplementary Table 12.** Rates of non-rejection for present-day populations in the OmniExpress Scandinavian reference set against the local reference distributions for Lund (Denmark and SE_Skåne) and Trondheim (NO_Trøndelag) in leave-one-out tests of the $D_M$ method for PCA.

The IBD-based leave-one-out tests show good performance in distinguishing between Scandinavian populations and closely related non-Scandinavian populations, with the notable exception of Germany (Table S13). Similar to the West Eurasian PCA, all Scandinavian populations have very high self-classification rates (>98%). The IBD-based analysis also effectively distinguishes Icelandic, British-Irish (represented by Orcadian and Shetlandic), and Dutch ancestry from Scandinavian ancestry. Notably, the non-rejection rate for the Dutch population decreases from 70% in the PCA-based analysis to 4% in the IBD-based analysis. However, the IBD-based method has very limited power to distinguish German ancestry from Scandinavian ancestry, with 91% of German individuals not rejected as Scandinavian, compared to 18% in the PCA-based analysis. Several factors likely contribute to this reduced power to distinguish German from Scandinavian ancestry. First, the sampling of present-day German individuals is relatively sparse in our IBD dataset (~1000 German individuals representing a population of ~83 million, compared to ~6000 Dutch individuals representing a population of 18 million). This reduces the probability of detecting long IBD fragments shared with distant relatives. Second, the German population likely had a larger effective population size over the last 30 generations than the other populations included in analysis[19], as reflected by the relatively low IBD sharing observed between Germans in our data. Third, northern German populations are known to share substantial admixture with Danish and southern Scandinavian populations. Fourth, our IBD-based test is conservative in identifying non-Scandinavian ancestry, because individuals are rejected as Scandinavian only when they share significantly more IBD with a non-Scandinavian reference population than with Scandinavian reference populations. Although most German individuals in the test set (randomly selected, N=100) have the greatest amount of IBD sharing with other German individuals, the amount of sharing is not significantly greater than their IBD sharing with Danes (Supplementary Fig. 28). Overall, the leave-one-out tests show that the IBD-based analysis performs well for most populations, but appears overly conservative in distinguishing German ancestry from Scandinavian ancestry.

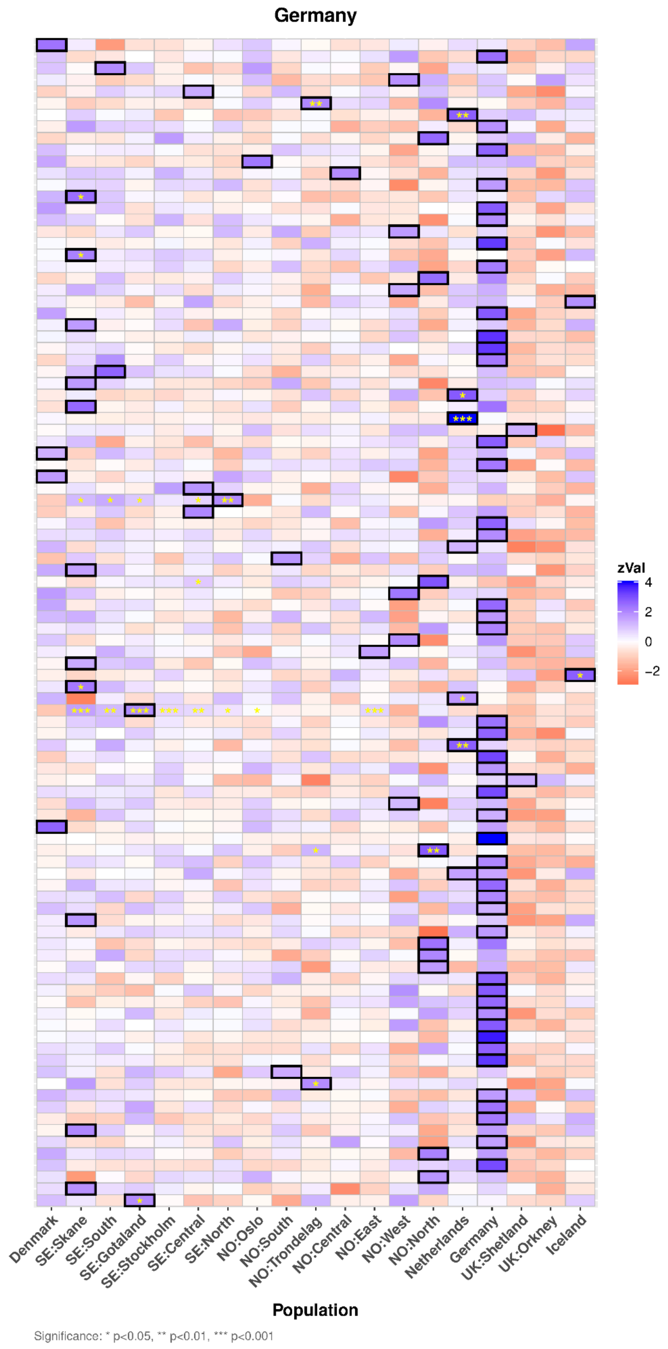


**Supplementary Figure 28. Heatmap of IBD sharing between 100 randomly selected present-day German individuals and reference populations.** Each row represents a present-day individual, and columns represent reference populations from the OmniExpress European datasets. IBD sharing was estimated using a leave-one-out procedure. For each individual, the average amount of IBD shared with each reference population was standardized across populations. The reference population with the highest standardized IBD sharing for each individual is indicated by a thick black outline. Significance markers indicate cases where an individual shares significantly more IBD with a reference population than with German reference population based on a one-sided Welch's t-test. Asterisks denote significance levels: $p<0.05$ (*), $p<0.01$ (**), and $p<0.001$ (***).

Given the limited power of our IBD-based analysis to distinguish German from Scandinavian ancestry, some ancient individuals classified as Scandinavian may have predominantly German ancestry. To address this possibility, we performed an additional IBD-based test. Specifically, we tested whether the ancient individuals classified as Scandinavian had significantly greater IBD sharing with at least one Scandinavian reference population than with the German reference population. This test was based on the expectation that individuals of predominantly German ancestry should not share significantly more IBD with any Scandinavian reference population than with the German reference population. Using this approach, 202 out of 206 ancient individuals from Lund classified as Scandinavian have significantly higher IBD sharing with at least one Scandinavian population than with the German reference population (Welch's *t*-test, $P<0.01$), as do all 109 ancient individuals classified as Scandinavian from Trondheim. By comparison, only four of out of 100 German individuals in the leave-one-out dataset show significantly greater IBD sharing with any Scandinavian population than with the German reference population.Together with the relatively low non-rejection rate (17.5%) of Germans as Scandinavian in the PCA-based analysis, these results suggest that the risk of misclassification of ancient individuals with predominantly German ancestry as Scandinavian is low.

| **Tested population** | **Proportion not rejected as Scandinavian** |
|---|---|
| Denmark | 0.99 |
| SE:Skåne | 1.00 |
| SE:South | 0.99 |
| SE:Gotaland | 1.00 |
| SE:Stockholm | 1.00 |
| SE:Central | 1.00 |
| SE:North | 1.00 |
| NO:Oslo | 0.99 |
| NO:South | 1.00 |
| NO:West | 1.00 |
| NO:Trøndelag | 1.00 |
| NO:East | 0.98 |
| NO:Central | 1.00 |
| NO:North | 1.00 |
| Germany | 0.91 |
| Netherlands | 0.04 |
| Orkney | 0.00 |
| Shetland | 0.00 |
| Iceland | 0.00 |

**Supplementary Table 13.** Rates of non-rejection as Scandinavian for present-day populations in the OmniExpress reference set in leave-one-out tests of the IBD method.

Last, and importantly, the IBD-based analysis shows remarkably improved power to differentiate ancestry within Scandinavia compared with PCA (Table S14). For the Lund local reference, 98% of Danish individuals and 81% of Skåne individuals are not rejected by the Denmark-Skåne reference. Although these self-classification rates are slightly lower than those observed in the PCA-based analysis, the IBD-based analysis more clearly distinguishes individuals from the local reference populations and those from other Scandinavian regions. Specifically, Norwegian populations can be distinguished from the Denmark-Skåne reference with very low non-rejection rates (0-2%). Individuals from most other Swedish regions show low to modest non-rejection rates (1-14%), with the highest non-rejection rate (28%) observed in southern Sweden. For the Trondheim local reference, 82% of Trøndelag individuals are not rejected by the Trøndelag reference, which is lower than in the PCA-based analysis. However, compared with PCA, the IBD-based analysis provides much clearer separation between the Trøndelag reference and other Scandinavian populations. Individuals from Denmark and Swedish regions have very low non-rejection rates (0-4%), whereas individuals from other Norwegian regions exhibit higher but moderate non-rejection rates (8-22%). These results indicate that IBD provides substantially improved power to resolve fine-scale population structure within Scandinavia compared with PCA.

| Tested population | | Proportion not rejected as local (Denmark + SE:Skåne) | Proportion not rejected as local (NO:Trøndelag) |
|---|---|---|---|
| Country | Region | | |
| Denmark | Denmark | 0.98 | 0.01 |
| Sweden | SE:Skåne | 0.81 | 0.00 |
| Sweden | SE:South | 0.28 | 0.00 |
| Sweden | SE:Gotaland | 0.10 | 0.03 |
| Sweden | SE:Stockholm | 0.14 | 0.00 |
| Sweden | SE:Central | 0.06 | 0.01 |
| Sweden | SE:North | 0.03 | 0.04 |
| Norway | NO:Oslo | 0.01 | 0.19 |
| Norway | NO:South | 0.00 | 0.08 |
| Norway | NO:West | 0.00 | 0.16 |
| Norway | NO:Trøndelag | 0.01 | 0.82 |
| Norway | NO:East | 0.01 | 0.18 |
| Norway | NO:Central | 0.02 | 0.16 |
| Norway | NO:North | 0.01 | 0.22 |

**Supplementary Table 14.** Rates of non-rejection for present-day populations in the OmniExpress Scandinavian reference set against the local populations used for Lund (Denmark and SE_Skåne) and Trondheim (NO_Trøndelag) in leave-one-out tests of the IBD method.

#### Supplementary Note 4: Comparison of ancestry assignment methods in ancient individuals

##### *Data and approach*

Supplementary Note 3 evaluates our PCA-based and IBD-based ancestry classification methods in present-day individuals, and focuses on defining outliers without explicitly attempting to identify their region of origin. In this note, we describe the use of PCA-based Mahalanobis distances ($D_M$) and rates of long IBD sharing with present-day references for identifying potential source regions. The results from these approaches are described in Supplementary Note 5 and summarised in the main text ("Assessing the origin of outliers and non-locals"). In this note, we compare these methods to each other and to qpAdm[20], a method that is often used in ancient DNA studies. Importantly, the questions addressed by these methods are related, but not directly equivalent. PCA-based $D_M$ measures to what extent individuals are consistent with a distribution in PCA space, long IBD sharing measures relatively recent relatedness, while qpAdm formally tests admixture models. Nonetheless, we find the comparison useful because we expect many readers to be more familiar with qpAdm than our $D_M$- and IBD- based approaches.

We use the Vilnius assemblage for this comparison for several reasons. Firstly, it represents a tractable test case, comprising 57 individuals exhibiting modest diversity in ancestry (Fig. 3A, Supplementary Table 10) which derives from well-characterised Baltic- and Slavic-associated sources[21,22]. Secondly, for Vilnius, the IBD reference data comprises an entirely different set of individuals (UK Biobank) than the West Eurasian reference data used for PCA-based Mahalanobis distances ($D_M$), so IBD results represent an independent evaluation.

##### *Methods*

First, we performed ancestry assessments using the PCA-based $D_M$ method. In other words, in this analysis an ancient individual was considered statistically consistent with the distribution for a present-day group when the squared $D_M$ value did not exceed the chi-square threshold (18.5) corresponding to $P \geq 0.01$. Due to stable population structure in much of Europe over the past 1000 years[23], consistency with a reference distribution can be interpreted as suggesting an origin in that region. However, we acknowledge that the PCA-based $D_M$ method is not a formal ancestry model using contemporary sources. We employed a different threshold for this "rule-in" ancestry assignment compared to the "rule-out" threshold of $P < 0.001$ used in the outlier detection procedure (Methods) for several reasons. Firstly, our sampling locations provide a strong geographic prior that individuals have ancestry typical of that region, and we require strong evidence to assign individuals as outliers. A threshold of $P < 0.001$ for ruling out consistency with the local distribution therefore represents a conservative criterion for outlier detection, whereas using $P \geq 0.001$ as a rule-in criterion for ancestry assignment would be too lax. Secondly, the more conservative threshold of $P < 0.001$ for defining outliers accommodates minor ancestry shifts between ancient populations and the present-day proxies used as references.

Next, for qpAdm, we ran two sets of models. The first used present-day genotypes from the West Eurasian PCA, with a source selection designed to be similar to the reference distributions we selected for the $D_M$ analysis. The second used shotgun sequenced ancient sources from the

AADR v62.0[24]. Here, we selected populations that broadly covered West Eurasia, generally slightly preceded our test populations, and were evaluated as independent ($P$<0.01) by qpWave. While the use of present-day populations as sources violates some assumptions of qpAdm, this approach is analogous to our PCA- and IBD-based methods and has been explored in previous studies[25]. To maximize both sample size and number of genotypes, we merged putatively related populations based on iterated qpWave analysis. Candidates for grouping for which qpAdm rejected cladality with any other group members $P$<0.01 were dropped. This process was iterated until all populations showed $P$≥0.01 within-group and $P$<0.01 between groups (Table S15). In both sets of qpAdm models, we ran a rotating set-up[20] using ADMIXTOOLS 2[26] with all sources in the leftright (competing sources) position. However, in the ancient run we placed China_YR_MN.SG in the rightfix (fixed outgroup) position, as including it in leftright led to unexpected passing models of China_YR_MN.SG and Russia_BolshoyOleniyOstrov_MBA.SG for many individuals, presumably because they are symmetrical outgroups to many test individuals. For the ancient-sources set, we restricted Vilnius genotypes to variants with MAF>0.05 in the reference panel used for imputation, merged it with AADR v62.0 data restricted to the defined source populations, and ran admixtools::extract_f2(maxmiss = 0.2, afprod = TRUE, blgsize = 0.05) (blocksize 5 cM) on the merged data to generate allele-frequency products to use as input. For the present-day sources, we used the West Eurasian reference (see Methods) and geno_to_afprods(poly_only_ap = FALSE, blgsize = 1000000) (blocksize 1 Mb) to generate allele-frequency products.

Ancestry assignment to specific sources based on IBD results was also considered but proved to be non-trivial. IBD sharing intensity is strongly shaped by the demographic history of the populations analysed[27]: a bottlenecked population with low haplotype diversity induces strong IBD sharing between all populations that receive ancestry from that source, which, depending on bottleneck intensity, can be clearly observed even when the proportion of ancestry received is small. Conversely, a source population with high haplotype diversity may induce only weak levels of IBD sharing between descendants. This explains a surprising observation in the IBD results for the Vilnius assemblage: of the individuals that we model as having fully Slavic-associated ancestry, most (62% to 73%, depending on method) show highest IBD sharing with present-day Baltic populations, not present-day Slavs (Supplementary Fig. 21, Supplementary Fig. 30). Importantly, the same observation is reported in a recent study on the origins of Slavs[22], in which a majority of ancient Slavic-associated individuals were found to share more IBD with present-day Baltic populations than Slavic ones. We believe that this is because Slavic populations derive substantial ancestry from a source maximised in present-day Baltic populations (the standard model for Slavic origins[22]), and this Baltic-associated ancestry source had low haplotype diversity, thus generating a strong IBD signal. Additionally, any Slavic-specific bottleneck was not intense enough to generate a similarly strong Slavic-associated IBD signal.

This is an example of a general phenomenon which complicates the interpretation of IBD sharing results. However, we do not believe that it is a significant concern for our study. Firstly, Baltic- and Slavic-associated ancestry can be distinguished from their broader profiles of IBD sharing (see below and Supplementary Fig. 30). Secondly, the results of our IBD-based procedure for defining outliers generally agree with the PCA-based method. Thirdly, a major focus of our analyses is Scandinavian ancestry, and, unlike in the Baltic/Slavic case, there does not appear to exist a present-day non-Scandinavian population that displays more IBD sharing with ancient Scandinavians than present-day Scandinavians. Finally, in our characterisation of the diversity of ancestry at each site (Supplementary Note 5), we mainly use IBD as a validation method to support PCA $D_M$-based assignments, with some exceptions in clear-cut cases like LUN114 (Supplementary Notes 5, 7, main text "Assessing the origin of outliers and non-locals").

Present-day qpAdm sources:

English, Dutch, Irish, German, French, Danish, Hungarian, Swedish, Spanish, Ukrainian, Bulgarian, Finnish, Italian, Latvian, Greek, Lebanese_Christian

| **Ancient source** | **Constituent populations** |
|---|---|
| China_YR_MN.SG | China_YR_MN.SG |
| Lithuania_IA.SG | Lithuania_Marvele_Roman.SG,Lithuania_Bailuliai_BarrowCulture.SG |
| Lebanon_LateAntiquity_and_Medieval.SG | Lebanon_ElJaouze_Phoenician.SG,Lebanon_Medieval.SG |
| Scandinavia_IA.SG | Denmark_IA.SG,Sweden_IA_2.SG,Estonia_EarlyViking.SG |
| Portugal_MonteDaNora_LateRoman.SG | Portugal_MonteDaNora_LateRoman.SG |
| Croatia_Slovenia_LateAntiquity.SG | Croatia_Zadar_Roman.SG,Slovenia_Emona_Roman.SG |
| Poland_Slavic_IA.SG | Poland_Niemcza_IA.SG,Poland_Santok_IA.SG |
| Armenia_Beniamin_EarlyMedieval.SG | Armenia_Beniamin_EarlyMedieval.SG |
| Russia_BolshoyOleniyOstrov_MBA.SG | Russia_BolshoyOleniyOstrov_MBA.SG |
| Scotland_and_Ireland_Viking_filtered.SG | Scotland_Viking.SG,Ireland_Viking.SG |

**Supplementary Table 15.** Ancient sources used in qpAdm evaluation.

*Results*

We see broadly consistent results when comparing qpAdm and $D_M$, with evidence of increased selectivity and power to distinguish ancestries for $D_M$. As expected, all analyses highlight Baltic- and Slavic-associated sources as explaining most ancestry, with especially good concordance in single-source cases (Fig. S29). For instance, when examining the 10 Vilnius individuals that were only statistically consistent with Baltic ancestry by $D_M$, the only passing present-day qpAdm models are single-source Latvian, and all passing ancient qpAdm models include Lithuania_IA (for 7/10 it is the only source). Similarly, for the 11 individuals who were only statistically consistent with East Europe by $D_M$, 9 have present-day qpAdm models of single-source Ukrainian, and all individuals have a passing ancient qpAdm model including Poland_Slavic_IA (for 10/11 it is the only source).

Some details are consistent across analyses. For example, the only individual statistically consistent with Uralic by $D_M$, LIT454, has a single passing model of Poland_Slavic_IA plus Russia_BolshoyOleniyOstrov_MBA, which is notable given that the latter was included to represent Uralic-associated ancestry.

However, the ancestry in the Vilnius assemblage appears clinal on PCA without clearly delineated clusters. Accordingly, many individuals are in at least one analysis assigned a mix of Baltic and Slavic ancestry or are consistent with both ancestries.

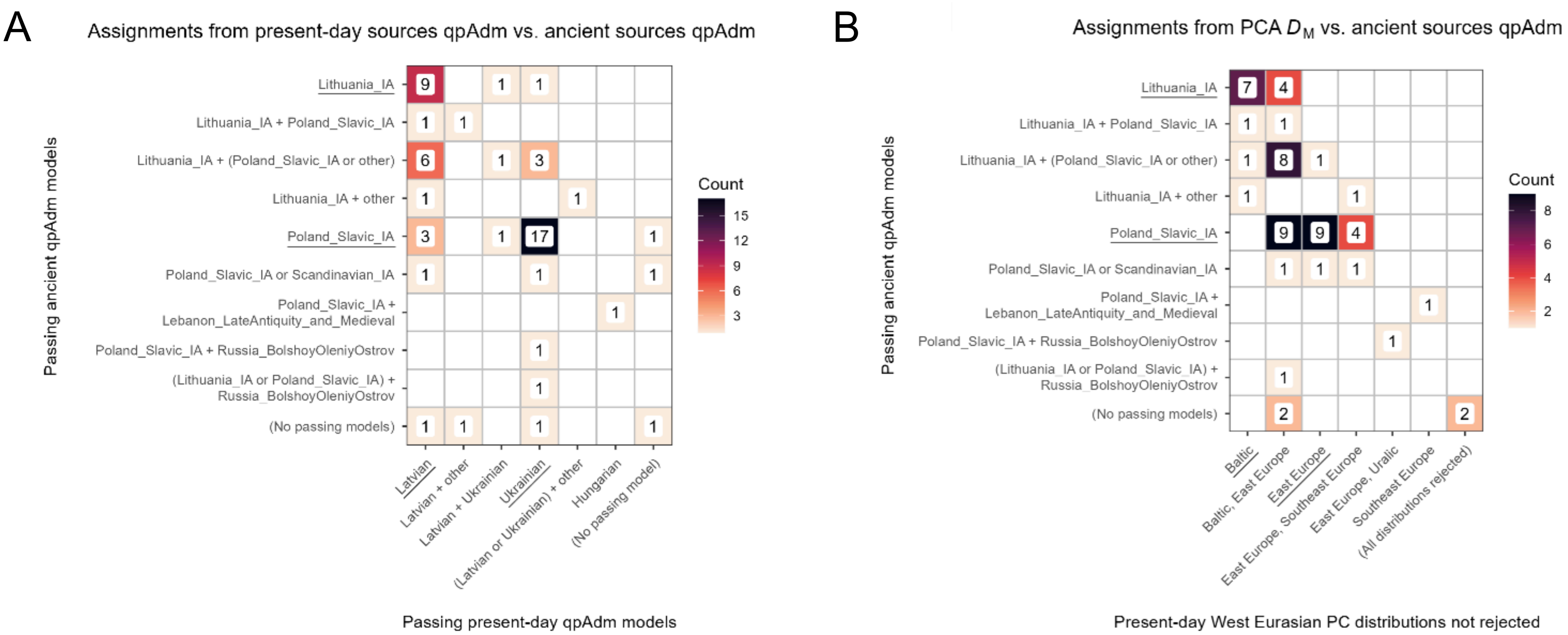


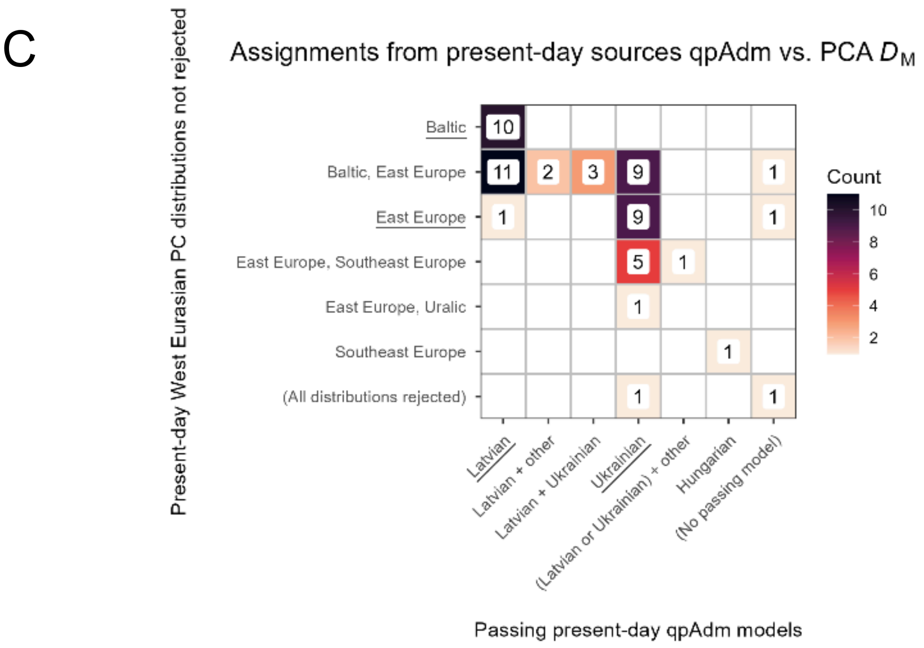


**Supplementary Figure 29:** Concordance of ancestry assessments for Vilnius individuals by method: Mahalanobis distances ($D_M$) on present-day West Eurasian PCA, qpAdm with present-day sources, and qpAdm with ancient sources. Panels cover all pairwise comparisons between the methods. Axis labels indicate qpAdm source populations or consistent PC distributions by $D_M$. Underlined axis labels represent Baltic- or Slavic-associated single-source (qpAdm) or exclusive distribution ($D_M$) assessments. For qpAdm axes, labels with "+", like "Latvian + Ukrainian", denote models with more than one source. Labels with "or", like "Poland_Slavic_IA or Scandinavian_IA", denote cases in which multiple models passed. For $D_M$ axes, labels with commas, like "Baltic, East Europe", denote multiple consistent distributions. Counts of individuals with the corresponding combination are shown by numeric labels and background fill colour. **a)** qpAdm with present-day sources vs. qpAdm with ancient sources. **b)** West Eurasian PCA $D_M$ vs. qpAdm with ancient sources. **c)** qpAdm with present-day sources vs. West Eurasian PCA $D_M$.

We also examined IBD sharing with present-day UK Biobank individuals. We restricted to Vilnius individuals consistent with a single ancestry source: only one statistically consistent PC distribution by $D_M$, or only one passing single-source model in qpAdm. Across methods, we find that individuals assigned Baltic- versus Slavic-associated ancestry are significantly differentiated by their IBD sharing with the most related present-day populations (Supplementary Fig. 30). As previously mentioned, Slavic- and Baltic-assigned groups both show high IBD sharing with present-day Baltic populations. However, IBD sharing with the "Czechia, Hungary, Slovakia" group is elevated for the Slavic-assigned individuals, helping to distinguish these groups.

There are fundamental differences between the PCA-based $D_M$ method and qpAdm, as qpAdm formally models ancestry proportions while the $D_M$ method assesses statistical consistency with reference distributions. Accordingly, we compare results across methods while recognising that they do not represent directly equivalent assignments. When restricting to individuals consistent with a single distribution or ancestry source, our PCA-based assessments appear to be more selective. Only 21 individuals are consistent with a single PC distribution, compared to the 47 single-source models identified in the present-day sources qpAdm analysis and the 33 single-source models in the ancient source qpAdm analysis. This selectivity appears to be meaningfully related to ancestry, as IBD sharing differences between Baltic- and Slavic-assigned individuals are larger and more significant for 3/5 of the reference groups when single-source ancients are defined using $D_M$ compared to the two qpAdm setups. Some qpAdm results reflect low power of the tested model setups to distinguish similar ancestries. For instance, several individuals' only passing models require a small proportion of ancestry from any one of a range of possible sources ("+ other" in Supplementary Fig. 29). Other individuals have passing models that contradict all other ancestry assessments (e.g. single-source Scandinavian_IA). The qpAdm setups also produce a slightly higher count of individuals rejecting all possible models than the count of individuals consistent with no distributions by the $D_M$ method, which may reflect qpAdm's sensitivity to details in model specification, systematic data generation differences between target and source populations, and population histories involving continuous migration[20].

To conclude, $D_M$ and the two qpAdm setups converge on similar ancestry assessments, which are historically sensible and concordant with patterns of IBD sharing. However, these methods may perform less well for ancestries that are closely related or for admixed individuals. Additionally, our PCA-based $D_M$ method, while not a formal ancestry model, produces assessments that are more selective than the qpAdm setups we explored. One likely reason is that the PCA we use includes a large number of reference individuals, providing more power to distinguish related ancestries.

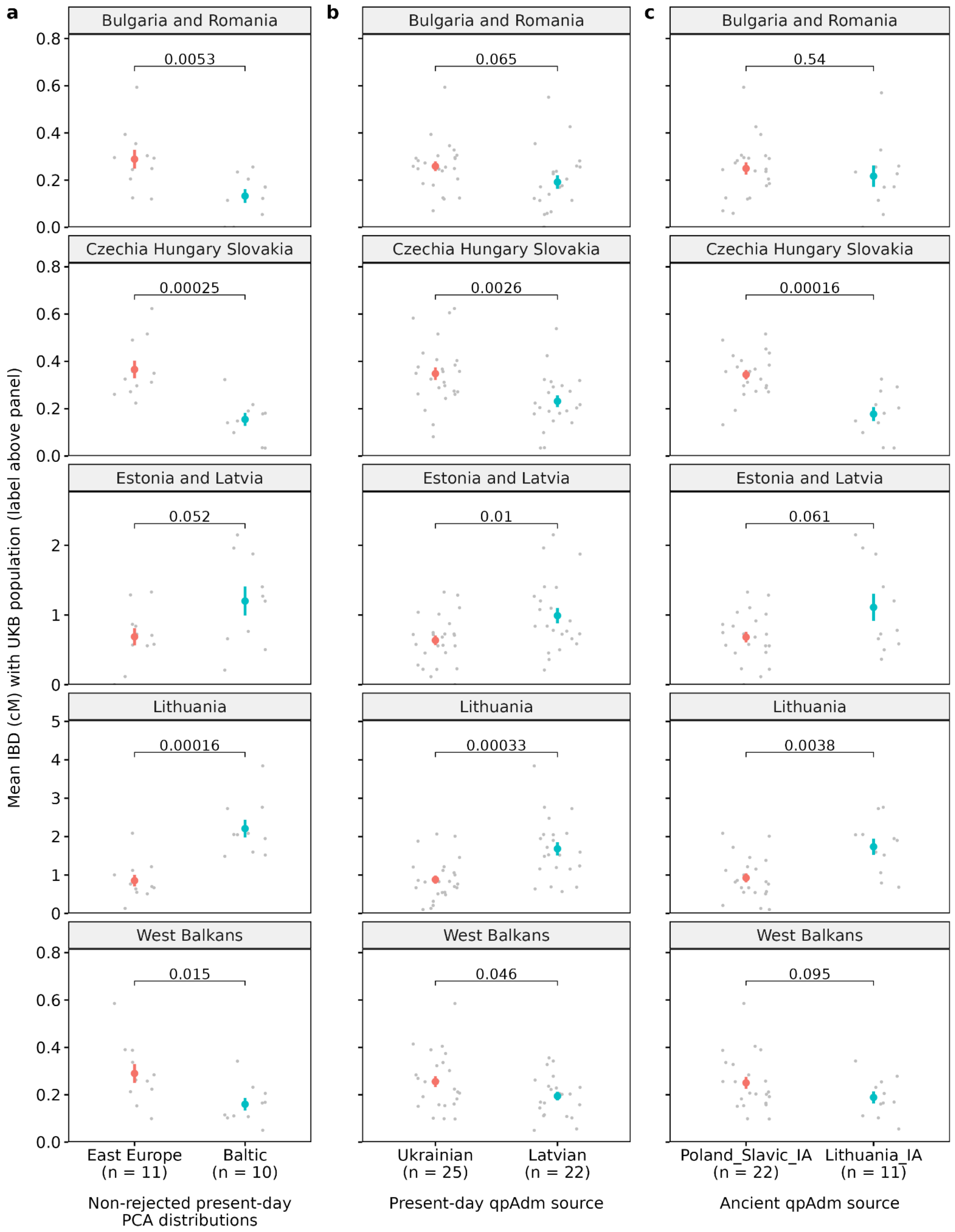


**Supplementary Figure 30:** IBD sharing between populations in UK Biobank and ancient Vilnius individuals assigned to or consistent with a single population by the three methods (PCA based $D_M$, qpAdm with present-day sources, qpAdm with ancient sources). Grey points represent ancient individuals and coloured points with bars show mean ± 2 standard errors. Text labels over brackets indicate *t*-test *P*-value. “West Balkans”: Serbia/Montenegro, Bosnia_and_Herzegovina, Croatia, Slovenia. **a)** Classification based on West Eurasian PCA $D_M$. **b)** Classification based on qpAdm with present-day sources. **c)** Classification based on qpAdm with ancient sources.

#### Supplementary Note 5: Characterisation of diversity by site and period

We identified a high proportion of outliers in Lund, with lower proportions in Trondheim and Vilnius. From visual inspection of Figure 1B, it is clear that the kinds of ancestry observed in the outliers differs by site and period. Below, we use PCA-based Mahalanobis distances ($D_M$) on the West Eurasian PCA described in Supplementary Note 4 to characterise likely sources of outlier ancestry. As demonstrated, this method accords well with qpAdm and IBD sharing. We also refer to IBD sharing rates (Supplementary Fig. 31) between the ancient individuals and our two present-day reference datasets (UK Biobank and OmniExpress European). The results from this section are summarised in the main text ("Assessing the origin of outliers and non-locals").

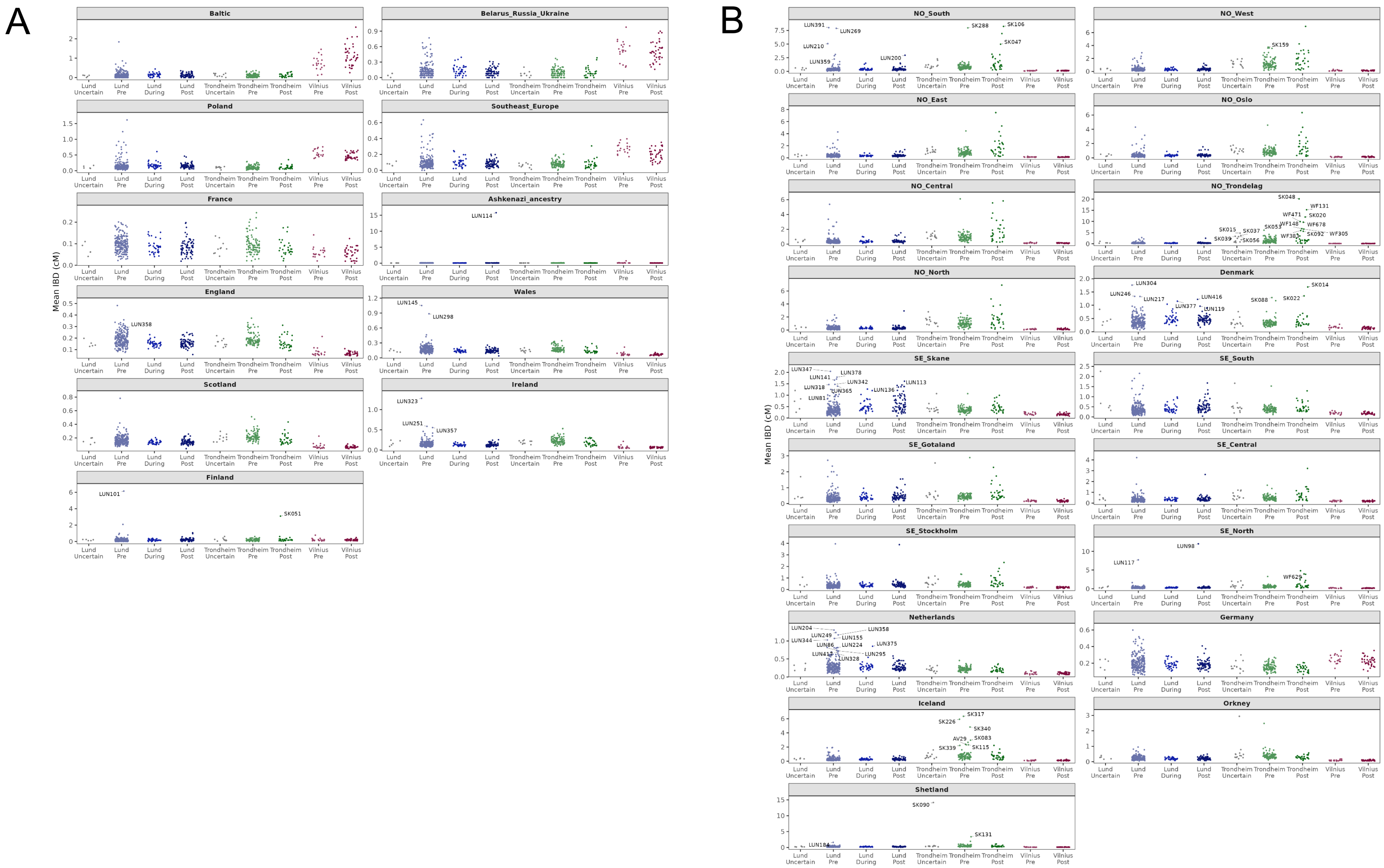


**Supplementary Figure 31:** Average IBD sharing in cM (*y*-axis) between present-day populations (panel titles) and ancient samples split by site and period (*x*-axis). Individuals who share significantly more with one population than all others at $P$ > (0.05 / number of comparisons in figure) according to Welch's *t*-test are labelled. **a)** Present-day populations from the UK Biobank reference set. See Supplementary Table 5 for definition of groups. **b)** Present-day populations from the European OmniExpress reference set.

*Lund*

Pre-BD Lund individuals are notably diverse, and on Figure 1B stretch from present-day French individuals to (northern) Uralic speakers. This spread is also reflected in the $D_M$-based classification using PCs (Supplementary Fig. S32A): 63 individuals are statistically consistent with the distribution for present-day France, and for 9 individuals it is the only reference population they are consistent with; at the other extreme, one individual is consistent with only the Uralic reference population. The low count of exclusive assignments (or consistent statuses) for France likely represents substantially overlapping distributions between France, Britain & Ireland, and Germany & Benelux. A high proportion of individuals (124/218) are consistent with Germany and Benelux, and for 11 individuals it is the sole distribution they are consistent with. We note that Lund was part of Denmark from its founding until 1658, and that Danish ancestry is hard to distinguish from (north) German ancestry (Supplementary Note 3), which may partly explain the large number of individuals consistent with Germany and Benelux. However, many individuals show significantly higher IBD sharing with present-day Dutch and German populations over Denmark (Fig. 3A, Supplementary Figure 31), suggesting actual origins in that region in some cases. 67 individuals are consistent with Britain and Ireland by $D_M$, and for 3 it is the only distribution they are consistent with. Further, 21 individuals are consistent with Baltic or East Europe. Many of these show elevated IBD sharing with Polish and Southeast Europeans (Fig. S31B), with some (e.g. LUN293, LUN88, LUN307) displaying more sharing than all ancient individuals from Vilnius. This suggests a form of Slavic-associated ancestry in pre-BD Lund that is not well represented in our Vilnius set. One candidate is Slavic speakers on the Baltic coast of Germany and Poland.

It is notable that of the 218 pre-BD Lund individuals, only 99 are consistent with Scandinavia by $D_M$. 21 of these are in fact inconsistent with all present-day distributions of West Eurasian populations in the PCA when using the $P>0.01$ threshold, which falls to 12 when $P \geq 0.001$ is used (Fig. 1). Some of these individuals may represent admixed ancestry that is not well approximated by any present-day population (Fig. 1B, Fig 4A), while others appear similar to Iron Age Scandinavians before the occurrence of Viking-related admixture, falling “beyond” present-day Scandinavia, as described in a previous study[28].

During-BD individuals appear much less diverse than the pre-BD set, concordant with the large reduction in the proportion of outliers from pre- to during BD (from 0.52 to 0.15, $P=1.84\times10^{-5}$) mentioned in the main text. A majority of during-BD individuals (31/39) are consistent with Scandinavia by $D_M$ at $P \geq 0.01$, and the remaining outliers are consistent with a narrower spread of geographic regions than the pre-BD outliers. Only one individual is consistent with France, and only two with Britain and Ireland. We observe a clear, Bonferroni-significant decline from pre-BD to during BD in long IBD sharing against all British-Irish groups analysed (Fig. S32B, *t*-test $P$ between $5.7\times10^{-5}$ and $1.3\times10^{-8}$). This decline (and others mentioned in this section) is especially notable given that the sharing of long IBD fragments is expected to be greater in later samples, as fewer meioses separate them from the present-day references. Unlike the pre-BD set, no during-BD individuals are consistent with East Europe or Baltic. Additionally, no clear non-Scandinavian outliers are observed in the IBD results, which might reflect a lower proportion of non-Scandinavian migrants or the presence of ancestry that does not provide strong IBD signals with the reference data.

Post-BD individuals in Lund are less diverse still, with 68/75 individuals consistent with Scandinavia by $D_M$. Interestingly, zero post-BD individuals are consistent with France compared to 63/218 pre-BD individuals. It is also notable that zero post-BD individuals are inconsistent with all present-day PCA distributions, compared to 21/218 for the pre-BD set ($P$=0.011 for pre-BD versus post-BD proportion by chi-squared test). This may indicate declining rates of non-Scandinavian admixture in the preceding few generations for these individuals and the stabilisation of European gene pools similar to those seen today. Of the 7 individuals who are inconsistent with Scandinavia, 6 are consistent with Germany and Benelux. The remaining outlier individual, LUN114, carries Ashkenazi-related ancestry (Supplementary Note 7).

The trend that we observe for Lund is one of decreasing diversity over time, especially from pre-BD to during BD. Pre-BD individuals show a wide spread of ancestry consistent with origins around the North and Baltic Seas, reflecting known movements associated with the Viking Age. Ancestry similar to present-day British and Irish – from the British-Irish Islands themselves or, plausibly, the north coast of continental Europe – appears to have been an important component of pre-BD Lund. The fact that the reduction in diversity is already seen in during-BD samples suggests that the Black Death was not the only driver of this increased homogeneity.

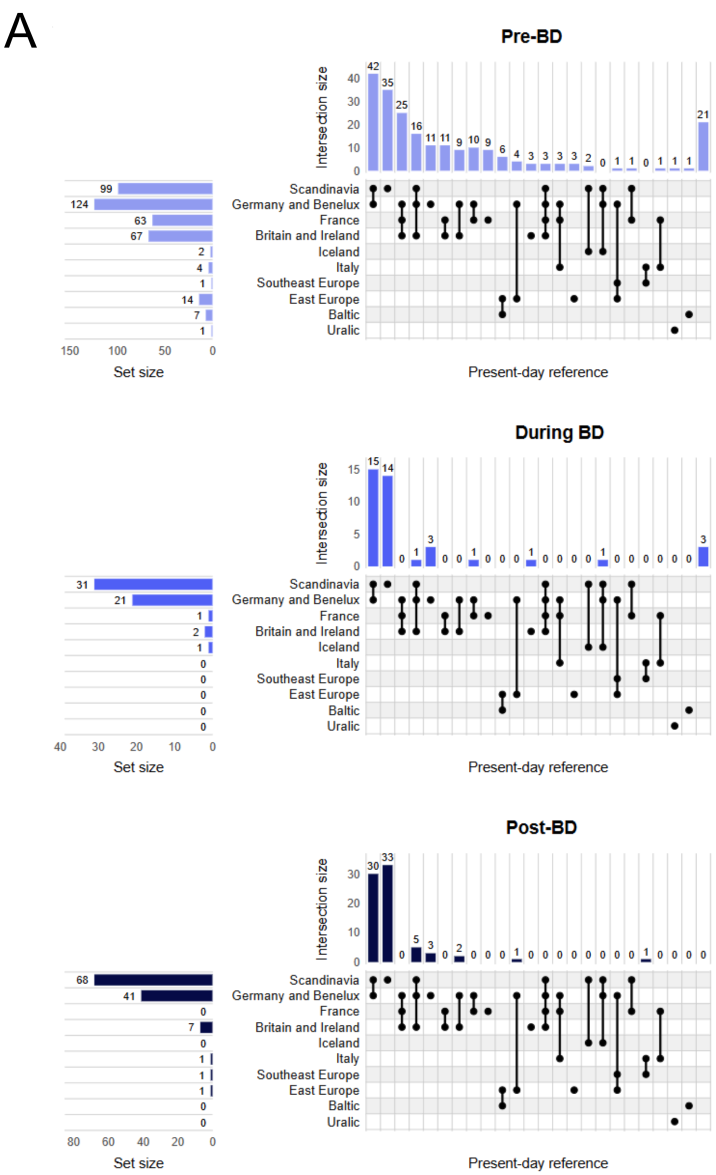



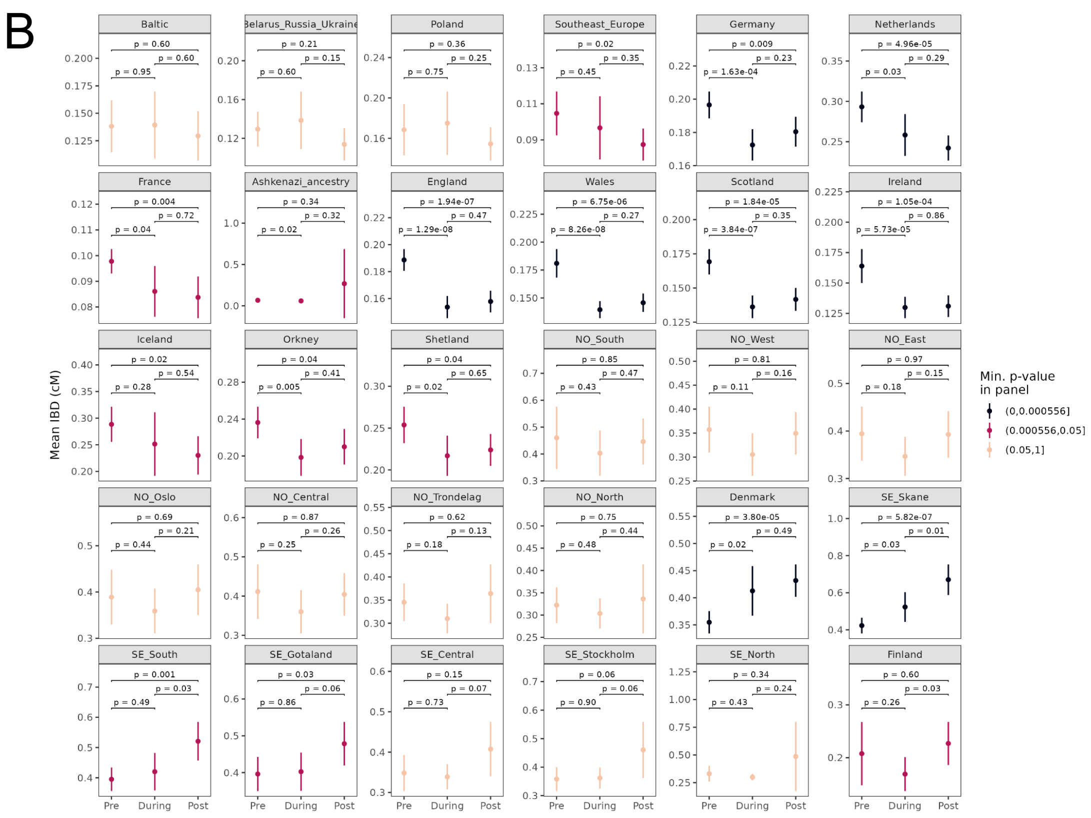



**Supplementary Figure 32:** Ancestry characterisation for Lund. **a)** UpSet plot summarizing regional assignments of ancient individuals according to PCA-based Mahalanobis distances ($D_M$). Filled-in circles in a column mark a combination of non-excluded populations, with vertical bars above indicating the count of individuals showing that combination ("intersection size"). Blank rightmost column indicates individuals with no non-excluded references. Horizontal bars to left indicate count of individuals that include the row's population among their non-excluded populations ("set size"). **b)** Comparison of mean IBD sharing (*y*-axis) by period (*x*-axis) and reference population (panel title). Mean value ± 2 standard errors given by points and vertical bars. Horizontal lines spanning *x*-axis and text labels indicate *p*-value for *t*-tests of IBD sharing values between periods. Points and bars are coloured in black if any *p*-value in the panel passes Bonferroni correction for number of comparisons at α = 0.05, burgundy if below 0.05 (marginal significance), and peach otherwise.

*Trondheim*

Consistent with our previous smaller-scale study of Trondheim[29], we find that pre-BD individuals are mostly explained by Scandinavian-related ancestry and another source related to populations in Britain and Ireland. 16 are consistent with Britain and Ireland by $D_M$, and for 3 individuals it is the only distribution they are consistent with. Notably, 22/95 individuals are not consistent with any present-day distribution using the $P \geq 0.01$ threshold. Many of these individuals (Fig. 4A) fall between Scandinavia and the British-Irish Isles on PCA, suggesting that they carry levels of admixture between these two sources not seen for any present-day distribution. This is also supported by Supplementary Figure 16 and our previous study. That the ancestry profile of pre-BD Trondheim may have been slightly shifted towards British-Irish ancestry compared to present-day Trøndelag is one justification for the more conservative threshold of P=0.001 in our outlier detection procedure; of the 22 individuals who are inconsistent with the Scandinavian distribution at $P<0.01$, 15 are accepted using the lower threshold of P≥0.001. This admixture between a Scandinavian- and British-Irish-associated source may also explain the large number of individuals who are consistent with the distribution for Iceland, whose inhabitants were formed by a similar admixture process, though many pre-BD individuals do in fact carry ancestry distinctly related to present-day Icelanders (Results). A similar admixture also characterises present-day populations in Orkney and Shetland, and several pre-BD individuals show elevated IBD sharing with these populations (Fig. S31, main text). However, Norwegian Iron Age and British-Irish ancestry does not explain all the ancestry observed: for instance, SK271 is consistent only with Germany and Benelux by $D_M$, and SK088 shows distinctly elevated IBD sharing with present-day Denmark.

Post-BD individuals show less British-Irish ancestry. None are consistent Britain and Ireland by $D_M$ (Supplementary Fig. 33A), and we find significant declines in IBD sharing with present-day Wales, Orkney, and Ireland from pre- to post-BD ($P$-values between $1.1 \times 10^{-4}$ and $8.2 \times 10^{-4}$, t-test). Post-BD individuals are more contained within the distribution of present-day Scandinavians (Fig. 4A), with 27/30 individuals inconsistent with Scandinavia by $D_M$.

As with Lund, we see lower diversity in the post-BD samples than in the pre-BD samples. However, the diversity in pre-BD Trondheim is lower than for pre-BD Lund, and appears to be mostly explained by a single cline of Scandinavian to British-Irish ancestry.

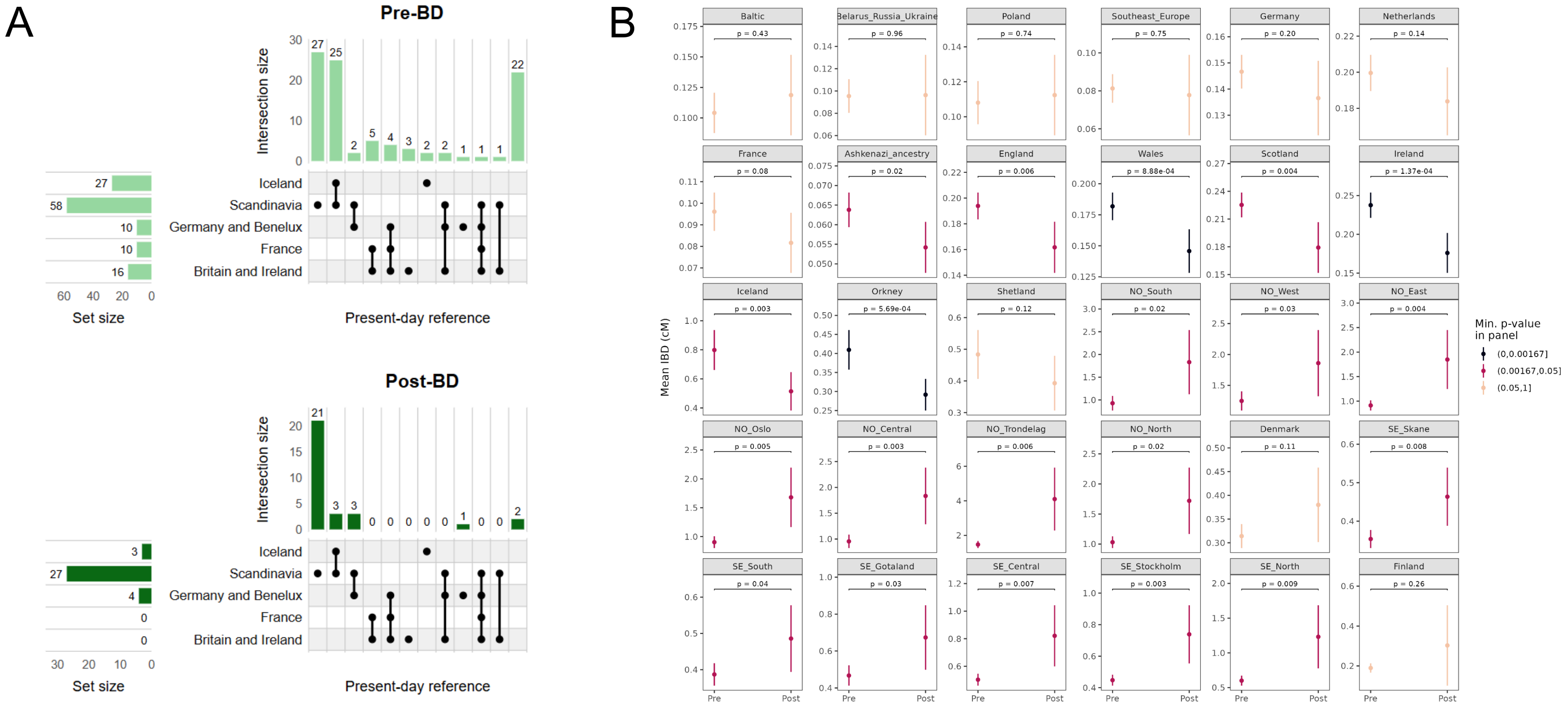


**Supplementary Figure 33:** Ancestry characterisation for Trondheim. **A)** UpSet plot summarizing regional assignments of ancient individuals according to PCA-based Mahalanobis distances ($D_M$). Filled-in circles in a column mark a combination of non-excluded populations, with vertical bars above indicating the count of individuals showing that combination ("intersection size"). Blank rightmost column indicates individuals with no non-excluded references. Horizontal bars to left indicate the count of individuals that include the row's population among their non-excluded populations ("set size"). **B)** Comparison of mean IBD sharing (*y*-axis) by period (*x*-axis) and reference population (panel title). Mean value ± 2 standard errors given by points and vertical bars. Horizontal lines spanning *x*-axis and text labels indicate *p*-value for *t*-tests of IBD sharing values between periods. Points and bars are coloured in black if any *p*-value in the panel passes Bonferroni correction for number of comparisons at α = 0.05, burgundy if below 0.05 (marginal significance), and peach otherwise.

*Vilnius*

The ancestry of the Vilnius individuals is described in more detail in Supplementary Note 4. 17 out of 19 pre-BD individuals are consistent with East Europe by $D_M$, and for 5 it is the only distribution they are consistent with. The East Europe group covers Slavic-speaking countries stretching from Czechia to Russia and is distinct from the Baltic grouping of Lithuania, Latvia, and Estonia. Strikingly, no pre-BD individuals are uniquely consistent with the Baltic distribution by $D_M$. We also observe two strong outliers in this period, LIT448 and LIT452 (Supplementary Note 6).

Post-BD individuals are excavated from two sampling sites. The first is a 15th-16th century mass burial outside the city walls featuring individuals positive for *Y. pestis*, which presumably includes victims of a post-BD wave of the second plague pandemic (Aguonų str.10). The other is a 17th-century Orthodox churchyard (Subačiaus str.7). When the sites are combined (Supplementary Fig. 34), we find a shift towards Baltic ancestry compared to pre-BD according to the $D_M$ method ($\chi^2$test $P$=3.4x$10^{-4}$ for proportion of individuals consistent with Baltic), which is supported by a significant increase in IBD sharing with Baltic and decrease in IBD sharing with Southeast European reference populations (both $P$<0.001 by Welch's *t*-test). We observed notable differences between the two post-BD sites (Supplementary Fig. 35). Baltic ancestry is most concentrated in the earlier plague-associated mass burial: this site features all six post-BD individuals who exclusively are consistent with Baltic by $D_M$, and also shows significantly more IBD sharing with present-day Baltic compared to the pre-BD set and the post-BD Orthodox churchyard (both $P$<2.6x$10^{-4}$ by Welch's *t*-test). The post-BD Orthodox churchyard appears more similar in ancestry to the pre-BD set. However, a higher proportion of pre-BD individuals are assigned as outliers in the outlier detection procedure (Fig. 3C, Methods) (9/19 vs. 1/19 for Post-BD Orthodox, chi-squared $P$=0.0099), and the pre-BD set shows marginally higher IBD sharing with Poland, Southeast Europe, and Germany (all $P$~0.01), suggesting some differences in ancestry composition. The Post-BD Orthodox site also shows an increase in individuals consistent with Baltic by $D_M$ compared to the pre-BD set ($P$=0.021, $\chi^2$test).

As in Lund and Trondheim, pre-BD individuals from Vilnius show greater diversity of ancestry than post-BD individuals. We also observe clear differences between the three sampling sites in Vilnius, with an apparent pulse of Baltic ancestry in the 15th-16th century non-churchyard site and more Slavic-associated ancestry in the earlier and later Orthodox churchyards. Because Orthodoxy spread to Lithuania partly through conquest of, and migration from, the Slavic East, it is possible that the Orthodox cemeteries are biased in favour of Slavic-associated ancestry compared to the contemporary demographics of Vilnius (see Discussion).

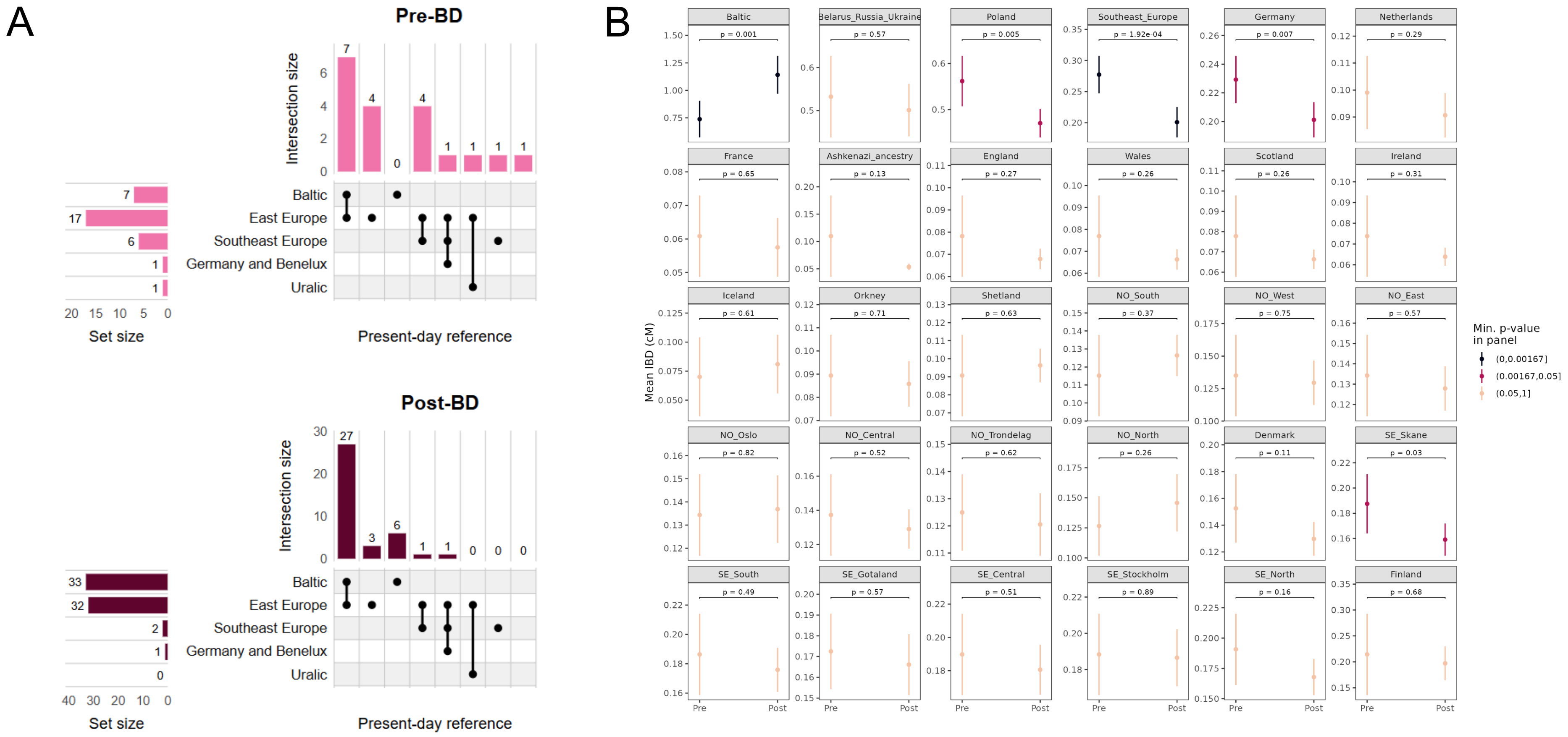


**Supplementary Figure 34:** Ancestry characterisation for Vilnius. **A)** UpSet plot summarizing regional assignments of ancient individuals according to PCA-based Mahalanobis distances ($D_M$). Filled-in circles in a column mark a combination of non-excluded populations, with vertical bars above indicating the count of individuals showing that combination ("intersection size"). Blank rightmost column indicates individuals with no non-excluded references. Horizontal bars to left indicate count of individuals that include the row's population among their non-excluded populations ("set size"). Bar colours indicate period. **B)** Comparison of mean IBD sharing (*y*-axis) by period (*x*-axis) and reference population (panel title). Mean value ± 2 standard errors given by points and vertical bars. Horizontal lines spanning *x*-axis and text labels indicate *p*-value for *t*-tests of IBD sharing values between periods. Points and bars are coloured in black if any *p*-value in the panel passes Bonferroni correction for number of comparisons at α = 0.05, burgundy if below 0.05 (marginal significance), and peach otherwise.

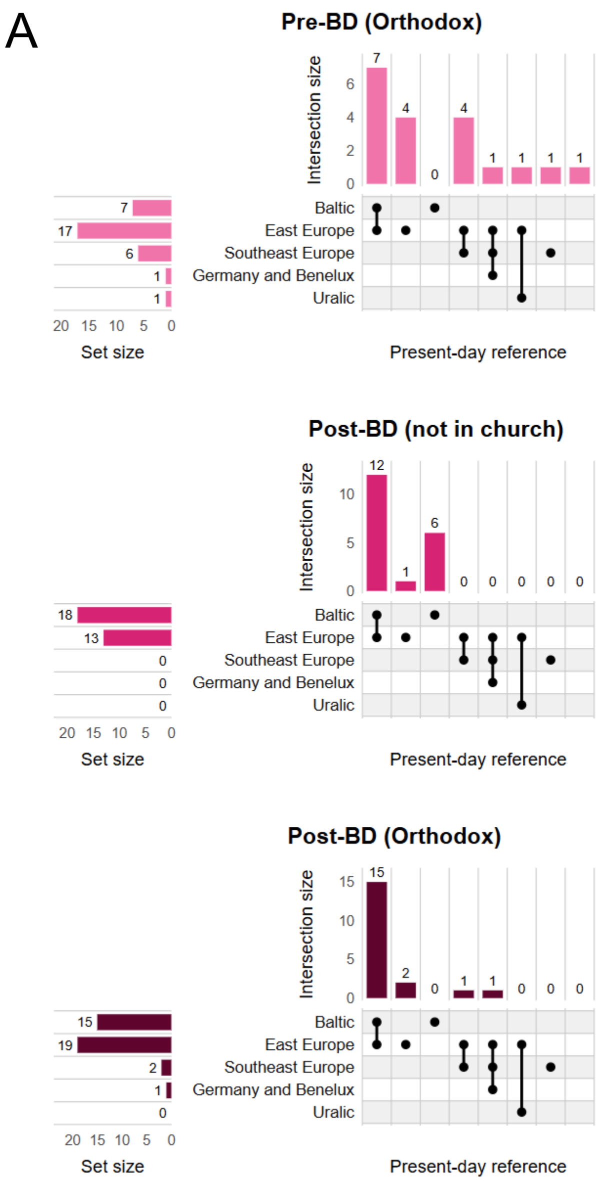


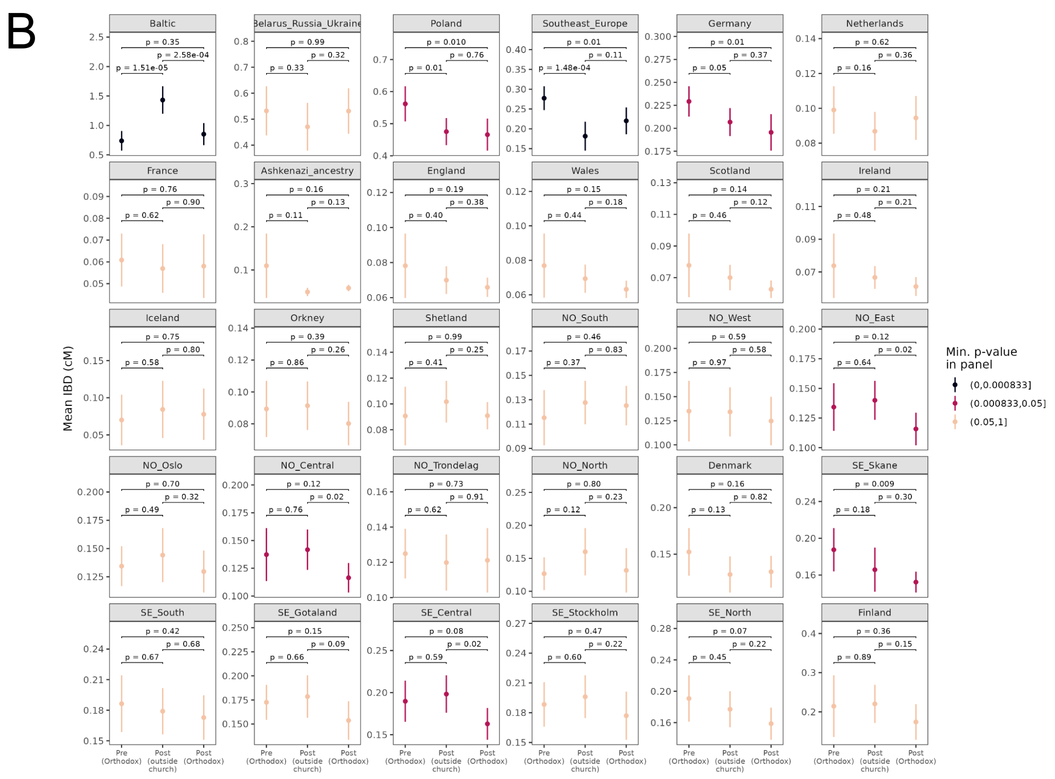


**Supplementary Figure 35:** Ancestry characterisation for Vilnius with 3-way site split: Pre-BD (Orthodox) for Bokšto str. 6, Post-BD (not in church) for Aguonų str.10, and Post-BD (Orthodox) for Subačiaus str.7. **A)** UpSet plot summarizing regional assignments of ancient individuals according to PCA-based Mahalanobis distances ($D_M$). Filled-in circles in a column mark a combination of non-excluded populations, with vertical bars above indicating the count of individuals showing that combination (“intersection size”). Blank rightmost column indicates individuals with no non-excluded references. Horizontal bars to left indicate count of individuals that include the row’s population among their non-excluded populations (“set size”). **B)** Comparison of mean IBD sharing (*y*-axis) by period (*x*-axis) and reference population (panel title). Mean value ± 2 standard errors given by points and vertical bars. Horizontal lines spanning *x*-axis and text labels indicate *p*-value for *t*-tests of IBD sharing values between periods. Points and bars are coloured in black if any *p*-value in the panel passes Bonferroni correction for number of comparisons at α = 0.05, burgundy if below 0.05 (marginal significance), and peach otherwise.

#### Supplementary Note 6: Ancestry outliers in Vilnius

Two individuals from the pre-BD assemblage at Vilnius carry ancestry that diverges substantially from the Baltic-Slavic majority ancestry, providing evidence for long-distance mobility into the emerging capital during the early-middle 14th century.

##### *LIT452*

LIT452 projects close to present-day Southeast Europeans on the West Eurasian PCA (Fig. 1B), though it is inconsistent with all present-day reference populations at $P$<0.01 by $D_M$. In a continent-level supervised $K$=5 ADMIXTURE analysis using the 1000 Genomes dataset, 8.7% of this individual's ancestry is assigned to the South/West Asian component and the remainder to European (Fig. S25). This moderately elevated assignment of South/West Asian ancestry is the second-highest in the Vilnius assemblage aside from the other outlier named in this note, LIT448. The sample's chrY haplogroup is J2a1a1a2a2b2a (J-PH4000), which according to databases of present-day individuals (YFull, FamilyTreeDNA) is only attested in Qatar, Bahrain, and Kuwait. Its mtDNA haplogroup U5b1b1a is broadly distributed across northern Eurasia and reaches its highest frequencies in northern/northeastern Europe, which might suggest a local maternal origin. The IBD sharing profile against modern individuals (Fig. S20) is not strikingly different from the other individuals in the assembly. This could be explained by the individual having ancestry from one source similar to other Vilnius samples and another source with low affinity to the present-day European references we use for IBD analysis.

The ancestry profile of LIT452 appears similar to contemporary southeast Europeans[22,30]. By the eighth century, a Slavic-associated population had entered the Balkan Peninsula and began to mix with an existing ancestry cline including individuals with Near Eastern ancestry. Byzantine-era samples from Montenegro whose ancestry was shaped by this process[23] display up to 4.5% West/South Asian ancestry in our K=5 ADMIXTURE analysis. While this is lower than the proportion we observe for LIT452, recently presented samples from Croatia[22] dating to roughly the same time as LIT452 feature individuals assigned high proportions of similar ancestries (up to 40% “West Asian” and 21% “Near Eastern” ancestry in their models). In our IBD analyses against UK Biobank, LIT452 does not show a distinctly higher affinity to southeast Europe than other Vilnius samples (Fig. S20). This might be due to low sample sizes of Southeast Europeans in UK Biobank or high haplotype diversity in present-day Southeast Europeans. In our evaluation of ancestry assignment methods in Supplementary Note 4, LIT452 has a passing qpAdm model involving Poland_Slavic_IA and Lebanon_LateAntiquity_and_Medieval.SG, and when using present-day sources has a passing single-source model for Hungarian.

To conclude, while LIT452’s signs of Slavic/Baltic and Near Eastern ancestry might indicate recent admixture, another, perhaps more parsimonious, hypothesis is that they belonged to a contemporary Southeast European population deriving from older admixture between a

Slavic/Baltic-associated source and another source with moderate levels of Near Eastern ancestry.

##### *LIT448*

The other individual, LIT448, appears to be pulled towards the Caucasus on the first two principal components of the West Eurasian reference panel (Fig. 1B) and does not fit any modern population at $P$>0.01 according to $D_M$. It is a strong ancestry outlier compared to the other samples in the set, being assigned 10.8% East Asian ancestry and 10.2% South/West Asian ancestry in the continental supervised ADMIXTURE analysis (Fig. S25). According to YFull and FamilyTreeDNA, its chrY haplogroup G2a2b2a1a1a1b1a2 (G-Y32924 / G-Y32922) is found among North Caucasus populations (Adyghe, Abazin, Chechen) and Russian Tatars, with scattered examples in populations historically affected by Caucasus-derived population movements (Bulgaria, Romania, Turkey). Its mtDNA haplogroup C4a1a1 is rare but broadly distributed across Poland, Russia, Finland, and China.

This individual was sequenced at low depth (0.11x). While this should still allow accurate inferences from analyses based on allele frequencies like PCA and ADMIXTURE, especially after imputation, IBD calling at this depth is expected to be less robust. With this caveat in mind, we note that LIT448 shows the highest IBD sharing with the UKB Baltic group, but its IBD affinities with all European populations are generally lower than the other Vilnius samples (Fig. S20).

The combination of admixed autosomal ancestry from European, East Asian, and West/South Asian sources, a Caucasus-associated chrY lineage, and an mtDNA haplogroup with northern/eastern Eurasian affinity is strongly reminiscent of contemporary Eurasian steppe populations. These diverse groups derived from a complex east-west mixture of Turkic, Mongolic, Iranic, and Caucasus-associated sources[31,32]. The Lipka Tatars, who are first attested in Vilnius at the end of the 14th century and are still present in Lithuania today, represent a similar mix of ancestries[33]. It is perhaps notable that the cited paper provides G2a2b2a1a1a1, slightly upstream of LIT448's assignment, as an example of a Lipka Tatar-associated chrY haplogroup. However, early Tatar migrants to Lithuania are generally held to have been Tengrian pagans or Muslims, which contrasts with LIT448 interment in an Orthodox Christian churchyard.

There are plausible historical explanations for the presence of ancestry outliers in Vilnius at this time. Notably, the first written record of Vilnius is in the 1323-1324 letters from Grand Duke of Lithuania Gediminas[12], which were sent across Europe and explicitly invited merchants and craftsmen to settle in the city. The Golden Horde, a Mongol Empire successor state, brought Eurasian steppe populations close to the borders of the Grand Duchy of Lithuania. The Lipka Tatars are one example of a group carrying Eurasian steppe ancestry whose ethnogenesis traces to emigration from the Golden Horde to Lithuania. While the settlement of Lipka Tatars in Lithuania is traditionally credited to Vytautas the Great (reigned 1392-1430)[34], LIT448 strongly suggests that individuals with recent Eurasian steppe ancestry were present in Vilnius decades before this period.

#### Supplementary Note 7: Ancestry outlier in Lund

LUN114 projects close to modern Greek and Albanian populations on the West Eurasian PCA (Fig 1B), appearing as a prominent outlier from the other Lund samples, which generally project near present-day populations around the North, Irish, and Baltic Seas. According to $D_M$ using 7 PCs, it is marginally consistent with the distribution of modern Italians ($P$ = 0.016). In the continental K=5 supervised ADMIXTURE analysis, LUN114 is assigned the lowest proportion of European autosomal ancestry of all Lund samples (91.7%, Fig. S25), with the remainder assigned to the West/South Asian component.

IBD analysis reveals that LUN114 shares remarkably many >7 cM IBD segments with UK Biobank individuals of Ashkenazi Jewish ancestry (Supplementary Fig. 36). This subset comprises around 5,000 individuals of majority or partial Ashkenazi ancestry and has been identified in previous studies[35,36]. LUN114 shows over 30 times higher average IBD sharing with the Ashkenazi subset than the next-highest UK Biobank grouping, Poland. This intense IBD sharing is consistent with the well-documented founder effect and subsequent endogamy of Ashkenazim[25,37].

LUN114 is assigned the chrY haplogroup E1b1b1b2a1b (E-PF2025/E-PF1975) by HaploGrouper[38], which appears highly enriched in Ashkenazi Jews: as assessed by the same method, the 1% of men in the Ashkenazi ancestry subset of UK Biobank carry 69% of carriers of this haplogroup. Notably, the only AADR v62.0 sample labelled as carrying this haplogroup is a 12th-century individual from England who has strong genetic affinity with present-day Ashkenazim[39]. Haplogroup calling with YLeaf followed by manual inspection of the BAM file further refines the lineage to E-FT158701, with a time to the most recent common ancestor around 1100 years ago according to Family Tree DNA Discover (https://discover.familytreedna.com/y-dna/E-FT158701/story) or 650 years ago according to YFull (https://www.yfull.com/tree/E-FT158701*/).

LUN114's mtDNA haplogroup - H27+T16093C - is broadly distributed across Europe, but is not characterised as a typical Ashkenazi mtDNA haplogroup[40].

Despite clear signals of Ashkenazi ancestry, LUN114’s autosomal profile is not a perfect match to present-day Ashkenazi Jews. On a version of the West Eurasian PCA (Supplementary Fig. 37) including reference Jewish populations from Medieval Norwich[39] and Erfurt[25], as well as modern Jewish samples from Behar et al. (2013)[41] and Gladstein and Hammer (2019)[42], LUN114 projects outside the cluster representing modern and putative ancient Ashkenazi populations and appears to be pulled towards populations in northern and western Europe. While LUN114 is nearest on PCA to medieval Ashkenazi Jews of Central/Eastern European origin (“Erfurt-EU”)[25], this may result from admixture and not from origin in this subgroup.

Another observation that is inconsistent with full Ashkenazi ancestry is the absence of detectable runs of homozygosity (ROH) in LUN114. Due to founder effects and historical endogamy, Ashkenazi individuals display significantly elevated ROH compared to other

European populations, which is already seen in the 14th century Erfurt assemblage and to some extent in 12th-century Norwich. Notably, LUN114 is one of the 8/337 Lund samples for which we identify no ROH fragments, and is the Lund sample with the highest autosomal heterozygosity (Supplementary Table 4). This could indicate mixed ancestry in LUN114 involving an Ashkenazi-related source and a non-Ashkenazi source.

Taken together, the strong IBD affinity with modern Ashkenazi populations, the Ashkenazi-enriched chrY haplogroup, the absence of a typical Ashkenazi mtDNA haplogroup, the absence of ROH, and the high heterozygosity suggest that LUN114 had Ashkenazi ancestry perhaps only via his paternal ancestors.

The identification of an individual with Ashkenazi ancestry in medieval Lund is noteworthy given that the earliest documented Jewish communities in Scandinavia date to the late 17th century[43]. There exist scattered attestations of individuals of Jewish descent in Scandinavian-controlled lands prior to this time, including a doctor employed by Swedish king Gustav Vasa in 1557[44] and a Sephardi merchant named Albert Dionis who was appointed master of the mint in Danish-controlled Glückstadt in 1619[45]. As burial activity at Trinitatis cemetery ceased by 1536 CE, LUN114 may represent the earliest known case of an individual with Ashkenazi ancestry in Scandinavia.

The events which led to an individual with partial Ashkenazi ancestry being interred in a medieval Lund churchyard are unclear. LUN114's strontium isotope ratio of 0.70975 falls just below the 0.710-0.712 reference level for Lund, possibly indicating that they spent their childhood outside the region. The Trinitatis cemetery from which LUN114 was excavated is thought to have served a higher-status congregation[5]. The individual was buried without a coffin on a shroud, consistent with the relatively plain burial practices documented for this period at the site. The burial in a Christian context without apparent distinction from other parishioners suggests integration into Christian communities regardless of the individual's precise origins.

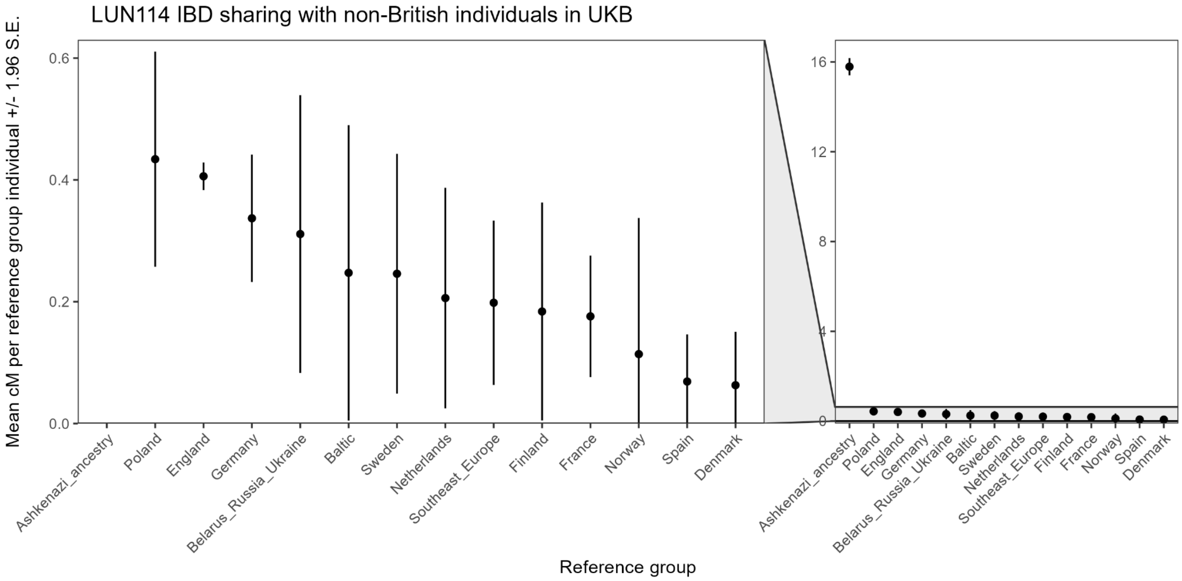


**Supplementary Figure 36.** IBD sharing between LUN114 and non-British groups in UK Biobank. The left panel represents a zoomed-in slice of the right panel (shaded).

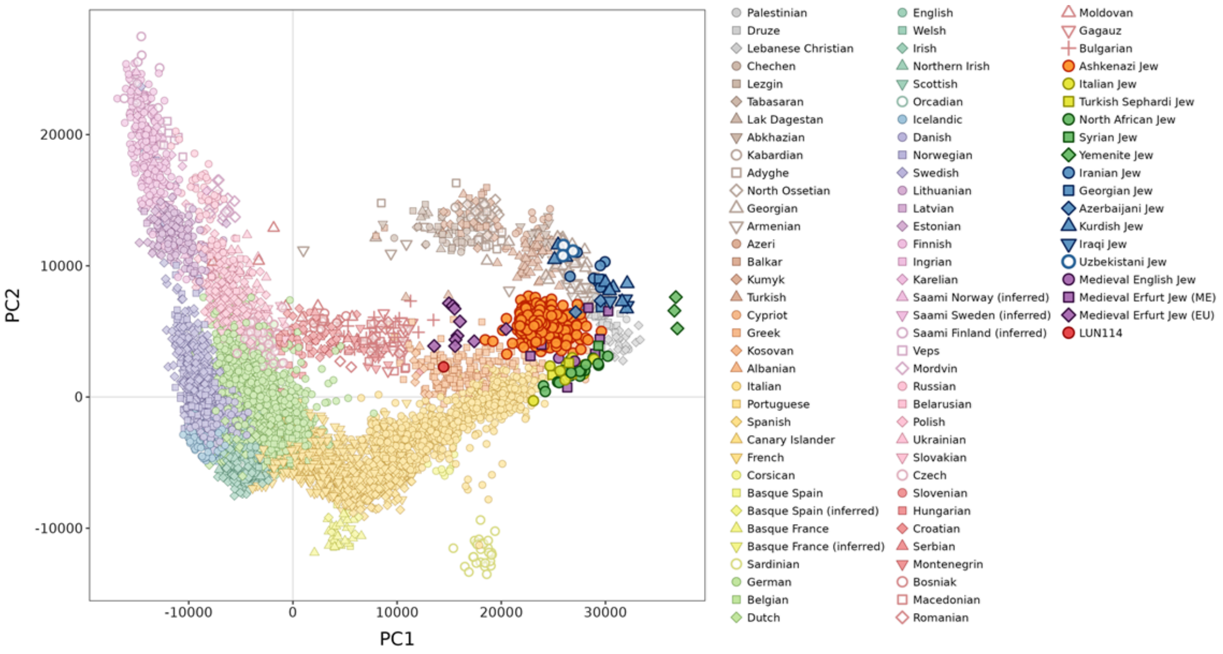


**Supplementary Figure 37.** PCA calculated on the West Eurasian reference set, with the addition of Jewish individuals sampled in Behar et al. 2013 and Gladstein and Hammer 2019. Projected coordinates for ancient Jewish-associated individuals from Brace et al. 2022 (*Medieval English Jew*) and Waldman et al. 2022 (*Erfurt Jew*), as well as LUN114 are also shown.

1. Cinthio, M. & Ödman, A. Vägar mot Lund: en antologi om stadens uppkomst, tidigaste utveckling och entrepenaden bakom de stora stenbyggnaderna. 11–29 (2018).
2. Arcini, C. Health and disease in early Lund, Osteo-pathologic studies of 3,305 individuals buried in the cemetery area of Lund 990-1536. **8**, (1999).
3. Arcini, C. *et al.* Living conditions in times of plague. in *Environment, Society and the Black Death* (ed. Lagerås, P.) 104–140 (Oxbow Books, 2016).
4. Wallin, C. Premonstratenserna och deras kloster i Skåne. *Skånes Hembygdsförbund Årsbok, 1987/88: Skånska kloster* 63–81 (1989).
5. Cinthio, M. De första stadsborna: medeltida gravar och människor i Lund. (2002).
6. Christophersen, A. *Under Trondheim: fortellinger fra bygrunnen*. (Museumsforlaget, 1992).
7. Christophersen, A. E. & Nordeide, S. W. Kaupangen ved Nidelva. Riksantikvarens skrifter nr. 7. (1994).
8. Ramstad, S. Gregoriuskirka": Rekonstruktion og funksjonell analyse. *Norges teknisk-naturvitenskaplige universitet (NTNU)* (2002).
9. Sæhle, I. *et al.* Arkeologiske undersøkelser i Søndre gate 7–11, Peter Egges plass, Krambugata 2–4 m. fl., Trondheim Trøndelag (TA 2016/21, TA 2017/03). Landskapsutvikling, tidlig urban aktivitet og middelaldersk kirkested. *Landskapsutvikling, tidlig urban aktivitet og middelaldersk kirkested. NIKU rapport* **97**, (2021).
10. Reed, I. *Excavations Outside the West Front of Nidaros Cathedral in Trondheim. D. 1*. (NIKU Norwegian Institute for Cultural Heritage Research, Oslo, 1998).
11. Davies, N. *Europe*. (Vintage Digital, London, England, 2010).
12. Ščavinskas, M. Darius baronas, S.c. rowell, the conversion of Lithuania. From pagan barbarians to the late medieval Christians, Vilnius: Institute of Lithuanian literature and folklore, 2015. 627 p. ISBN 978-609-425-152-8. *Lith. Hist. Stud.* **22**, 161–169 (2018).
13. Jonaitis, R. & Kaplūnaitė, I. *Senkapis Vilniuje, Bokšto Gatvėje. XIII - XV . Laidosenos Lietuvoje Bruožai*. (Lietuvos istorijos institutas, Vilnius, 2020).

26. Maier, R. *et al.* On the limits of fitting complex models of population history to -statistics.

*Elife* **12**, (2023).

27. Palamara, P. F., Lencz, T., Darvasi, A. & Pe'er, I. Length distributions of identity by descent reveal fine-scale demographic history. *Am J Hum Genet* **91**, 809–822 (2012).
28. Rodríguez-Varela, R. *et al.* The genetic history of Scandinavia from the Roman Iron Age to the present. *Cell* **186**, 32–46.e19 (2023).
29. Gopalakrishnan, S. *et al.* The population genomic legacy of the second plague pandemic. *Curr Biol* **32**, 4743–4751.e6 (2022).
30. Olalde, I. *et al.* A genetic history of the Balkans from Roman frontier to Slavic migrations. *Cell* **186**, 5472–5485.e9 (2023).
31. Damgaard, P. de B. *et al.* 137 ancient human genomes from across the Eurasian steppes. *Nature* **557**, 369–374 (2018).
32. Saag, L. *et al.* North Pontic crossroads: Mobility in Ukraine from the Bronze Age to the early modern period. *Sci Adv* **11**, eadr0695 (2025).
33. Pankratov, V. *et al.* East Eurasian ancestry in the middle of Europe: genetic footprints of Steppe nomads in the genomes of Belarusian Lipka Tatars. *Sci Rep* **6**, 30197 (2016).
34. Jakulytė-Vasil, M. TATARS' ASSIMILATION/ INTEGRATION IN THE SOCIAL FABRIC OF THE GRAND DUCHY OF LITHUANIA. in *Orient in the Social Tradition of the Grand Duchy of Lithuania: Tatars and Karaims* (ed. Pšibilskis, V. B.) (Vilniaus universiteto leidykla, 2008).
35. Naseri, A. *et al.* Personalized genealogical history of UK individuals inferred from biobank-scale IBD segments. *BMC Biol* **19**, 32 (2021).
36. UK Biobank Whole-Genome Sequencing Consortium. Whole-genome sequencing of 490,640 UK Biobank participants. *Nature* **645**, 692–701 (2025).
37. Carmi, S. *et al.* Sequencing an Ashkenazi reference panel supports population-targeted personal genomics and illuminates Jewish and European origins. *Nat Commun* **5**, 4835 (2014).
38. Jagadeesan, A. *et al.* HaploGrouper: a generalized approach to haplogroup classification.

*Bioinformatics* **37**, 570–572 (2021).

39. Brace, S. *et al.* Genomes from a medieval mass burial show Ashkenazi-associated hereditary diseases pre-date the 12th century. *Curr Biol* **32**, 4350–4359.e6 (2022).
40. Costa, M. D. *et al.* A substantial prehistoric European ancestry amongst Ashkenazi maternal lineages. *Nat Commun* **4**, 2543 (2013).
41. Behar, D. M. *et al.* No evidence from genome-wide data of a Khazar origin for the Ashkenazi Jews. *Hum Biol* **85**, 859–900 (2013).
42. Gladstein, A. L. & Hammer, M. F. Substructured Population Growth in the Ashkenazi Jews Inferred with Approximate Bayesian Computation. *Mol Biol Evol* **36**, 1162–1171 (2019).
43. Lausten, M. S. *Jews and Christians in Denmark: From the Middle Ages to Recent Times, Ca. 1100-1948*. (Brill, 2015). doi:10.1163/9789004304376.
44. Grimberg, C. *440 (Svenska folkets underbara öden / X. Supplement I)*. (1924).
45. Kisch, C. The Jewish community in Denmark : History and present status. *Judaism* **47**, 214–231 (1998).
